# Copulas and their potential for ecology

**DOI:** 10.1101/650838

**Authors:** Shyamolina Ghosh, Lawrence W. Sheppard, Mark T. Holder, Terrance D. Loecke, Philip C. Reid, James D. Bever, Daniel C. Reuman

**Affiliations:** Department of Ecology and Evolutionary Biology and Kansas Biological Survey, University of Kansas, Lawrence, KS, 66045, USA; Department of Ecology and Evolutionary Biology and Biodiversity Institute, University of Kansas, Lawrence, KS, 66045, USA; Environmental Studies Program and Kansas Biological Survey, University of Kansas, Lawrence, KS, 66047, USA; School of Biological & Marine Sciences, University of Plymouth, Drake Circus, Plymouth PL4 8AA, UK; Continuous Plankton Recorder Survey, The Marine Biological Association, The Laboratory, Citadel Hill, Plymouth PL1 2PB, UK; Department of Ecology and Evolutionary Biology and Kansas Biological Survey, University of Kansas, Lawrence, KS, 66047, USA; Laboratory of Populations, Rockefeller University, 1230 York Ave, New York, NY, 10065, USA

**Author notes:** Correspondence:* Daniel Reuman, 2101 Constant Ave, Lawrence, KS, 66047.

**Keywords:** dependence, regression, correlation, copula, population, community, ecosystem functioning

## Abstract

All branches of ecology study relationships among and between environmental and biological variables. However, standard approaches to studying such relationships, based on correlation and regression, provide only a small slice of the complex information contained in the relationships. Other statistical approaches exist that provide a complete description of relationships between variables, based on the concept of the *copula*; they are applied in finance, neuroscience and other fields, but rarely in ecology. We here explore the concepts that underpin copulas and examine the potential for those concepts to improve our understanding of ecology. We find that informative copula structure in dependencies between variables is common across all the environmental, species-trait, phenological, population, community, and ecosystem functioning datasets we considered. Many datasets exhibited asymmetric tail associations, whereby two variables were more strongly related in their left compared to right tails, or *vice versa*. We describe mechanisms by which observed copula structure and asymmetric tail associations can arise in ecological data, including a Moran-like effect whereby dependence structures between environmental variables are inherited by ecological variables; and asymmetric or nonlinear influences of environments on ecological variables, such as under Liebig’s law of the minimum. We also describe consequences of copula structure for ecological phenomena, including impacts on extinction risk, Taylor’s law, and the stability through time of ecosystem services. By documenting the importance of a complete description of dependence between variables, advancing conceptual frameworks, and demonstrating a powerful approach, we aim to encourage widespread use of copulas in ecology, which we believe can benefit the discipline.

## 1 Introduction

All branches of ecology study relationships among biological variables and relationships between environmental and biological variables. However, commonly used correlation and regression approaches to studying such relationships are limited, and provide only a partial description of a relationship between variables. For instance, datasets showing markedly different relationships may have the same correlation coefficient (Fig. 1A-B). The variables of Fig. 1A (respectively, Fig. 1B) are principally related in the left (respectively, right) portions of their distributions, an asymmetric pattern of dependence that can have ecological significance, as discussed below, but that is not captured by correlation. Although correlation is only one way to study relationships between variables, other common metrics also provide only partial information.

**Figure 1:**
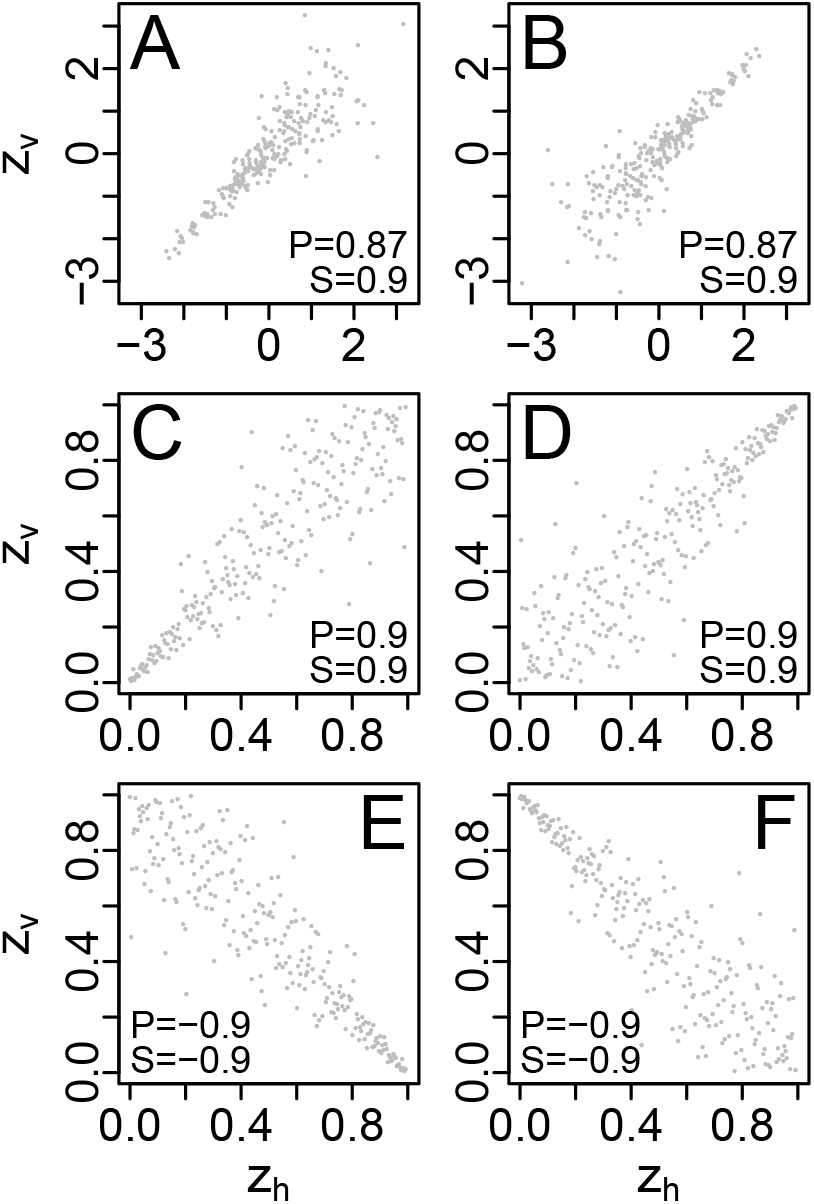
(A-D) Bivariate datasets showing diverse relationships between variables. Some of these datasets nevertheless have the same Pearson (P) or Spearman (S) correlation coefficients. C and D show normalized ranks of A, B, respectively. (E,F) Reflections of C, D, respectively, about a vertical axis. Subscripts h and v in the axis labels stand for “horizontal” and “vertical”.

Well-developed approaches do exist, however, and are applied widely in other fields, that provide a description of the dependence between two or more variables that can be mathematically proven to be complete (Nelsen 2006; Joe 2014; Mai & Scherer 2017). The approaches are based on the concept of the *copula*. Copula approaches separate the information of a bivariate random variable into two distinct parts: the information in the marginal distributions (which says nothing about the dependence between the variables), and the remaining information (which is solely about dependence). We here introduce some simple concepts, based on taking ranks, that relate to copulas and give a conceptual flavor of copula ideas to be introduced more formally in the “Background on copulas” section below (section 2). Given a sample (*x_i_*, *y_i_*), *i* = 1, 2, …, *n* (e.g., Fig. 1A-B), information about the structure of the dependence between *x* and *y* can be separated from information contained in the marginal distributions by considering the bivariate plot of *u_i_* versus *v_i_*, where *u_i_* is the rank of *x_i_* in the set *x_j_* (*j* = 1, …, *n*), divided by *n* + 1; and *v_i_* is defined in the same way but using the *y_i_*. The rank of the smallest element in a set is 1. The *u_i_* and *v_i_* are called *normalized ranks* of the *x_i_* and *y_i_*; they relate to the empirical cumulative distribution function of the *x_i_* and *y_i_*, respectively. We henceforth abbreviate “cumulative distribution function” as “distribution function” or “cdf”. See, for instance, Fig. 1C-D, which show the normalized ranks of A-B, respectively. Ranking makes the marginal distributions of the component datasets uniform, isolating the dependence structure. Dependence structure and marginals can then be studied separately. Normalized rank plots such as Fig. 1C-D relate to copulas in a way to be described in the “Background on copulas” section below (section 2), after a definition of copulas is provided. Note that Pearson correlation, even though it is the most commonly used measure of relationships between variables, is modified by monotonic tranformation of the component variables (Appendix S1, Fig. S1), and therefore reflects not only dependence information, but also information on the marginals (Genest & Favre 2007). Spearman and Kendall correlations, being rank-based, are not influenced by monotonic transformations of the component variables. They provide information solely on the dependence between those variables.

A main benefit of a copula approach is that it can detect associations in the tails of distributions, and asymmetries of such associations. Tail association (introduced formally in the “Background on copulas” section, below, section 2) is association between extreme values of variables. If smaller values of two positively associated variables are more strongly associated than are larger values, the variables are said to have stronger lower- or left-tail association than right- or upper-tail association (Fig. 1C); and *vice versa* if larger values are more strongly associated than smaller ones (Fig. 1D). Datasets of the same Spearman or Kendall correlation can have a range of tail associations (Fig. S2), which can be quantified (see sections on Background and methods for Q1 below, sections 2 and 4). Although copulas can be used to model the entire complex dependence structure of data, we have found tail associations to usefully describe the full structure of dependence, so tail association is one of the main foci of this paper.

The goal of this paper is to explore the potential for applications of copulas in ecology and to help determine to what extent ecological understanding may benefit generally from using copulas. We provide a basic introduction to copula concepts in the “Background on copulas” section below (section 2). Readers can find additional abbreviated (Genest & Favre 2007; Anderson *et al.* 2018) and comprehensive (Nelsen 2006; Joe 2014; Mai & Scherer 2017) introductions elsewhere. Copulas have already been used very effectively in a few studies to illuminate ecological phenomena (Valpine *et al.* 2014; Anderson *et al.* 2018; Popovic *et al.* 2018), but such usage is still rare. Our results suggest that environmental, ecological and evolutionary processes may commonly generate and transmit complex dependence structures, including asymmetric tail associations, and that greater use of copulas can illuminate the underlying processes. We believe copula approaches are among the tools all ecologists, including tropical ecologists, should be considering for analysis of their data in the 21st century.

One way we may expect, *a priori*, a copula study of complex dependence structures between ecological variables to better reflect ecological relationships relates to Liebig’s law of the minimum. Liebig’s law is the idea that growth is controlled not by total resources but by the resources which are scarcest relative to organism needs. If, for instance, the growth of a plant depends on soil nitrogen, N, and other factors, a plot of plant growth rates versus soil N may look more like Fig. 1A or C than like Fig. 1B or D. Nitrogen will control plant growth, producing a clear relationship, only when it is limiting, in the left portions of the distributions. The portions of the distributions that dominate the association are visible in a rank-based plot, but may not be revealed by standard correlation techniques.

A second reason why we may expect understanding of complex dependence structures to benefit ecological research is that prior work has demonstrated, using copulas, complex stucture in the spatial dependence of environmental variables (Serinaldi 2008; Li *et al.* 2013; Goswami *et al.* 2018; She & Xia 2018). One may expect an environmental variable measured through time in two locations to show strong tail associations between the locations if intense meteorological events are also widespread in their effects, as seems frequently to be the case: extreme values are associated with intense events, so happen in both places at the same time, whereas moderate values of the environmental variable may be associated with local phenomena, which differ between the locations. Spatial dependence in an environmental variable tends to beget spatial dependence in fluctuations of populations or other ecosystem variables (this is called *spatial synchrony*) which are influenced by the environmental variable. This influence is called the *Moran effect*. If complex dependence structure is transmitted, possibly in modified form, from environmental to ecological variables, then we may expect complex dependence structure to be a common feature of the spatial synchrony of population, community, biogeochemical and other environmentally influenced ecological variables. Synchrony is a phenomenon of substantial interest in ecology (Liebhold *et al.* 2004; Sheppard *et al.* 2016; Walter *et al.* 2017).

A third reason why we may expect an understanding of complex dependence structures to benefit ecological research is that such an approach may help illuminate the causal mechanisms between variables. For instance, if two species Sp1 and Sp2 are strong competitors, abundances of the two species across quadrats should be negatively related, as in Fig. 1E-F. If Sp1 is plotted on the horizontal axis and Sp2 on the vertical, then Fig. 1E may suggest Sp1 is the dominant competitor: when Sp1 is abundant, Sp2 is necessarily rare because it is suppressed; whereas when Sp1 is rare, Sp2 may be abundant, or may also be relatively rare due to limiting factors other than Sp1. Alternatively, Fig. 1F may suggest that Sp2 is the dominant competitor. Other causal hypotheses may produce similar dependence structure, and it is usually impossible to obtain complete information on causal pathways from analyses which are fundamentally based on associations. However, more detailed examination of the patterns of dependence of variables may well be combined with biological and other reasoning to rule out some causal hypotheses which could not be eliminated solely with standard correlation approaches. Popovic *et al.* (2018) has previously used copulas to help illuminate causal aspects of relationships between species.

A fourth reason why we may expect a copula understanding of complex dependence structures to be useful for ecology has to do with spatially aggregated or averaged quantities. Many ecological variables of applied importance depend on the spatial average or sum of local quantities. For instance, regional methane and CO_2_ fluxes are the sum of local fluxes; and the total economic value of a fishery is the sum of local catches. We will explore how details, which can be captured using copulas, of the dependence between fluctuations of local quantities can influence fluctuations of the spatial mean or sum, and how this may influence higher organizational levels in ecology and human concerns. To illustrate this idea we cite Li *et al.* (2013), who have demonstrated that the overall reliability of wind-generated electricity depends sensitively on details of the dependence between wind speeds at multiple generator sites. Spatially aggregated ecological variables may be subject to a similar effect, with possible consequences. For instance, if populations of a pest species in different locations are all positively associated and are also more strongly related to each other in their right tails, then local outbreaks will tend to occur together, creating regional epidemics. On the other hand, stronger left-tail associations in a pest species, even if overall correlation were unchanged, would have more benign effects.

We approached our overall goal of exploring the potential of copula approaches for ecology by addressing the following specific questions. (Q1) Do datasets in ecology have dependence structure distinct from that of a multivariate Gaussian/normal distribution (here called non-normal copula structure)? Do positvely associated ecological variables show tail associations distinct from those of a normal distribution, and in particular do they show asymmetric tail associations? Normal copula structure is assumed by standard approaches that use multivariate normal distributions or distributions obtained by transforming the marginals of a normal distribution. So Q1 asks whether ecological data contain dependency information distinct from a standard or default case. (Q2) If the answer to Q1 is “yes,” then what are some possible causes/mechanisms of non-normal copula structure and asymmetric tail associations in ecology? And (Q3), what are the consequences of non-normal copula structure and asymmetric tail associations for ecological understanding and applications? After section 2, ‘Background on copulas’, we address Q1 (sections 3 through 5 of the paper) by analyzing several datasets, including environmental, species-trait, phenological, population, community, and ecosystem functioning data, selected to span multiple sub-fields and organizational levels of ecology. We address Q2 (sections 6 through 7 of the paper), using simple models. We address Q3 (sections 8 through 9 of the paper) using both data and models. Section 10 is the Discussion. Multiple analyses are brought to bear on each question, and these are summarized diagramatically in Fig. 2, which can serve as a *post hoc* guide to the paper. Copula approaches have been used to great effect in neuroscience (Onken *et al.* 2009), bioinformatics (Kim *et al.* 2008), medical research (Emura & Chen 2016), direct study of environmental variables (Serinaldi 2008; Li *et al.* 2013; Goswami *et al.* 2018; She & Xia 2018), and finance (Li 2000), and have also been used very effectively, but rarely so far, in ecology (Valpine *et al.* 2014; Anderson *et al.* 2018; Popovic *et al.* 2018). We argue that benefits of wider usage of complete copula descriptions of dependence in ecology will be substantial.

**Figure 2:**
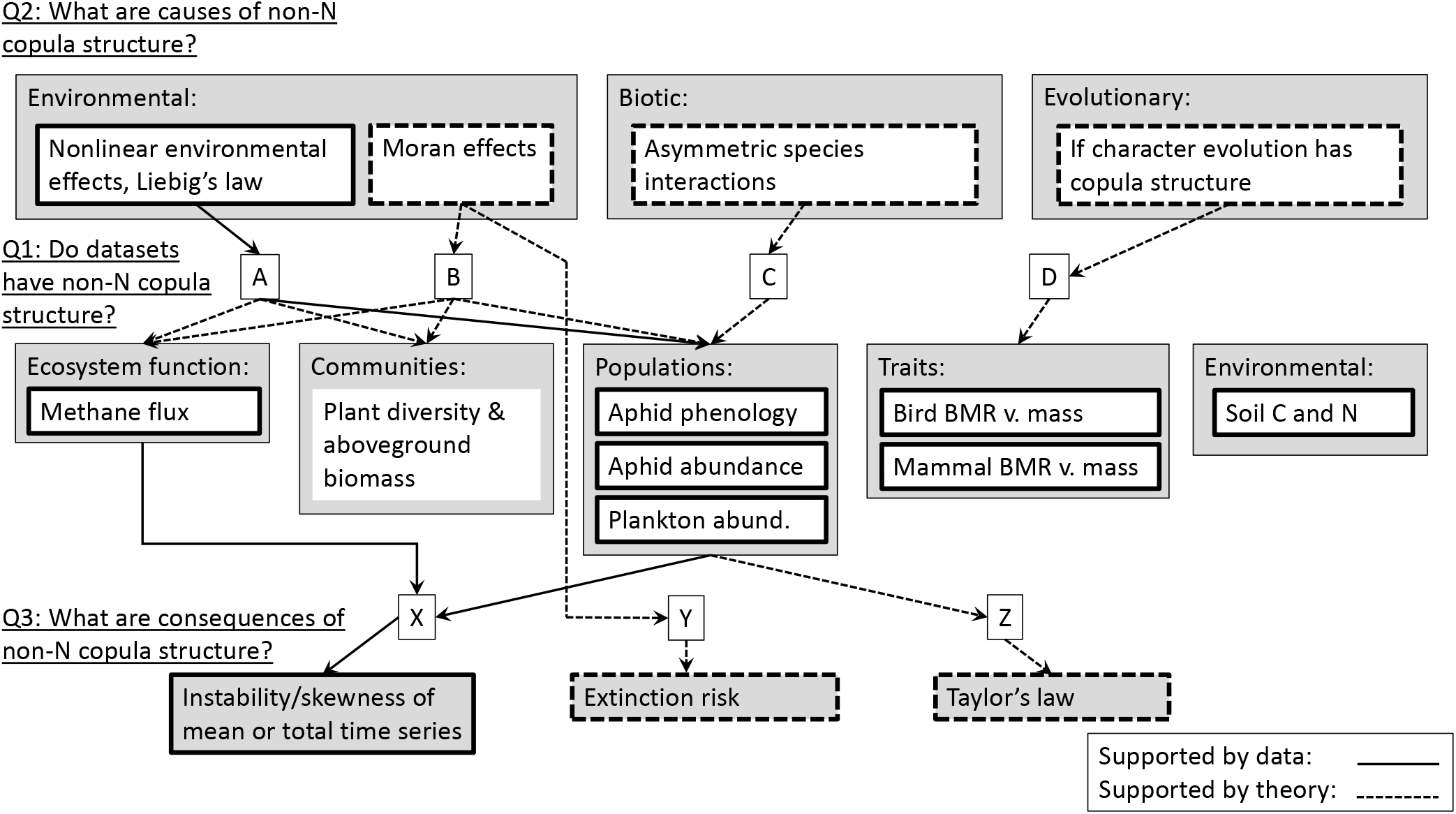
Summary and guide for analyses presented in the text. Middle boxes correspond to ecological datasets for which Q1 was examined (in sections 3-5, see also Table 1). Upper boxes correspond to potential causes of non-normal (non-N) copula structure that were examined (in sections 6-7 of the text). Lower boxes correspond to potential consequences we examined of non-normal copula structure for ecological understanding and for applications (in sections 8-9 of the text). Arrow labels (A-D for analyses pertaining to causes, Q2; and X-Z for analyses pertaining to consequences, Q3) link to locations in the text.

## 2 Background on copulas

We give a brief introduction to copulas, focusing, for simplicity, on bivariate copulas and on concepts needed for the rest of the paper. We assume familiarity with the basic language of probability distributions and random variables. See Nelsen (2006), Genest & Favre (2007), Joe (2014), and Mai & Scherer (2017) for general background on copulas, and see also Anderson *et al.* (2018), part of which is a short introduction to copulas for ecologists. A bivariate *copula* can be defined as a bivariate distribution function with both margins uniform on (0, 1) (Joe (2014), p. 7). This will be the distribution function of a bivariate random variable with uniform marginals, and the terminology *copula* is sometimes also applied to this random vector. A foundational theorem of Sklar (1959) (see also Mai & Scherer (2017), p. 16) says that if *F* is the bivariate distribution function of a random vector (*X, Y*), with margins *F_X_* and *F_Y_*, then there exists a copula *C* such that for all (*x, y*) in the Euclidean plane, *F*(*x,y*) = *C*(*F_X_* (*x*), *F_Y_* (*y*)). The theorem also says *C* is unique if *F_X_* and *F_Y_* are continuous. Thus *C* couples the bivariate distribution function, *F*, with the distribution functions of the marginals. Finally, Sklar’s theorem says that if *D* is any bivariate copula and *G_A_* and *G_B_* are univariate distribution functions, then *D*(*G_A_*(*x*), *G_B_* (*y*)) is a valid distribution function of some bivariate random variable. The applications of this study fall within the case in which univariate marginal distributions have continuous, strictly monotonic distribution functions, and this case is simpler. So we henceforth make such an assumption.

Sklar’s theorem implies that any random vector can be constructed from a unique copula and marginal distribution functions, and, furthermore, any copula and any univariate distribution functions give rise to a random vector; so Sklar’s theorem provides a correspondence between random vectors and pairs consisting of a copula and two univariate distribution functions, thereby making it possible to study a random vector by studying these two constituents. Marginals contain no information about the dependence structure of a random vector, so the copula contains all such information – it is a complete description of the dependence between variables. The univariate distribution functions associated with a random vector (*X, Y*) are its margins, *F_X_* and *F_Y_*, and the associated copula is the copula, *C*, guaranteed by Sklar’s theorem (and guaranteed unique, thanks to the assumptions of continuity and monotonicity made above). In fact, and making Sklar’s correspondence more concrete, we show in Appendix S2 that *C* is the bivariate distribution function of the random variable (*F_X_*(*X*), *F_Y_* (*Y*)). Conversely, given a copula *D* and univariate distribution functions *G_A_* and *G_B_*, Sklar’s theorem guarantees that *D*(*G_A_*(*x*), *G_B_* (*y*)) is a distribution function, so the random variable with this distribution function is the random variable corresponding to *D*, *G_A_* and *G_B_* under Sklar’s correspondence. Again making Sklar’s correspondence more concrete, we show in Appendix S2 that this random variable is 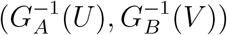, where 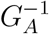 and 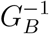 are the inverses of *G_A_* and *G_B_* and (*U, V*) is the random variable with distribution function *D* (which has uniform marginals). If we conflate a copula with the random vector of which the copula is the distribution function, then Sklar’s correspondence is between random variables, (*U, V*), with uniform marginals and random variables, (*X, Y*), with arbitrary continuous, strictly monotonic marginals. The correspondence is simply via the application of the univariate distribution functions, or their inverses: 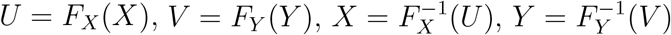. Application of the distribution functions of the component random variables translates between the uniform-marginal context, (*U, V*), which isolates the dependence information, and the context of the original random variable, (*X, Y*).

Thus Sklar’s theorem makes it possible to construct bivariate distributions in two separate steps: by specifying the marginal distributions and by specifying the dependence structure. For instance, construction of several bivariate distributions is carried out in this way in Fig. 3. The contrast between Fig. 3C and D illustrates in particular how a bivariate distribution can be changed, while retaining the same marginals, by changing the copula. Fig. 3C is a bivariate normal distribution, but Fig. 3D clearly is not, although its marginals are the same. Fig. 3D shows stronger association of the two variables in the left than in the right tails. Sklar’s theorem also makes it possible to study and model dependence separately from marginal distributions. For instance, in a pedagogical example, Anderson *et al.* (2018) modelled the dependence between abundance data for two fish species, red moki and black angelfish, by first modelling the marginal distributions and then modelling the dependence using Gaussian copulas (their section 2). See the Discussion for a few words on multivariate copulas, which can be developed in the same way as above (Nelsen 2006; Joe 2014; Mai & Scherer 2017; Anderson *et al.* 2018).

**Figure 3:**
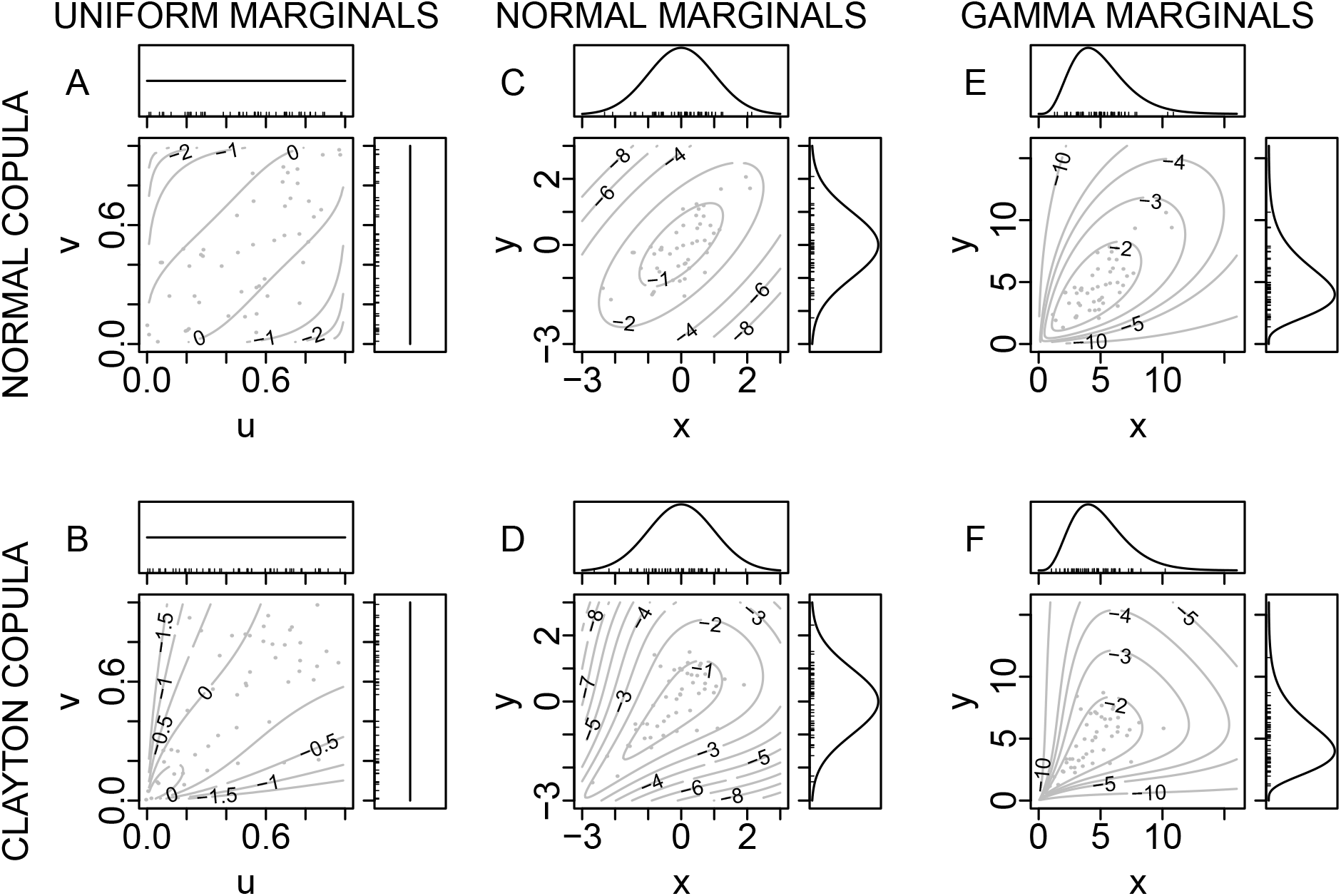
Normal (A) and Clayton (B) copulas were combined with standard normal marginals (C, D) and gamma marginals (E, F) via Sklar’s theorem to produce bivariate distributions. Note that a normal copula is distinct from a normal marginal distribution: a normal copula is the copula of a bivariate normal distribution. See section 2 for more information on normal, Clayton, and other copulas. Each copula was used with both sets of marginals and each set of marginals was used with both copulas, to demonstrate that: both copula and marginals contribute fundamentally to the resulting distribution, the copula contributing the information on the dependence between the variables; and that one can select the copula and marginals independently. Bivariate distributions, including the copulas, are depicted via their log-scale probability density functions (pdfs), and marginals are depicted via their linear-scale pdfs. Grey dots are 50 random samples from each distribution. The parameter 0.7 was used for the normal copula, the parameter 2 for the Clayton copula, and shape and scale parameters 5 and 1 for the gamma marginals. Variables *u* and *v* were used for copulas and *x* and *y* for distributions created by pairing a copula with (non-uniform) marginals.

We introduce some standard copula families, as examples of copulas and also because we will fit these families to data later. The bivariate normal copula family, which consists of copulas of bivariate normal distributions, is a one-parameter family. The parameter, which we here call *p*, corresponds to the correlation of the related bivariate normal distribution, and controls the degree of association between the variables. A normal copula with *p* = 0.7 was already introduced (Fig. 3A). The formula for the copula is 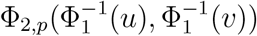, where 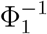 is the inverse of the cummulative dstribution function (cdf) of a (univariate) standard normal and Φ_2,*p*_ is the joint cdf of a bivariate normal distribution with mean (0, 0) and covariance matrix having 1s on the diagonal and *p* in the off diagonal positions. However, formulas for copulas are often not as directly informative about copula properties as probability density function (pdf) pictures; Fig. 4 has example pdfs corresponding to bivariate normal copulas for various values of *p*. Note that the pdfs are symmetric around the diagonal line *v* = − *u* + 1 (Fig. 4), so normal copulas have symmetric associations between the two variables in the left and right tails. The Clayton copula family, of which we have already pictured an example (Fig. 3B), is another one-parameter family. In contrast to normal copulas, Clayton copulas have stronger left-than right-tail association. The formula is [max(*u*^−*p*^ + *v* ^−*p*^ − 1, 0)]^−1/*p*^, for parameter *p*; though this again provides less direct intuition than do example pdfs (Fig. 5). Higher values of the parameter, *p* produce copulas from the Clayton family with higher Kendall or Spearman correlation. The survival Clayton family is the symmetric opposite of the Clayton family, showing stronger right-than left tail association (Fig. S3). The BB1 copula family is a two-parameter family, thereby providing more flexibility: for some parameters BB1 copulas have stronger left-than right-tail association, and for others the reverse (Fig. 6). See Nelsen (2006) and Joe (2014) for further information on these and other copula families.

**Figure 4:**
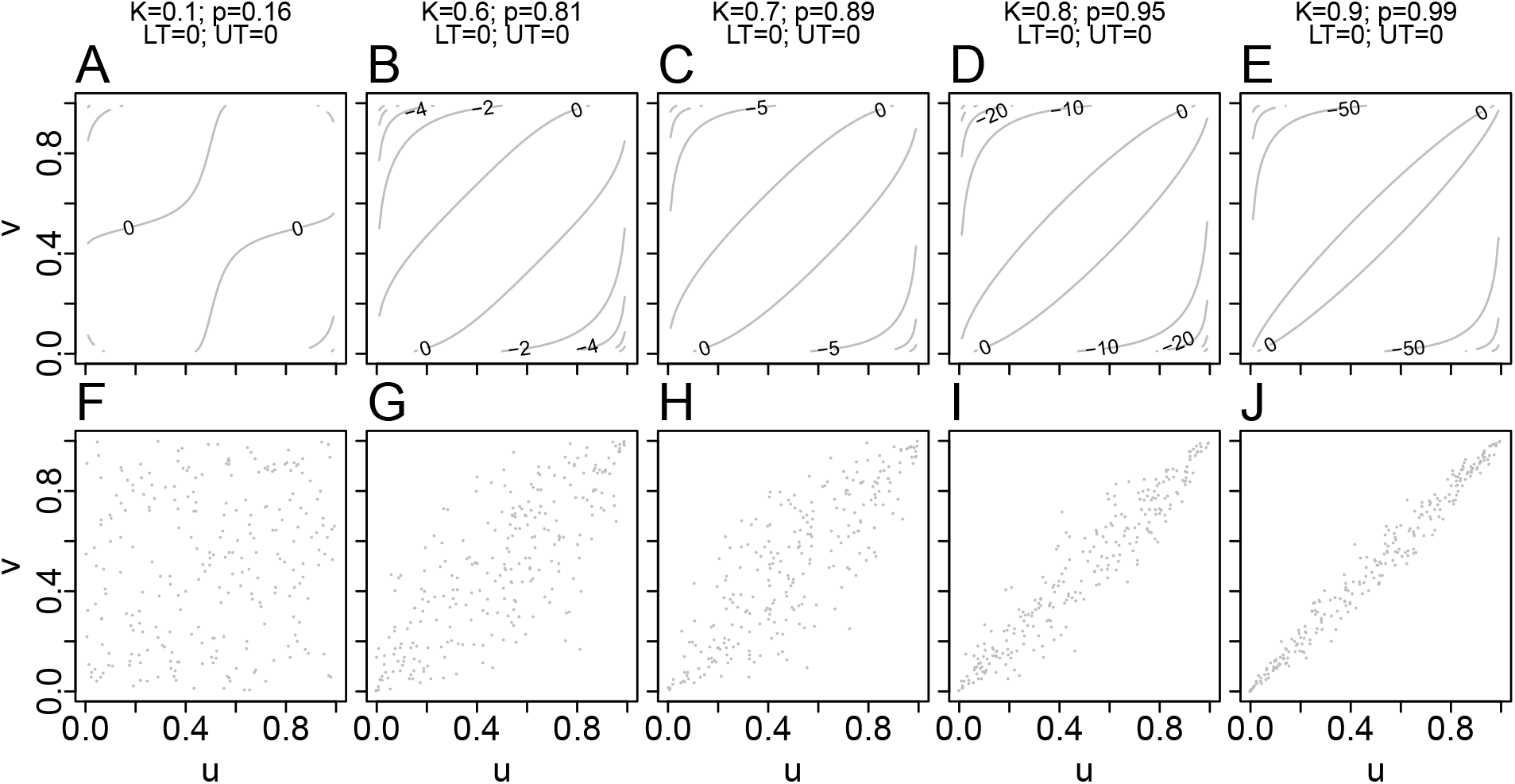
Log-transformed pdfs (A-E) and samples (F-J) from example normal copulas. K is Kendall correlation; *p* is the value of the parameter for the normal family (it is a one-parameter family); and LT and UT are the measures of lower- and upper-tail dependence, respectively (for the definitions of these, see the section on Background on copulas, section 2). The parameter range for the family is *p* ∈ [−1, 1], and lower- and upper-tail dependence are always 0.

**Figure 5:**
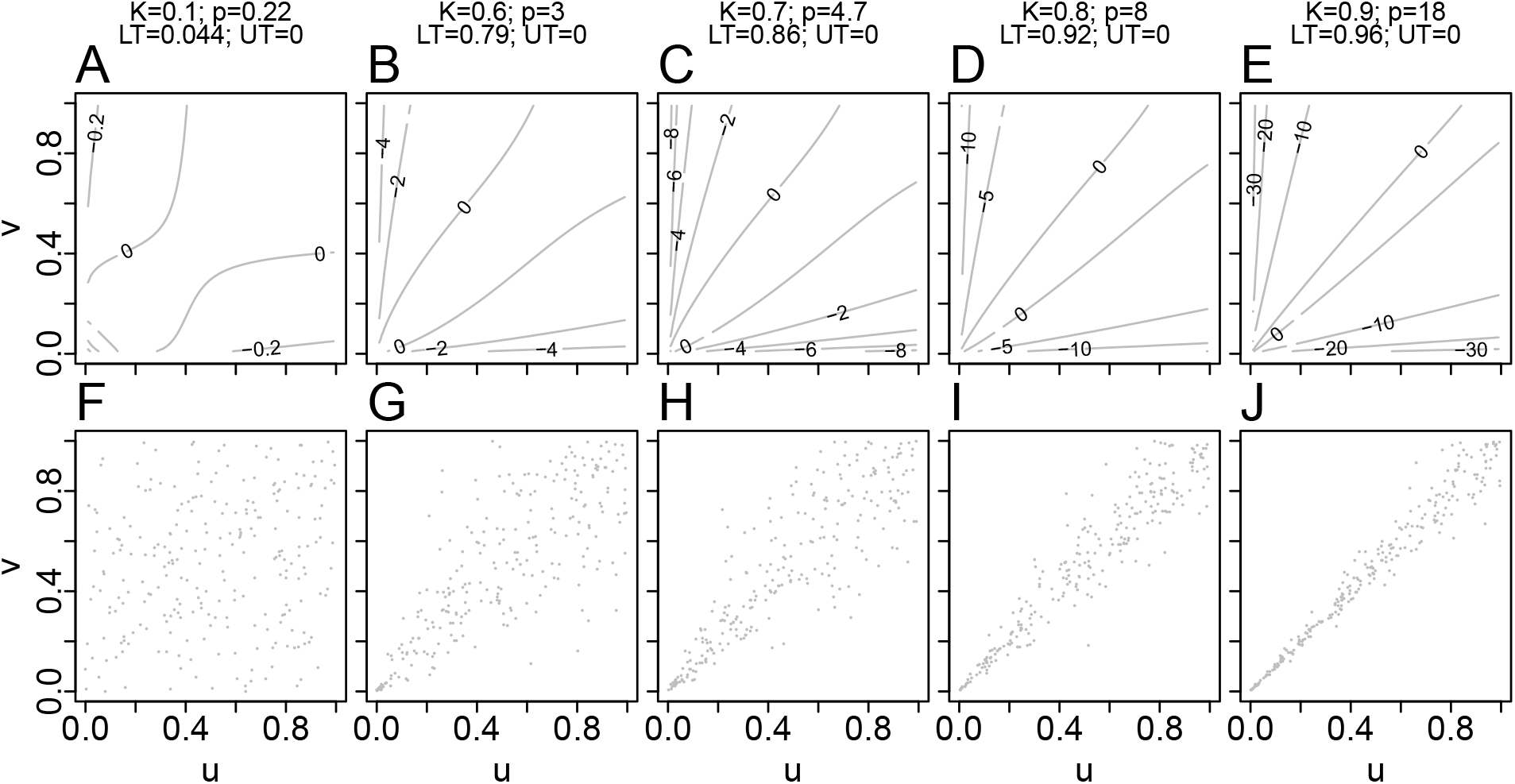
Log-transformed pdfs (A-E) and samples (F-J) from example Clayton copulas. K is Kendall correlation; *p* is the value of the parameter for the Clayton family (it is a one-parameter family); and LT and UT are the measures of lower- and upper-tail dependence, respectively (for the definitions of these, see the section on Background on copulas, section 2). The parameter range for the family is *p* ∈ (0, ∞), lower-tail dependence is 2^−1/*p*^ and upper-tail dependence is 0.

**Figure 6:**
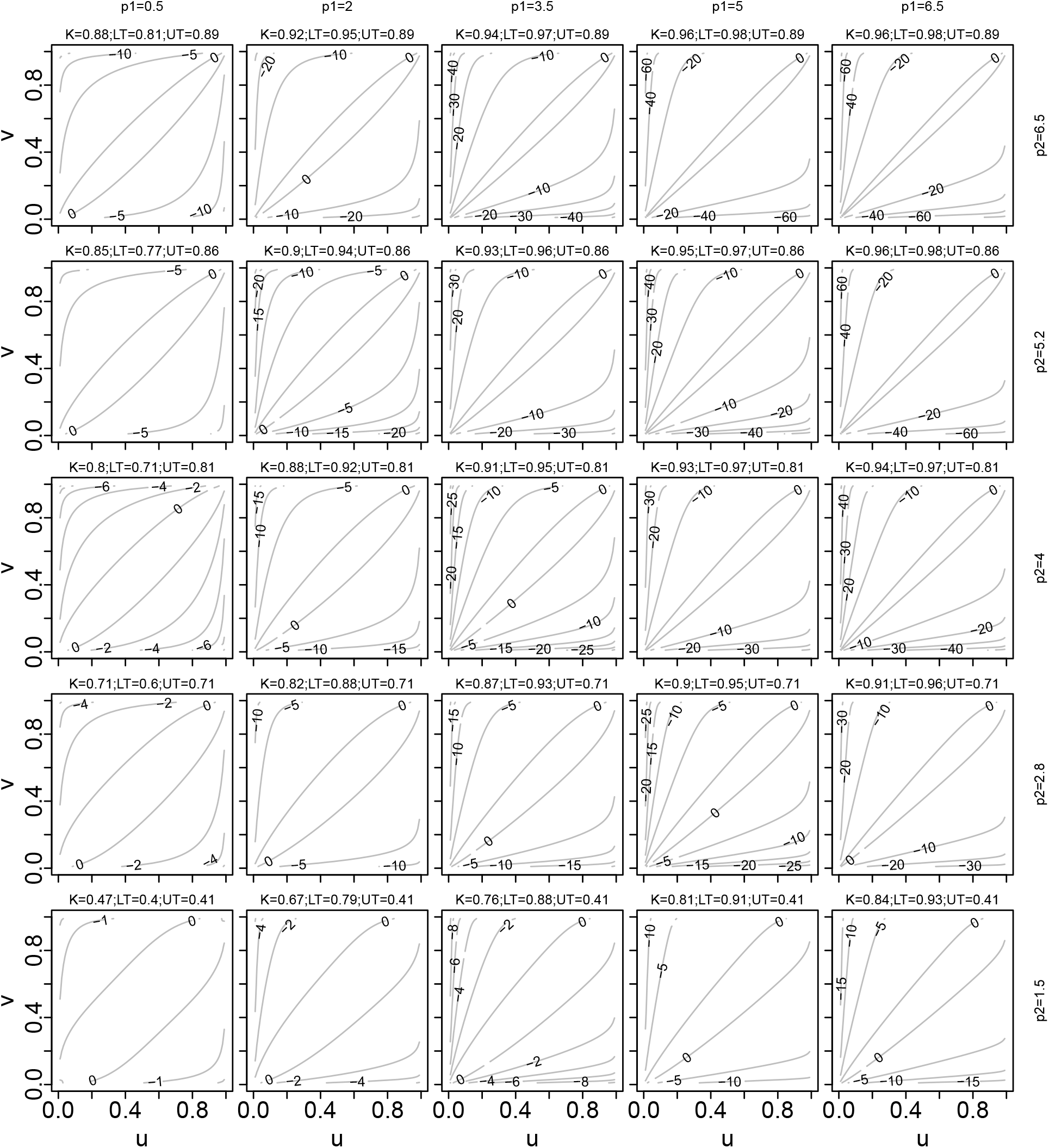
Log-transformed pdfs for example BB1 copulas. K is Kendall correlation; *p*_1_ and *p*_2_ denote the two parameters of the family (it is a two-parameter family); and LT and UT are our measures of lower- and upper-tail dependence, respectively (for the definitions of these, see the section on Background on copulas, section 2). The parameter ranges for the family are *p*_1_ ∈ (0, ∞) and *p*_2_ ∈ [1, ∞), lower-tail dependence is 2^−1/(*p*_1_ *p*_2_)^ and upper-tail dependence is 2 − 2^1/*p*_2_^.

Having introduced tail association conceptually and referred to it in examples, we now need a more precise definition of a measure of tail association. We use the term *tail association* to describe the general idea of the strength of associations between two variables in the tails of their distributions. One of the ways this is measured is called *tail dependence* (Joe (2014), section 2.13; Nelsen (2006), section 5.4), which we here define. Given a random vector (*X, Y*) with margins *F_X_*, *F_Y_* and defining *U* = *F_X_* (*X*), *V* = *F_Y_* (*Y*), the upper-tail dependence of *X* and *Y* is defined as 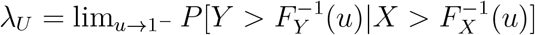. This equals lim_*u*→1−_ *P* [*U* > *u*|*V* > *u*], which in turn equals lim_*u*→1−_ *P* [*U* > *u, V* > *u*]/*P* [*V* > *u*] = lim_*u*→1−_ *P* [*U* > *u, V* > *u*]/(1 − *u*), which shows upper-tail dependence is defined symmetrically in the two random variables. All the variables we consider were positively associated when they were significantly associated, so we think of positively associated variables when conceptualizing the definitions presented here. Lower tail dependence is defined analogously, as 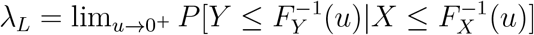. Tail dependence is a property of the random variable (*U, V*), so depends only on the copula and not on the marginals combined with it. It can be shown (Nelsen (2006), section 5.4) that *λ_L_* = *λ_U_* = 0 for the normal copula, *λ_L_* = 2^−1/*p*^ and *λ_U_* = 0 for a Clayton copula with parameter *p* > 0, and *λ_L_* = 0 and *λ_U_* = 2^−1/*p*^ for a survival Clayton copula with parameter *p* > 0. In section 4 we will introduce ways tail dependence can be applied to data, by first fitting copulas to the data. We will also introduce nonparametric measures of tail association that apply directly to data.

Plots using normalized ranks were proposed in the Introduction as being conceptually similar to copulas. We here explain the connection. As stated above, the copula associated with a random variable (*X, Y*) is the joint distribution function of the random variable (*F_X_*(*X*), *F_Y_*(*Y*)). Given a sample (*x_i_*, *y_i_*), *i* = 1,…, *n*, from (*X,Y*), let 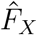 and 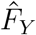 be the empirical cdfs associated with the *x_i_* and *y_i_*, respectively. These are step functions of *x* and *y*, respectively, that start at *0* for low values of *x* and *y* and jump by 1/*n* at each of the data points. Therefore 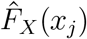 is the rank of *x_j_* in the set {*x_i_* : *i* = 1,…, *n*}, dividided by *n*, and analogously 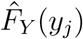 is the rank of *y_j_* in the set {*y_i_* : *i* = 1,…, *n*}, divided by *n*. For large *n*, 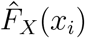 and 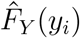 approximate the cdfs *F_X_* and *F_Y_*, respectively. But 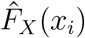 equals the normalized rank of *x_i_* times *n*/(*n* + 1), and likewise 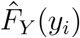 equals the normalized rank of *y__i__* times *n*/(*n* + 1) So the normalized rank in turn approximates the empirical cdf and therefore the cdf. Thus the normalized rank pairs (*u_i_*, *v_i_*) for *i* = 1, …, *n* can be regarded as approximate samples from the random variable (*F_X_*(*X*), *F_Y_*(*Y*)), which is the random variable associated with the copula of (*X, Y*). The scatterplot of *v_i_* versus *u_i_* can be used to infer aspects of copula structure (see section 4 below).

## 3 Data

The datasets we used included environmental, species-trait, phenological, population, community, and ecosystem functioning data (Table 1), selected to span multiple fields and levels of organization within ecology. Copula structure describing the relationship between atmospheric weather variables such as rainfall or wind speed, as measured in multiple locations through time, has been examined previously in the meteorological literature (e.g., Serinaldi 2008; Li *et al.* 2013). We therefore examined environmental variables from the soil instead, using the Rapid Carbon Assessment database (RaCA; Wills *et al.* 2014). The database comprises measurements of soil organic carbon and total soil nitrogen (Mg C or N per hectare of soil surface) from 5907 locations across the coterminous United States (Fig. S4 for sampling locations). Species-trait data were average species basal metabolic rate (BMR, KJ per hour) and body mass (grams) for 533 species of birds (McNab 2009) and 638 species of mammals (McNab 2008). These data have been much studied, but to our knowledge the copula structure of the dependence between metabolic rate and body mass has not been examined. Species-trait data such as these reflect the coevolution of the two traits. Phenological data were aphid first-flight dates from 10 locations (Fig. S5) across the United Kingdom (UK) for 20 aphid species (Table S1), for the 35 years 1976 to 2010. These time series were computed from the Rothamsted Insect Survey (RIS) suction-trap dataset (Harrington 2014; Bell *et al.* 2015). The first of our two population-level datasets was also derived from the RIS suction-trap data, and comprised total counts of aphids trapped for the same locations, species, and years. Our second population dataset comprised average annual plankton abundance estimates for 14 locations (Fig. S6) in the North Sea and British seas for 22 taxa (Table S2) for the 56 years 1958 to 2013. The locations are 2^°^ by 2^°^ grid cells.

**Table 1:** Summary table for the data we used. Bold entries are the multivariate data, the rest are bivariate datasets (see section efData). Basal metabolic rate=BMR.

| Data | Category | No. of measurements we used | References |
| --- | --- | --- | --- |
| Soil C and N | Environmental | 5907 locations | US Rapid Carbon Assessment database (RaCA), Wills et al. (2014) |
| Bird body masses and BMR | Species-trait | 533 birds | McNab (2009) |
| Mammal body masses and BMR | Species-trait | 638 mammals | McNab (2008) |
| **Green spruce and other aphid species abundances** | **Population** | **10 locations with at least 30-yr. timeseries for each of 20 sp.** | **Rothamsted Insect Survey** |
| **Leaf-curling plum and other aphid first flight dates** | **Phenological** | **10 locations with at least 30-yr. timeseries for each of 20 sp.** | **Rothamsted Insect Survey** |
| ***Ceratium furca* and other plankton taxa abundances** | **Population** | **14 locations with at least 45-yr. timeseries for each of 22 taxa** | **Continuous Plankton Recorder Survey** |
| Plant diversity and aboveground biomass | Community level | 168 plots | Cedar Creek Ecosystem Science Reserve, biodiversity experiment e120 |
| **Methane-flux** | **Ecosystem functioning** | **13 locations with at least 50 dates of data** | **Great Miami Wetland Mitigation Bank, Smyth et al. (2019)** |

These data were computed from the Continuous Plankton Recorder survey hosted by the UK Marine Biological Survey. Community-level data, obtained from the Cedar Creek Ecosystem Science Reserve, were plant aboveground biomass (Tilman 2018a) and Shannon’s diversity index (computed from plant species percent cover data (Tilman 2018b)) for 168 plots (Tilman 2018c), each 9m by 9m, as sampled in the years 1996-2000 and 2007, each year analyzed separately (Tilman *et al.* 2001, 2006). Finally, ecosystem functioning data were methane (CH_4_) fluxes between the soil or water surface and the troposphere, measured at 13 locations at daily to weekly intervals from September 2015 to September 2016 at the Great Miami Wetland mitigation bank in Trotwood, Ohio (Holland *et al.* 1999; Jarecke *et al.* 2016; Smyth *et al.* 2019). Each included location was measured on at least 50 dates. See Appendix S3 for additional data details.

Our environmental, species-trait, and community datasets happen to be “bivariate” datasets in the sense that they comprise two quantities measured at different locations or for different species (Table 1). Our phenology, population, and ecosystem functioning datasets are “multivariate” in that they comprise, for each taxon, measurements through time at multiple locations; copula structure can be studied for each pair of locations.

## 4 Concepts and methods for Q1

We addressed Q1 first via a model selection procedure in which several families of copulas were fitted to our ecological datasets and fits were compared via the Akaike and Bayesian Information Criteria (AIC and BIC). One of the fitted copulas was the normal copula, making possible comparisons of the degree to which the normal copula versus other copula families were good descriptions of data.

For bivariate datasets (*x_i_*, *y_i_*) for *i* = 1, …, *n*, model selection involved several steps. First, we produced normalized ranks *u_i_* and *v_i_* as in the Introduction. Second, we tested the independence of the *u_i_* and *v_i_* using the statistic 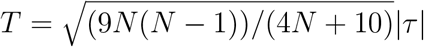, where *τ* is Kendall’s tau for the data. Genest & Favre (2007) argue that this statistic is approximately standard normally distributed. We used the implementation of this test given in the `BiCopIndTest` in the `VineCopula` package in the R programming language. We tested for independence because our model selection algorithms were ineffective if data could not be distinguished from independent data since many of the copula families we considered include the independent copula. If independence could be rejected (0.05 significance level), model selection proceeded. Third, we fit 16 bivariate copula families (see below for information on the families used) to the normalized ranks via maximum likelihood. The approach of fitting copula families to normalized ranks was studied in depth and recommended by Genest *et al.* (1995) and Shih & Louis (1995). Their estimator of copula parameters, which we use, was shown to be consistent, asymptotically normal, and fully efficient at indepencence (Genest *et al.* 1995). Genest & Favre (2007) recommend carrying out inferences of dependence structures (which was our goal here) using normalized ranks. We used the fitting implementation given in `BiCopEst` in `VineCopula`. Fourth, we obtained AIC and BIC values and accompanying model weights AIC_w_ and BIC_w_ (Burnham & Anderson 2003) for each fitted copula. `BiCopEst` also provided lower- and upper-tail dependence of the best-fitting member of each family. AIC_w_ values were used to get model-averaged lower- and upper-tail dependence values using standard model averaging formulas (Burnham & Anderson 2003); likewise for BIC.

Thus for each bivariate dataset, the end products of the procedure just described were threefold, corresponding to the three parts of Q1 listed in the Introduction: A) the AIC, BIC, AIC_w_ and BIC_w_ values for each of our 16 copula families (see below for list), including the normal family, providing an inference as to whether the normal copula or an alternative was better supported by data; B) lower- and upper-tail dependence measures for each fitted copula and model-averaged tail dependence measures, providing information on whether, and to what extent, tail dependence differed from the tail dependence of a normal copula (i.e., 0); and C) the difference of lower- and upper-tail dependence measures for each fitted copula and model averages of those quantities, providing information on whether, and to what extent, tail dependence was asymmetric.

Model selection methods give the *relative* support of several models, but do not indicate whether any of the models are an objectively good fit. To test that, we tested the goodness of fit of our AIC-best copula family using a bootstrapping procedure developed and studied by Wang & Wells (2000) and Genest *et al.* (2011) and implemented as `BiCopGofTest` of `VineCopula`. The procedure performed one test based on a Cramer-von Mises statistic and another based on a Kolmogorov-Smirnov statistic. To keep computation times reasonable, a run using 100 bootstraps was performed; if the *p*-value from either test was less than 0.2, tests were re-run with 1000 boostraps.

We fit 16 bivariate copula families, exhibiting a variety of lower- and upper-tail dependence characteristics, with bivariate datasets. The purpose of using a large collection of families was solely to include a variety of alternative dependence structures to have a robust model selection and multi-model inference procedure. For that purpose, it is not important for the reader to understand the details of these copulas, and, additonally, these copulas have been described in more detail elsewhere (Joe 1997); so we only briefly identify each family, say a few words about tail dependence of the families, and provide pictorial descriptions in the Appendices. We used several families that can exhibit positive lower-tail dependence (of strength depending on parameters) and zero upper-tail dependence: the Clayton, survival Gumbel, survival Joe and survival BB6 families. We used several families that can exhibit positive upper-tail dependence (strength depends on parameters) and zero lower-tail dependence: the survival Clayton, Gumbel, Joe and BB6 families. We used families that show zero upper- and lower-tail dependence: the normal and Frank families. These families have pdfs symmetric about the line *v* = −*u* + 1. We also used several families that can show both upper- and lower-tail dependence, in relative amounts that depend on parameter values: the BB1, survival BB1, BB7 and survival BB7 families. We also used the BB8 family, which shows zero lower-tail dependence, and zero upper-tail dependence except for a boundary case for the parameters. And we used the survival BB8 copula, which shows zero upper-tail dependence, and zero lower-tail dependence except for a boundary case. “Survival” families are rotations of the copula with the similar name by 180 degrees. See Joe (1997) for details on all these families. We used the implementations of these families provided in the `VineCopula` package for the R programming language. See Figs 4-6 and S3 and S7-S18 for visual depictions of the pdfs of these copulas and how pdfs and tail dependence are influenced by the parameters.

For multivariate datasets, we performed the bivariate analysis for all possible pairwise combinations of distinct locations. We carried out pairwise bivariate analyses instead of trying to fit a multivariate copula, for simplicity and because that approach was sufficient to answer our research questions; but see the Discussion for a few words on multivariate copulas. We present the number of pairs for which a non-normal copula was the AIC-best copula, and we characterize AIC differences between the normal and AIC-best copulas across location pairs. We also computed the model-averaged lower- and upper-tail statistics, and differences between these, for each pair of locations, and we characterize the distributions of these values across location pairs.

In addition to our model selection approach, we also used a nonparametric approach, to provide greater confidence in our answers to Q1. We used three statistics which quantify the extent to which the normalized ranks *u_i_* and *v_i_* are related in any part of their distributions. We here describe the statistics, with some additional details in Appendix S4. The statistics are defined with positively associated variables in mind. All our variables were positively associated when they were significantly associated. Given two bounds 0 ≤ *l_b_* < *u_b_* ≤ 1, define the lines *u* + *v* = 2*l_b_* and *u* + *v* = 2*u_b_*, which intersect the unit square (Fig. 7). Our statistics quantify the association between *u_i_* and *v_i_* in the region bounded by these lines. Using *l_b_* = 0 and *u_b_* ≤ 0.5 gives information about association in the left parts of the distributions of *u* and *v*, and using *u_b_* = 1 and *l_b_* ≥ 0.5 gives information about association in the right parts. The first statistic, a partial Spearman correlation,

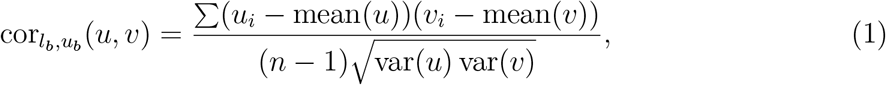

is the portion of the Spearman correlation of *u_i_* and *v_i_* that is attributable to the points between the bounds. Here sample means and variances are computed using all *n* data points, but the sum is over only the indices *i* for which *u_i_* + *v_i_* > 2*l_b_* and *u_i_* + *v_i_* < 2*u_b_*. Larger values of the partial Spearman correlation indicate stronger positive association. The sum of cor_0,0.5_(*u, v*) and cor_0.5,1_(*u, v*) (or some other choice of cor*_l_b_k__,u_b_k___* (*u, v*) for bounds *l_b_k__,u_b_k__* that partition the interval (0, 1)) equals the standard Spearman correlation as long as no points fall exactly on the bounds. We also defined a statistic P*_l_b_,u_b__* (Appendix S4), which has a similar qualitative interpretation to cor*_l_b_,u_b__*. Our third statistic, 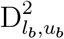 is the average squared distance between points satisfying *u_i_* + *v_i_* > 2*l_b_* and *u_i_* + *v_i_* < 2*u*_*b*_ and the line *v* = *u*. Unlike cor*_l_b_,u_b__* and P*_l_b_,u_b__*, for which large values indicate strong association between the bounds, small values of 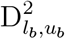 indicate strong association. These statistics are not estimators of the tail dependence quantities defined previously, but rather are conceptually similar measures of associations in the tail portions of the distributions when appropriate values of *l_b_* and *u_b_* are selected.

**Figure 7:**
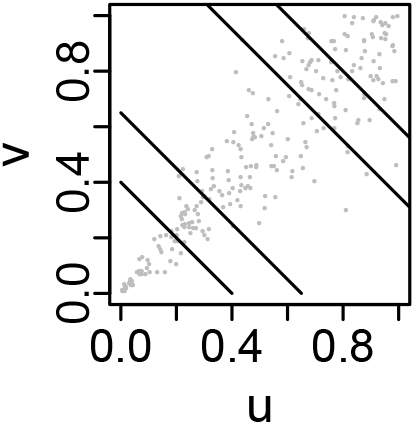
The partial Spearman correlation, cor*_l_b_,u_b__* (*u, v*), within a band can be computed for any band (section 4) to describe how the strength of association between *u* and *v* varies from one part of the two distributions to another, as can the statistics P*_l_b_,u_b__*(*u, v*) and 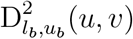. Diagonal lines show two bands, the data in the left/lower band showing stronger association than those in the right/upper band.

For large datasets (large *n*), we used *l_b_* and *u_b_* close together without incurring undue sampling variation in our statistics, and we considered multiple bands (*l_b_, u_b_*) to understand how association varies across different parts of the distributions. But for datasets with smaller *n* we considered only *l_b_* = 0, *u_b_* = 0.5 and *l_b_* = 0.5, *u_b_* = 1, abbreviating cor_*l*_ = cor_0,0.5_ (*l* is for “lower”) and cor_*u*_ = cor_0.5,1_ (*u* is for “upper”). Likewise P_*l*_ = P_0,.5_, P_*u*_ = P_0.5,1_, 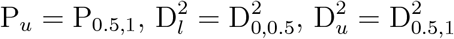. To test for asymmetry of association in upper and lower portions of distributions, we used differences cor_*l*_ – cor_*u*_, P_*l*_ – P_*u*_ and 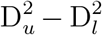 for smaller datasets (note the opposite order in the last of these); and cor_0,*u*_*b*__ – cor_1−*u*_*b*_,1_, P_0,*u*_*b*__ – P_1−*u*_*b*_,1_ and 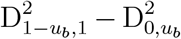 with *u_b_* close to 0 for large datasets. We tested our statistics in Appendix S5 (see also Figs S19-S20).

For bivariate datasets, we compared values of the statistics 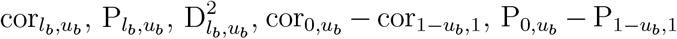 and 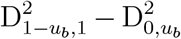 to distributions of values of the same statistics computed on surrogate datasets that were produced from the empirical data by randomizing it in a special way to have no tail dependence (Appendix S6). The surrogate/randomized datasets had exactly the same marginal distributions as the empirical data and had very similar Kendall or Spearman correlation (the surrogate algorithm had two versions, one for preserving each correlation coefficient), but had normal copula structure. Thus our comparisons tested the null hypothesis that our statistics took values on the empirical data no different from what would have been expected if the copula structure of the data were normal, but the data were otherwise statistically unchanged. The comparison of one of the statistics 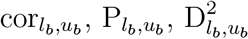, as computed on the empirical data, to the distribution of its values computed on surrogate datasets provides a test of whether association between the variables in the part of the distributions specified by *l_b_* and *u_b_* is different from what would be expected from a null hypothesis of normal copula structure. Thus the comparison addresses the first two parts of Q1 for bivariate datasets. Significant deviations correspond to deviations from normal copula structure. In particular, deviations using *l_b_* = 0 and *u_b_* small (say, 0.1) correspond to lower-tail associations different from that of a normal copula; likewise, using *l_b_* = 0.9 and *u_b_* = 1 tests for upper-tail associations different from that of a normal copula. The comparison of one of the statistics 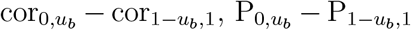 and 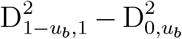, as computed on the empirical data, to the distribution of its values on many surrogate datasets provides a test of asymmetry of tail associations. Thus this comparison helps address the third part of Q1 for bivariate datasets.

For multivariate datasets, we calculated Spearman and Kendall correlations and the statistics cor_*l*_, cor_*u*_, P_*l*_, P_*u*_, 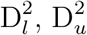, cor_*l*_ – cor_*u*_, P_*l*_ – P_*u*_, and 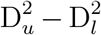 for all pairs of sampling locations, and we then computed means across the pairs for each statistic. We used a spatial resampling scheme (Appendix S7) to calculate confidence intervals of these means. The scheme is identical to that proposed by Bjørnstad & Falck (2001) for application in their now widely used spline correlogram method. The same scheme was also used by Walter *et al.* (2017) as part of a method for performing model selection among matrix regression models. Code and data for this project are archived at www.github.com/sghosh89/BIVAN.

## 5 Results for Q1: Ecological datasets have non-normal copula structure and asymmetric tail associations

Copula structures were non-normal and showed asymmetric tail associations for most, but not all datasets we examined, answering Q1 in the affirmative. To make results easier to absorb, we first present results for an example bivariate dataset, then we summarize results for all bivariate datasets, then we present results for an example multivariate dataset, then we summarize for all multivariate datasets.

For the soil C and N data (section 3, Table 1, Fig. 8A), variables were non-independent (*p* = 0, to within the precision available from `BiCopIndTest` in the `VineCopula` package). Non-independence is also visually apparent. The Kendall correlation was 0.6. We fitted our 16 copulas to the normalized ranks (Fig. 8E) and computed AIC and BIC weights and corresponding lower- and upper-tail statistics for the fitted distributions (Table 2). BB1 was the minimum-AIC copula, with AIC −6708.25, whereas the normal copula had a much higher AIC, −6333.16, answering the first part of Q1 for these data. Recall that AIC differences of 2 or 3 are considered meaningful and differences of 8 or greater are considered definitive, so that data provide essentially no support for the higher-AIC model in that case (Burnham & Anderson 2003). Model-averaged lower- and upper-tail dependence statistics were 0.453 and 0.615, respectively (AIC weights were used for averaging). These values are distinct from what a normal copula would give, namely, 0, helping to answer the second part Q1 for these data. The values also differed substantially from each other, helping answer the third part of Q1. The numbers reflect stronger upper-than lower-tail dependence, and this is also visible in the extreme upper-right and lower-left corners of the copula plot (Fig. 8E).

**Figure 8:**
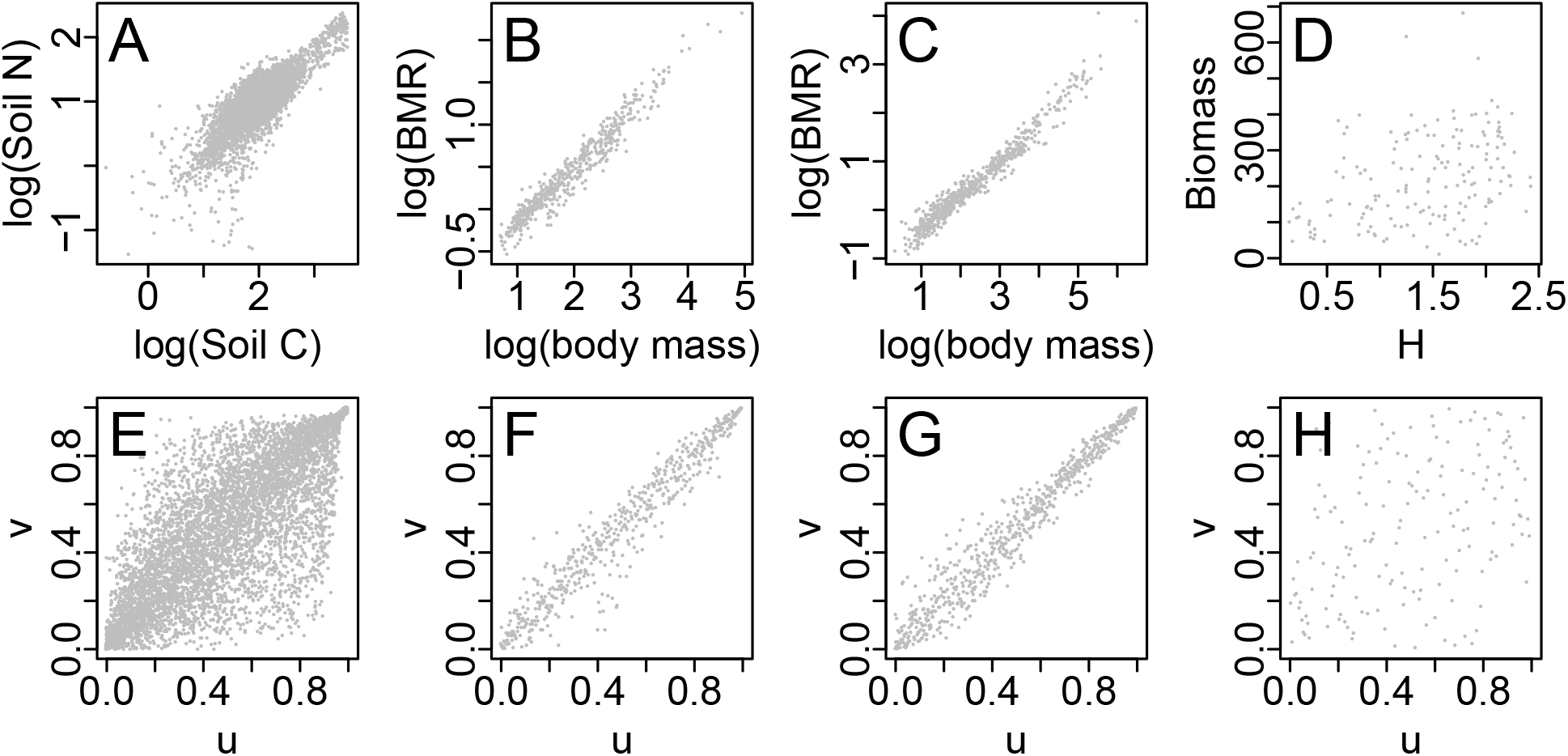
Upper panels show raw data plots for (A) log_10_(soil C) and log_10_ (soil N) data, (B) log_10_ (basal metabolic rate (BMR)) vs. log_10_ (body mass) for birds, (C) the same for mammals, and (D) above-ground plant biomass vs. Shannon’s index (*H*) from Cedar Creek. Bottom panels show the corresponding normalized rank plots. See section 3 for units used in upper panels.

**Table 2:**
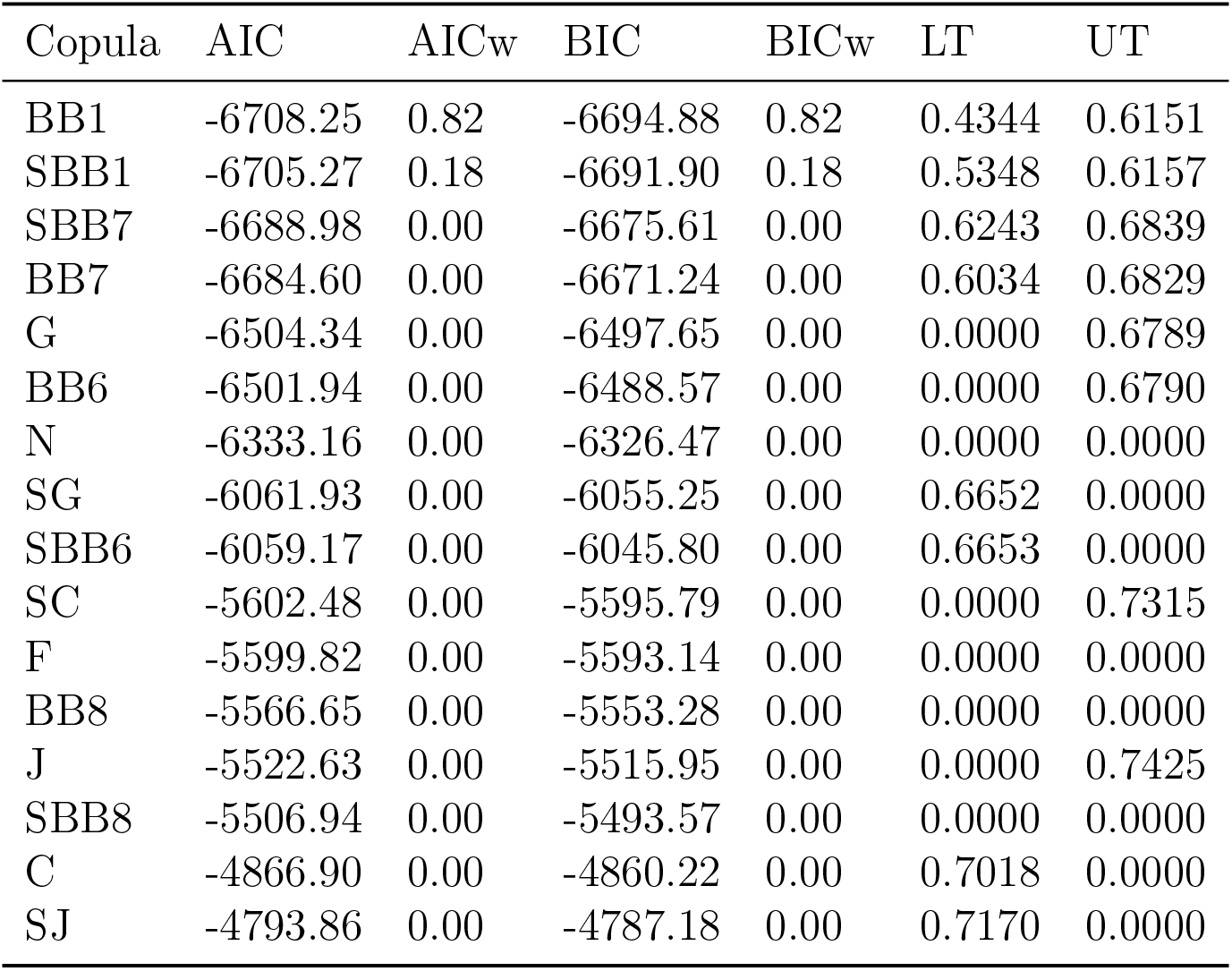
Model fitting results for soil C and N dataset using 16 copula families: normal (N), Frank (F), Clayton (C), survival Clayton (SC), Gumbel (G), survival Gumbel (SG), Joe (J), survival Joe (SJ), BB1, survival BB1 (SBB1), BB6, survival BB6 (SBB6), BB7, survival BB7 (SBB7), BB8, and survival BB8 (SBB8). Table rows are sorted by AIC. AICw = AIC weight; BICw = BIC weight; LT = the lower-tail dependence statistic for the indicated copula family with fitted parameters; UT = the same for upper-tail dependence.

Our model selection procedures do not reveal whether tail dependence parameters are *significantly* different from 0 and from each other. Furthermore, we caution that, while our model selection results do convincingly show non-normal copula structure, model-averaged tail-dependence statistics may have been biased because even the lowest-AIC copula (BB1) was a poor fit (*p* = 0 and 0.001 for the Cramer-von Mises and Kolmogorov-Smirnov goodness-of-fit tests, respectively, to within the precision available from `BiCopGofTest`). Our alternative nonparametric approach, results detailed next, provides information about significance of tail associations and asymmetries of tail associations.

The values of the statistics cor*_l_b_,u_b__*, P*_l_b_,u_b__* and 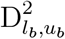 for the soil C and N data were compared to distributions of their values on 1000 Kendall- or Spearman-preserving normal surrogates of the data, separately in two comparisons for each of the ranges (*l_b_, u_b_*) = (0, 0.1), …, (0.9, 1) (Fig. 9). Results showed that tail associations of data are stronger in both the lower and upper tails than would be expected under a null hypothesis of a normal copula (Fig. 9B-F). The values of the statistics cor_0,0.1_ − cor_0.9,1_, P_0,0.1_ − P_0.9,1_ and 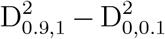 for the soil C and N data were also compared to distributions of their values on surrogates, separately in two comparisons (Table 3). Results showed that upper-tail association was significantly stronger than lower-tail association; i.e., C and N values are more related when high than when low. These results are consistent with the model selection results, but go beyond them by providing information about statistical significance. Thus our results provide an affirmative answer to Q1 for the soil C and N data; this answer is represented in Fig. 2 as a solid outline around “soil C and N” in the right-most box in the middle row.

**Figure 9:**
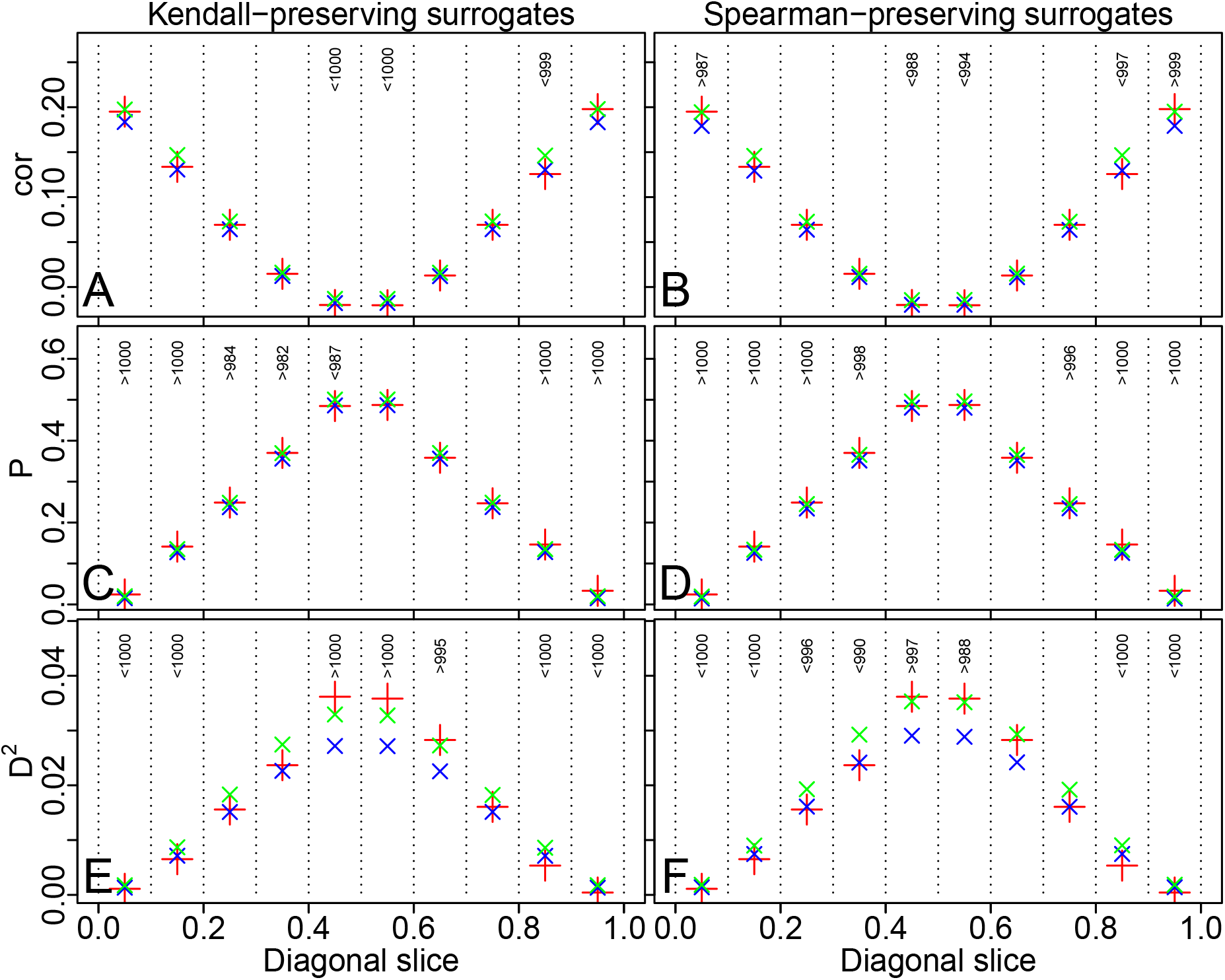
Nonparametric tests for tail association and other deviations from normal copula structure, for soil C and N data. As described in the main text, the values of the statistics cor*_l_b_,u_b__*, P*_l_b_,u_b__* and 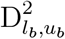 for real data (red crosses) were compared to distributions of their values on 1000 Kendall- or Spearman-preserving normal surrogates of the data (blue and green x’s show 0.025 and 0.975 quantiles), separately in two comparisons for each of the ranges (*l_b_, u_b_*) = (0,0.1), …, (0.9, 1). Whenever the red cross was outside the range given by the x’s, text at the top of panels indicates the number of surrogate values the real-data value was greater than or less than. For instance, a value > *N* (respectively, < *N*) means the value of the statistic on real data was greater than (respectively, less than) its value on *N* surrogates. When the statistic was greater than 975 or less than 975 surrogate values, it indicates significance (95% confidence level). When cor or P values (respectively, D^2^ values) were greater than surrogates, it means association in that part of the distributions was stronger than (respectively, weaker than) expected from a normal-copula null hypothesis.

**Table 3:** Soil C and N results for nonparametric tests of whether asymmetry of tail association was significant, compared to a normal-copula null hypothesis. The values of the listed statistics (cor_*FS*_-cor_*LS*_ = cor_0,0.1_ − cor_0.9,1_, P_*FS*_-P_*LS*_ = P_0,0.1_ − P_0.9,1_, 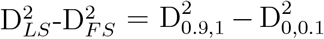; here FS stands for ‘first slice’ and LS for ‘last slice’) for real data were compared to their values for 1000 Kendall- or Spearman-preserving normal surrogates, in seperate comparisons. A table entry < *X* indicates the value of the given statistic on the data was less than its value on *X* of the surrogates, so entries of the form < *X* for *X* equal to 975 or above indicate that upper-tail dependence was significantly stronger than lower-tail dependence. Results were significant only for the *P* and *D* ^2^ statistics.

| Statistic | Kendall | Spearman |
| --- | --- | --- |
| $\text{cor}_{FS}-\text{cor}_{LS}$ | <695 | <696 |
| $P_{FS}-P_{LS}$ | <1000 | <1000 |
| $D_{LS}^2-D_{FS}^2$ | <1000 | <1000 |

Table 4 summarizes Q1 results for the bivariate datasets, with some details in the Appendices. The two variables were significantly related for all datasets (Table 4, row 1). A non-normal copula always emerged as the lowest-AIC copula (Table 4, row 2). The normal copula was a poor fit compared to the lowest-AIC copula (Table 4, rows 3-4), except for Cedar Creek, for which the AIC difference between these fits was marginal. Thus we answer the first part of Q1 in the affirmative for 3 of 4 of our bivariate datasets. Often, either lower- or upper-tail dependence statistics differed substantially from 0 (Table 4, rows 7-8), and/or these statistics differed from each other (Table 4, row 9), helping to answer the second and third parts of Q1. Though again the model selection results do not provide information on statistical significance of these differences, and are subject to the caveat that, for some datasets, even the lowest-AIC copula was not an objectively good fit (Table 4, rows 5 and 6), nonparametric results (Table 4, row 10) showed that tail associations deviated significantly from what would be expected from a normal-copula null model, and were also significantly asymmetric, except for the Cedar Creek data. See Figs S21-S23 for analogues to Fig. 9 for the bivariate datasets other than the soil C and N data, see Tables S3-S5 for analogues to Table 2, and see Table S6 for analogues to Table 3. The first three datasets had stronger upper- than lower-tail dependence, and the last dataset had the reverse, though non-significantly. Our generally affirmative (except for Cedar Creek), empirically based answer to Q1 for the bivariate datasets is represented in Fig. 2 as solid outlines around the names of the soil C and N and bird and mammal body mass versus metabolic rate datasets, in the middle row of boxes. We did the same analyses with Cedar Creek data from other available years, 1996-1999 and 2007. For 1996 and 2007, independence of biomass and Shannon’s index could not be rejected; for 1997-1999, results were similar to 2000.

**Table 4:** Summary of Q1 results for bivariate datasets. The *p*-values (rows 5-6) are for the minimum-AIC copula. Model averaging used for rows 7-9 was based on AIC weights. Row 10 shows values as in Table 3, upper right table entry. Although the result shown in row 10 for the soil C and N data is non-significant, see Table 3 for significant results using the *P* and *D*^2^ statistics. The first 3 datasets use (*l_b_, u_b_*) = (0, 0.1) for the first slice (FS) and (0.9, 1) for the last slice (LS), whereas (0, 0.25) and (0.75, 1) were used for Cedar Creek due to fewer data for that system. Results for Cedar Creek for the year 2000 are shown. See Table 2 for copula family abbreviations.

|  | Soil C<br>and N | Bird<br>masses<br>and<br>BMR | Mammal<br>masses<br>and<br>BMR | Cedar<br>Creek<br>data |
| --- | --- | --- | --- | --- |
| 1. p-value, independence test | 0 | 0 | 0 | 0 |
| 2. Minimum-AIC copula | BB1 | G | G | F |
| 3. AIC for best copula | -6708.25 | -1447.36 | -1854.31 | -22.68 |
| 4. AIC for normal copula | -6333.16 | -1336.77 | -1678.76 | -20.42 |
| 5. p-value, Cramer-von Mises goodness of fit test | 0 | 0.067 | 0.002 | 0.39 |
| 6. p-value, Kolmogorov-Smirnov goodness of fit test | 0.001 | 0.279 | 0 | 0.67 |
| 7. Model averaged lower-tail dependence | 0.4529 | 0 | 0 | 0.0551 |
| 8. Model averaged upper-tail dependence | 0.6152 | 0.877 | 0.893 | 0.0034 |
| 9. Model averaged lower- minus upper-tail dependence | -0.1623 | -0.877 | -0.893 | 0.0517 |
| 10. Placement of $\text{cor}_{FS}\text{-cor}_{LS}$ in surrogate distribution (see caption) | <696 | <999 | <993 | >897 |

We present green spruce aphid abundance (section 3; Table 1) results as an example multivariate analysis. Independence was rejected for each pair of the 10 sampling locations. Best-fitting (lowest-AIC) copulas were non-normal for the large majority of location pairs (72 of 90 pairs), and the AIC of the normal copula minus the minimum AIC, averaged across location pairs, was 2.732. Whereas 2.732 would be only a marginally meaningful AIC difference for a single location pair, it is more meaningful as an average across many pairs. Relatedly, and illustrating the concept here, the chances of getting 72 or more non-normal location pairs if the chances were equal of getting a normal versus a non-normal result (and taking into account that the location pair (A,B) will necessarily produce the same result as the pair (B,A)) is low, 3.287 × 10^−5^. The average AIC difference 2.732 was much less than the AIC difference between −6708.25 and −6333.16 that was obtained for the soil C and N data at least in part because the soil C and N data were much more numerous (Table 1). These results help answer the first part of Q1: non-normal copula structure appears to be meaningfully common in these data, even if not universal. Goodness of fit tests in every case failed to reject the hypothesis that the AIC-best copula family was also an objectively adequate description of the data; i.e., the collection of copula families we used was sufficiently broad to characterize these data. Model-averaged lower- and upper-tail dependence statistics had 2.5^*th*^ and 97.5^*th*^ quantiles (across location pairs) (0.123, 0.721) and (0.002, 0.41), respectively, thereby commonly differing from what a normal copula would give (i.e., 0), and helping to answer the second part of Q1: these data have greater tail dependence (lower and upper) than would be expected from a normal-copula null hypothesis. We note however that our model selection procedures again do not reveal whether tail-dependence parameters are *significantly* different from 0, and we refer the reader to nonparametric results below for information on significance. Model-averaged lower- minus upper-tail dependence statistics were positive for all but a few location pairs (86 out of 90). The chances of getting 86 or more positive values, here, under a null hypothesis of equal chances for positive and negative values (and again accounting for the fact that location pairs (A,B) and (B,A) will show the same result) was again low, 2.944 × 10^−11^. Thus the spatial synchrony of rarity in the green spruce aphid is stronger than the spatial synchrony of outbreaks. This answers the third part of Q1.

Nonparametric statistics verified that tail associations were asymmetric for the green spruce aphid abundance data. Mean values across all pairs of locations of the Spearman and Kendall correlations and the statistics cor_*l*_, cor_*u*_, P_*l*_, P_*u*_, 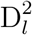, and 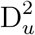 were positive and confidence intervals excluded 0 (Table 5). Mean values of the statistics cor_*l*_ – cor_*u*_, P_*l*_ – P_*u*_, and 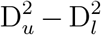 also were always positive and had confidence intervals that excluded 0 (Table 5).

**Table 5:** Average values of statistics across all locations pairs and confidence intervals based on spatial resampling for green spruce aphid abundance data.

|  | 2.5<sup>th</sup> quantile | Mean | 97.5<sup>th</sup> quantile |
| --- | --- | --- | --- |
| Spearman | 0.490 | 0.580 | 0.668 |
| Kendall | 0.351 | 0.428 | 0.505 |
| cor<sub>l</sub> | 0.291 | 0.335 | 0.375 |
| cor<sub>u</sub> | 0.195 | 0.250 | 0.307 |
| P<sub>l</sub> | 0.097 | 0.116 | 0.135 |
| P<sub>u</sub> | 0.063 | 0.084 | 0.106 |
| D<sub>l</sub><sup>2</sup> | 0.019 | 0.026 | 0.033 |
| D<sub>u</sub><sup>2</sup> | 0.030 | 0.039 | 0.047 |
| cor<sub>l</sub> - cor<sub>u</sub> | 0.034 | 0.085 | 0.128 |
| P<sub>l</sub> - P<sub>u</sub> | 0.015 | 0.032 | 0.048 |
| D<sub>u</sub><sup>2</sup> - D<sub>l</sub><sup>2</sup> | 0.005 | 0.013 | 0.020 |

Table 6 summarizes Q1 results for multivariate data. Results supported the conclusions that non-normal copula structure, non-normal tail dependence, and asymmetric tail associations were common, though not universal, answering Q1 in the affirmative for these data. Most location pairs were non-independent (Table 6, row 2). The large majority of non-independent location pairs had best-fitting copulas that were not the normal copula (Table 6, row 3), and AIC values of best-fitting copulas were, on average across location pairs, between 2.714 (*Ceratium furca* abundance data) and 5.764 (leaf-curling plum aphid first-flight data) lower than AIC values for the normal copula (Table 6, row 4). Best-fitting copulas were nearly always considered an adequate fit (Table 6, row 5). Some of the datasets (green spruce aphid abundance, *Ceratium furca* abundance, methane-flux) showed stronger lower- than upper-tail dependence (Table 6, rows 6-8), whereas leaf-curling plum aphid first flight data showed the reverse. All of the values in Table 6, rows 8 deviated highly significantly from what would have been expected under a null hypothesis of equal chances for positive and negative values.

**Table 6:** Summary of Q1 results for multivariate datasets. Rows 3-8 of the table were computed for the non-independent pairs (row 2) only. A well-fitting location pair (row 5) was one for which the best-fitting copula had *p*-values > 0.01 for both the Cramer-von Mises and Kolmogorov-Smirnov goodness of fit tests.

|  | Green spruce<br>aphid<br>abundance | Leaf-curling<br>plum aphid<br>first flight | *Ceratium*<br>*furca*<br>abundance | Methane-flux |
| --- | --- | --- | --- | --- |
| 1. Location pairs (excluding self comparisons) | 90 | 90 | 182 | 156 |
| 2. Number of non-independent pairs | 90 | 90 | 162 | 102 |
| 3. Number of pairs with non-normal copula as best fit | 72 | 82 | 134 | 96 |
| 4. Average of AIC differences (normal AIC minus min AIC) across location pairs | 2.732 | 5.764 | 2.714 | 4.482 |
| 5. Number of well-fitting location pairs | 90 | 83 | 160 | 102 |
| 6. 2.5<sup>th</sup> and 97.5<sup>th</sup> quantiles of model-avg. lower-tail dependence | (0.123, 0.721) | (0, 0.152) | (0.019, 0.629) | (0.004, 0.461) |
| 7. 2.5<sup>th</sup> and 97.5<sup>th</sup> quantiles of model-avg. upper-tail dependence | (0.002, 0.41) | (0.175, 0.798) | (0, 0.505) | (0, 0.592) |
| 8. Percent pairs showing stronger lower tail dependence | 95.56 | 0 | 79.01 | 72.55 |
| 9. Mean with 2.5<sup>th</sup> and 97.5<sup>th</sup> quantiles of $\text{cor}_l - \text{cor}_u$ across all location pair | 0.085, (0.034, 0.128) | -0.15, (-0.207, -0.096) | 0.077, (0.031, 0.118) | 0.007, (-0.039, 0.05) |

Asymmetry results were generally verified by nonparametric approaches, with the exception of the methane-flux data. For instance, the 95% confidence intervals of the mean over pairs of sampling locations of the statistic cor_*l*_ – cor_*u*_ were (−0.207, −0.096) for the leaf-curling plum aphid first flight data (Table 6, row 9), indicating greater upper-tail dependence, and consistent with the results of Table 6, row 8. For the *Ceratium furca* abundance data, confidence intervals were (0.031, 0.118), indicating greater lower-tail dependence, and again consistent with the results of Table 6, row 8. For the methane-flux data, confidence intervals were (−0.039, 0.05); the asymmetry in tail dependence revealed by the model selection results for the methane-flux data was apparently not strong enough to also be revealed by the nonparametric analyses. Analogues to Table 5 are in Tables S7-S9. Our generally affirmative, empirically based answer to Q1 for the multivariate datasets is represented in Fig. 2 as solid outlines around the names of those datasets, in the middle row of boxes. We also carried out the same analyses for abundance and first-flight data for the 18 aphid species for which we had data other than the green spruce and leaf-curling plum aphids, as well as for the 21 plankton taxa for which we had abundance data other than *Ceratium furca* (results not shown). Results supported the conclusion that non-normal copula structure and tail dependence, and asymmetry of tail associations, are common.

## 6 Concepts and methods for Q2

Having demonstrated that non-normal copula structure and asymmetric tail associations are addressed Q2 by exploring, using models, three possible mechanisms by which these phenomena may arise. Models presented are initial explorations, only, of whether the proposed mechanisms have the potential to explain observed patterns. As such, simple models were used. Comprehensive explorations of model parameter space and alternative model formulations were left for future work.

The first mechanism relates to the ideas in the Introduction about Liebig’s law of the minimum, and to nonlinear influences of environmental variables on ecological variables. If an environmental variable influences an ecological variable disproportionately in one of its tails, we explored whether the ecological variable could then exhibit asymmetric tail associations across space. Let *E_i_*(*t*) be an ecological variable measured at location *i* (*i* = 1, 2) at time *t.* Assume the dynamics *E_i_*(*t* + 1) = *bE_i_*(*t*) + *g*(*ϵ_i_*(*t*)) + *aδ_i_*(*t*), where the *δ_i_*(*t*) are standard-normally distributed and independent across time and locations, *a* = 0.2, *b* = 0.1, –0.1,0.5, −0.5 in different simulations, and the (*ϵ*_1_(*t*), *ϵ*_2_(*t*)) were drawn, independently through time, from a bivariate normal distribution with var(*ϵ_i_*) = 1 and cov(*ϵ*_1_, *ϵ*_2_) = 0.8. Thus basic ecological dynamics follow the very simple AR(1) formulation, influenced by a “regional” environmental factor which is correlated across locations (*ϵ*) and by an entirely local factor (*δ*). The function *g* describes how the environmental factor *ϵ_i_*(*t*) influences *E_i_*(*t*). We used *g* equal to *g*_1_ or *g*_2_ in different simulations: *g*_1_(*ϵ*) equals *ϵ* if *ϵ* < 0 and 0 otherwise; *g*_2_(*ϵ*) equals *ϵ* if *ϵ* > 0 and 0 otherwise. Thus *g*_1_ represents environmental effects that negatively impact populations, and that occur only below a threshold of *ϵ* = 0; and *g*_2_ represents effects that positively impact populations and that occur only above a threshold of *ϵ* = 0. The values of *b* selected provide a modicum of exploration of the potential for ecological dynamics to also influence how complex dependencies between the *E_i_*(*t*) may arise: negative values of *b* correspond to overcompensating dynamics and positive values to undercompensating dynamics; larger |*b*| means slower return to equilibrium after a disturbance.

For each *b* and *g_i_*, we simulated the model for 25000 time steps and retained the *E_i_*(*t*) for the final 2500 time steps. We applied our nonparametric statistic cor_0,0.1_ – cor_0.9,1_ and our Spearman- or Kendall-preserving normal-copula surrogate comparison methods (see section 4) to these outputs to discover if the model could produce asymmetric tail associations (and therefore non-normal copula structure) between the *E_i_*(*t*). Because (*ϵ*_1_, *ϵ*_2_) and (*δ*_1_, *δ*_2_) have normal copula structure (they were drawn from bivariate normal distributions), the Moran mechanism analyzed below does not operate here.

Our second mechanism is an extension of the well known Moran effect, and was summarized conceptually in the Introduction. We consider a linear model, as well as two parameterizations of a nonlinear population model which includes density dependence of population growth. For the linear model, let *E_i_*(*t*) again be an ecological variable, *i* = 1, 2. We use AR(1) dynamics, 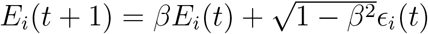, with *β* = 0.5. The environmental noises *ϵ_i_*(*t*) were standard-normal random variables that were independent for distinct times, *t*, but exhibited different kinds of dependence across locations in different simulations (see below). The variable *E* is very general, and could represent deviations of a population density from a carrying capacity, deviations of total plant community biomass from an average value, flux of a biogeochemical variable such as methane, or other quantities. The nonlinear population model was a stochastic, multi-habitat-patch version of the Ricker model, *P_i_*(*t* + 1) = *P_i_*(*t*) exp [*r* (1 – *P_i_*(*t*)/*K*) + *σϵ_i_*(*t*)] for *i* =1, 2, using *r* = 0.5, *K* = 100, *ϵ_i_*(*t*) as above, and *σ* = 0.1 or *σ* = 1 in different simulations. When *σ* = 0.1, population dynamics stay close to the carrying capacity, *K*, where the model equation can be well approximated by a linear equation, and the nonlinearities of the model therefore have limited influence on dynamics. When *σ* = 1, model dynamics are strongly nonlinear because the stochastic component of the model causes populations to stray far from the carrying capacity, *K*. We refer to these as the weak-noise and strong-noise cases, though the importance of the noise here is that, when it is strong, it brings the nonlinearities of the model into play.

For each of the three model setups above (the linear model and the nonlinear model with weak and strong stochasticity), for the Clayton and survival Clayton copula families, for each *τ* = 0.1, 0.2,…, 0.9, and for each of 50 replicate simulations, we generated 5000 noise pairs (*ϵ*_1_(*t*), *ϵ*_2_(*t*)) from the bivariate random variable with standard-normal marginals and with the given copula family and the given Kendall correlation *τ*. We then used this noise to drive the model, and retained both the noise and population values for the final 500 time steps. For each simulation, the following statistics were then computed for both noises and populations: Pearson, Spearman and Kendall correlations, cor_*l*_, cor_*u*_, P_*l*_, P_*u*_, 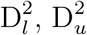, cor_*l*_ – cor_*u*_, P_*l*_ – P_*u*_, and 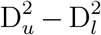. Values were plotted against *τ* for noises and populations. If the hypothesis from the Introduction was reasonable that characteristics of the copula structure of spatial dependence in an ecological variable may be inherited from characteristics of spatial dependence in an environmental variable through a Moran-like effect, then plots should be similar for populations and noises.

The next mechanism we investigated is evolutionary, and pertains to bivariate trait data across species, e.g., our bird and mammal data. This mechanism is a hypothetical explanation for the bias toward right tail association observed in those data (Fig. 8F,G). The hypothesis is that asymmetric tail association occurs in evolutionary changes in bivariate characters, and gives rise to asymmtric tail association between the two character values across extant species. We simulated bivariate character evolution on an estimate of the phylogeny, taken from Genoud *et al.* (2018), of 817 mammal species. The root character state and the change across each branch were randomly chosen from matrices of one million independent draws from bivariate distributions showing one of five distinct types of copula structure: 1) extreme or 2) moderate left-tail dependence, 3) symmetric tail dependence, or 4) moderate or 5) extreme right-tail dependence (Appendix S8). All distributions had standard-normal marginals and Spearman correlation 0.875 between components, so our simulations assess the impact of copula structure only. For each of the five copulas, mammalian character evolution was simulated 100 times. For each simulation, symmetry of tail associations of the two characters across phylogeny tips was assessed using our nonparametric statistics. We hypothesized that cases 1 and 2 above would yield stronger left- than right-tail associations in tip characters, and cases 4 and 5 would yield the reverse. The simulator was written in Python and used version 4.4 of the DendroPy package (Sukumaran & Holder 2010).

## 7 Results for Q2: Moran effects and asymmetric dependencies produce non-normal copula structure

Our model with asymmetric environmental effects produced outputs with visually apparent asymmetry of tail associations between the ecological variables *E_i_* in the two locations, to an extent that depended on the value of *b*. For *b* = 0.1 (Fig. 10) and *b* = –0.1 (Fig. S24A,B), for both *g*_1_ and *g*_2_, asymmetry of tail association was strong; for *b* = ±0.5, asymmetry was weaker but still apparent (Fig. S24C-F). Lower-tail (respectively, upper-tail) spatial associations in the effects of noise (*g*_1_, respectively, *g*_2_) produced lower-tail (respectively, upper-tail) spatial associations in the ecological variable, *E_i_*. Results using our statistic cor_0,0.1_ – cor_0.9,1_ strongly reflected the visually apparent asymmetry (Table 7). Thus asymmetry of environmental effects is a mechanism that may be partly responsible for non-normal copula structure and asymmetric tail associations across space in ecological variables. This result is represented in Fig. 2 as the solid box around “Nonlinear environmental effects, Liebig’s law” and the arrows labelled “A”. It is explained in the Discussion why the box and some of the arrows are solid, instead of dashed, although the results of this section are theoretical.

**Figure 10:**
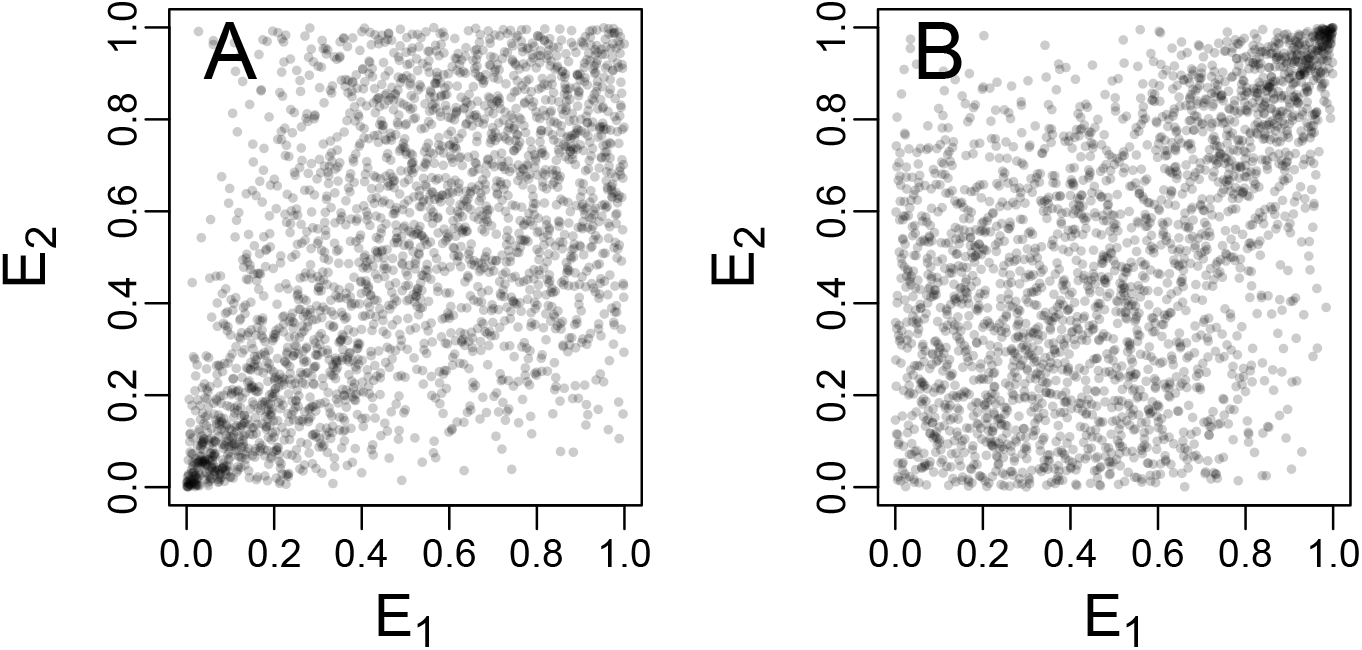
If environmental effects operate asymmetrically in their tails on ecological variables, it can result in non-normal copula structure and asymmetric tail dependence across space in the ecological variables. Shown are the last 500 points for (A) *g* = *g*_1_ and (B) *g* = *g*_2_ from simulations described in the text, *b* = 0.1. Asymmetric tail associations are visually apparent, but were also elaborated statistically in Table 7.

**Table 7:** Asymmetric sensitivity model, nonparametric statistics results. Outputs of simulations from the first model detailed in section 6 (i.e., Figs 10 and S24) were subjected to nonparamtric statistical analyses decribed in section 4. The statistic cor_*FS*_ – cor_*LS*_ was computed for the output of the model with the indicated b and g, and the value was compared to 1000 values of the same statistic computed on surrogate time series randomized to have normal copula structure (see section 4). A table entry < *X* (respectively, > *X*) indicates the value of the given statistic on the data was less (respectively, more) than its value on *X* of the surrogates, so entries of the form < *X* (respectively, > *X*) for *X* equal to 975 or above indicate that upper-tail (respectively, lower-tail) dependence was significantly stronger than lower-tail (respectively, upper-tail) associations compared to a normal-copula null hypothesis. ‘FS’ stands for ‘first slice’ and refers to the bounds 0 to 0.1; ‘LS’ stands for ‘last slice’ and refers to the bounds 0.9 to 1.

| $b$ | $g$ | Kendall | Spearman |
| --- | --- | --- | --- |
| 0.1 | $g_1$ | $>1000$ | $>1000$ |
| 0.1 | $g_2$ | $<1000$ | $<1000$ |
| -0.1 | $g_1$ | $>1000$ | $>1000$ |
| -0.1 | $g_2$ | $<1000$ | $<1000$ |
| 0.5 | $g_1$ | $>1000$ | $>1000$ |
| 0.5 | $g_2$ | $<1000$ | $<1000$ |
| -0.5 | $g_1$ | $>1000$ | $>1000$ |
| -0.5 | $g_2$ | $<1000$ | $<1000$ |

The hypothesis that characteristics of the copula structure of spatial dependence in an ecological variable can be inherited from an environmental variable was found to be reasonable, because it held for the models we examined; however, similarities between environmental- and ecological-variable copula structure were reduced when dynamics were strongly nonlinear. For simulations using AR(1) models, our correlation and tail asymmetry statistics were always similar for noise and populations (Fig. 11 for the Clayton copula, Fig. S25 for the survival Clayton copula). Though there were significant differences for many statistics and simulations, these were small compared to the overall tendency for larger (respectively, smaller) values of our statistics, as applied to noise, to be paired with larger (respectively, smaller) values of the same statistics as applied to model outputs. For our nonlinear model with weak noise, values of the statistics were again quite similar for noise and model outputs (Fig. S26 for Clayton, Fig. S27 for survival Clayton). Though there were again significant differences for many statistics and simulations, these were again small relative to overall variation of values of the statistics. Since many ecological models are nonlinear, this result provides the reasonable expectation that there may be a Moran-effect-like correspondence between noise and model-output dependence structure across space for typical ecological dynamics, as long as environmental noise is small enough that dynamics stay relatively close to the model equilibrium. Theoretical results that hold for “weak noise” in this sense are common in ecology. For strong noise and using our nonlinear model, our correlation and tail-association-asymmetry statistics, generally speaking, were approximately similar between noise and model outputs; however, similarity was reduced relative to previous results, and for a few simulations, asymmetry statistics had opposite signs for noise and model outputs (Figs S28 and S29). For instance, using a Clayton copula with a large Kendall *τ*, cor_*l*_ – cor_*u*_ was slightly positive for noise, but slightly negative for population model outputs (Fig. S28).

**Figure 11:**
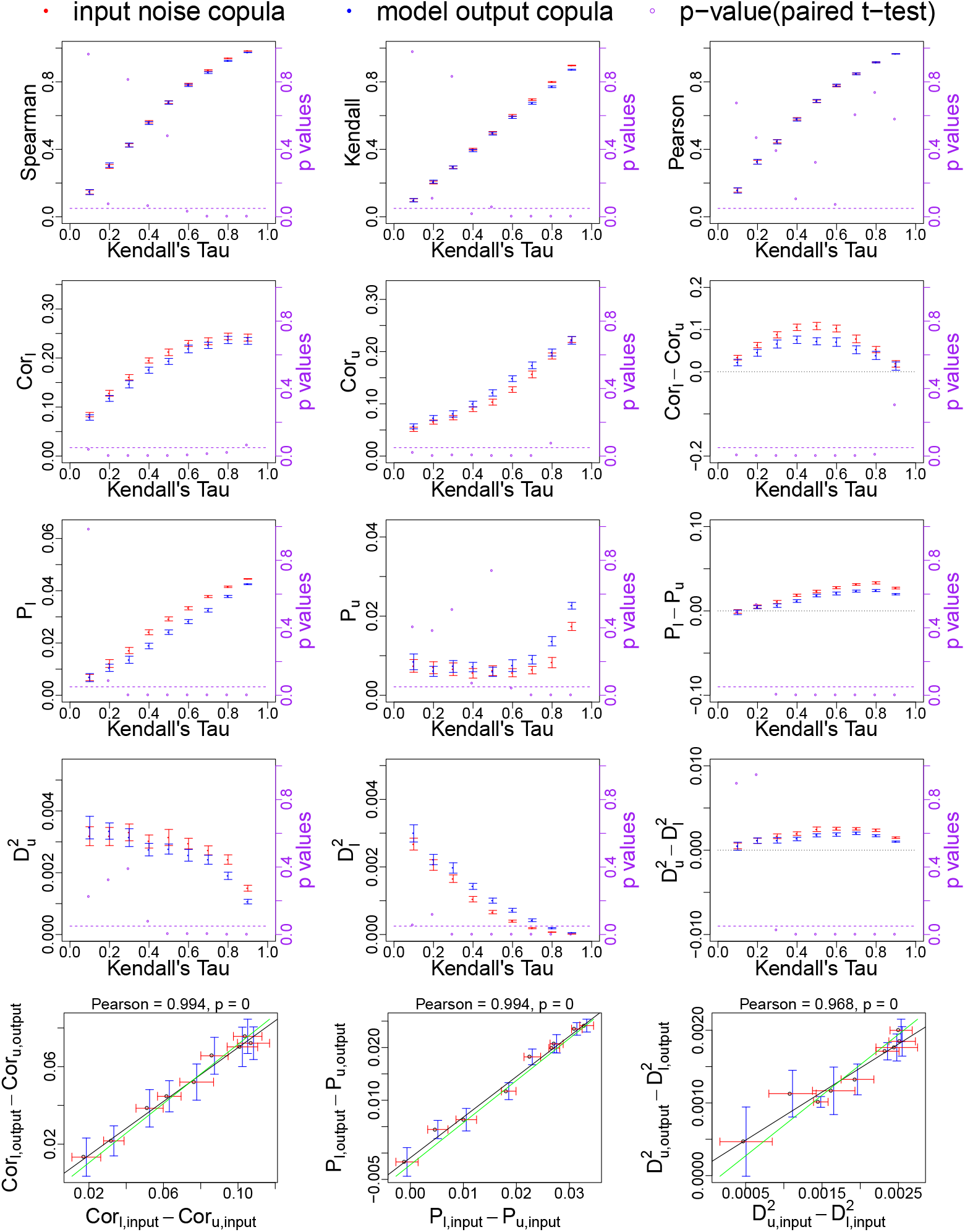
Comparison of correlation and tail association statistics between environmental-noise inputs and ecological-variable outputs, AR(1) model, Clayton copula. Red and blue points give means over 50 replicate simulations of the listed statistics, red points are for noise inputs and blue points are for model outputs. Error bars are standard errors. *P*-values (purple points, right axis) are for a paired *t*-test of the null hypothesis that the distributions have the same mean. For bottom panels, headers give Pearson correlations for the points. The regression line through the points (black) was similar to the 1-1 line (green). See section 6 for details.

We repeated our analyses using the nonlinear model with *r* = 1.3. The deterministic one-habitat-patch Ricker model exhibits a monotonic approach to a stable equilibrium at *K* when *r* < 1 (undercompensating dynamics, e.g., the value *r* = 0.5 used initially), but exhibits an oscillatory approach to *K* when *r* > 1 (overcompensating dynamics, e.g., *r* = 1.3). For weak noise, similarities were again dominant between values of our statistics on noise and population time series. For strong noise, however, discrepancies were often glaring. Apparently noise of standard deviation 1 interacted especially strongly with model nonlinearities when the model was in its overcompensatory regime. We repeated all analyses using the Gumbel, survival Gumbel, Joe, and survival Joe copulas, with substantially similar main conclusions (not shown).

Thus it is reasonable to hypothesize that a Moran-effect-like mechanism may produce non-normal copula structure and asymmetric tail associations across space in ecological variables. This result is represented in Fig. 2 as the dashed box around “Moran effects” and the arrows labelled “B”.

For our evolutionary model, the hypothesis was correct, for the models we employed, that asymmetric tail association in evolutionary changes can produce similarly asymmetric tail association between characters across phylogeny tips. Once character evolution was simulated 100 times for each of the dependence structures we considered for evolutionary changes, we had 817 bivariate characters for each of 500 simulations (see section 6). We computed our nonparametric asymmetry statistics cor_0,0.2_ – cor_0.8,1_, P_0,0.2_ – P_0.8,1_, and 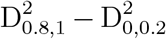 for each simulation output and produced a histogram for each statistic and for each dependence structure (Fig. 12). Results showed that asymmetries of tail associations in evolutionary changes were associated with similar asymmetries of tail associations in extant characters. Thus, based on our simulations, we cannot reject the possibility that this is a proximate mechanism behind observed asymmetric tail associations in our bird and mammal data. This result is represented in Fig. 2 as the dashed box around “If character evolution has copula structure” and the arrow labelled “D”. This topic is revisited in the Discussion, as is the box “Asymmetric species interactions” and the arrow labelled “C” in Fig. 2.

**Figure 12:**
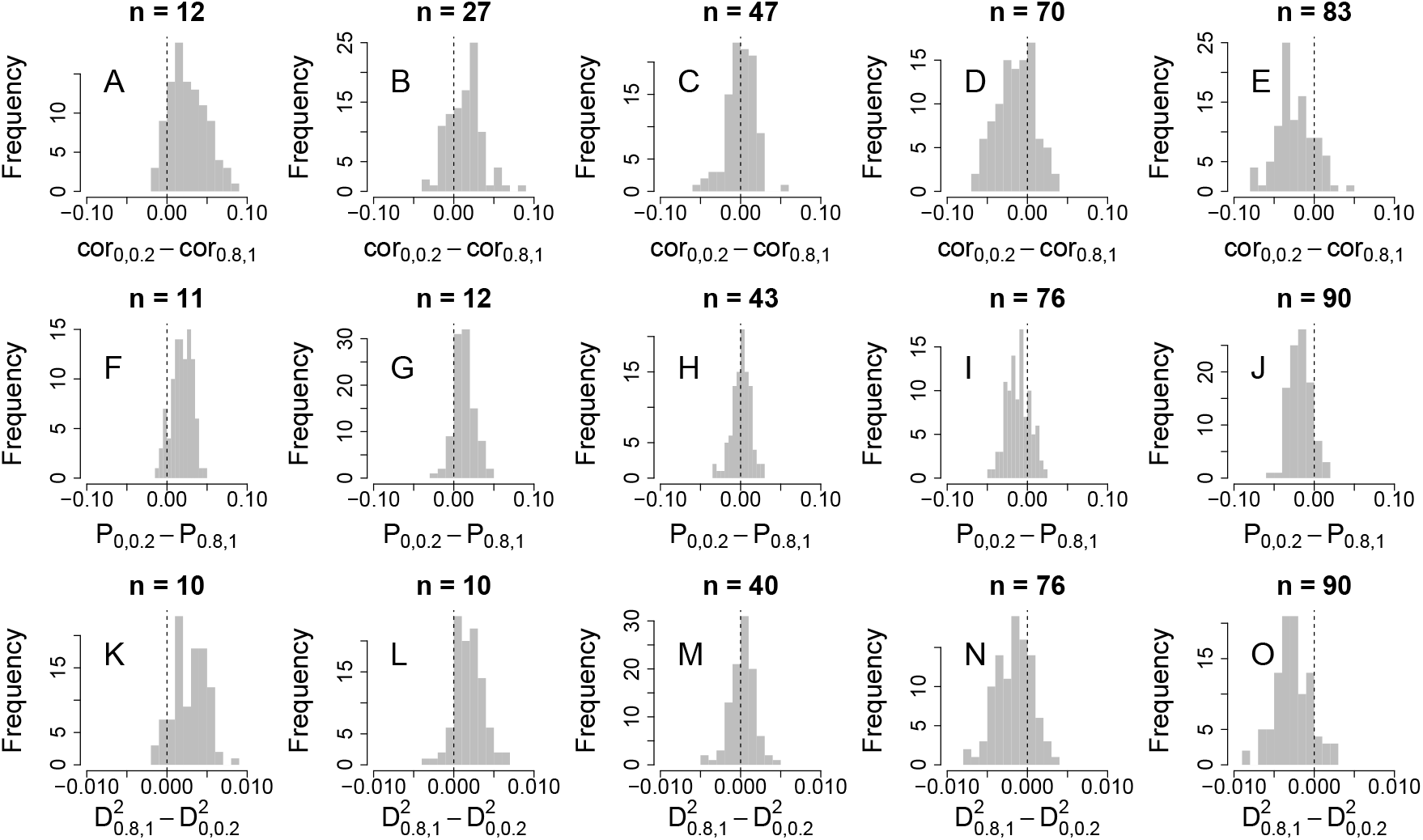
Three measures of asymmetry of tail associations (cor_0,0.2_ − cor_0.8,1_, A-E; P_0,0.2_ − P_0.8,1_, F-J; 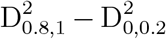, K-O) between two characters, across extant species, were computed for each of 100 simulations of mammalian character evolution for each of five types of tail dependence between evolutionary changes in the characters (extreme left-tail dependence, A, F, K; moderate left-tail dependence, B, G, L; symmetry of tail dependence, C, H, M; moderate right-tail dependence, D, I, N; and extreme right-tail dependence, E, J, O). See section 6 for details. The number above each panel indicates how many statistics out of 100 were less than or equal to 0. Values substantially lower (respectively, higher) than 50 indicate greater left (respectively, right) tail-dependence between characters.

## 8 Concepts and methods for Q3

We addressed Q3 by exploring, using both data and models, three hypothesized ecological consequences of non-normal copula structure and asymmetric tail associations. As for Q2, the models we used are intended as initial explorations, only, of whether copula structure may have a meaningful influence on important ecological phenomena. Therefore only simple models are used, and comprehensive explorations of the sensitivity of results to model strucuture and parameters are left for future work.

The hypothesis was presented in the Introduction that the distribution (through time) of a spatially averaged quantity should be influenced by dependencies between the local quantities being averaged, including their copula structure and tail associations. To make this hypothesis more precise, suppose an ecological variable *E_i_*(*t*) is measured at locations *i* = 1,…, *N* and times *t* = 1,…,*T*, and the spatial mean ∑*_i_E_i_*(*t*)/*N* is of interest. The *E_i_*(*t*) could be, for instance, local abundances of a pest or exploited species, or local fluxes of a greenhouse gas. If *E_i_* and *E_j_* are associated primarily in their right tails for most location pairs *i* and *j*, then exceptionally large values tend to occur at the same time in most locations. We hypothesize that this will tend to increase the skewness of the distribution of the spatial mean. Similarly, left-tail associations between local variables should tend to decrease skewness. Strong positive skewness of the spatial-mean time series corresponds to “spikiness” of that time series, i.e., occassional very large values, which corresponds to instability through time. The spatial-mean time series and its skewness may be quantities of principal importance for pest or resource abundance, for which extreme values (spikes) in the spatial mean may have large effects.

We tested the above hypothesis using our multivariate datasets (Table 1). For each dataset we calculated the spatial mean time series, and then the skewness through time of that mean. Then, for each dataset, we compared the value obtained to a distribution of values of the same quantity for each of 10000 surrogate datasets. Surrogate datasets were produced by randomizing the empirical data in a special way to have the copula structure of a multivariate-normal distribution, but to retain exactly the same distributions of values for each sampling location as the original data (Appendix S9). Surrogates also had very similar Spearman correlations between pairs of sampling locations as the data. Our comparisons therefore tested the null hypothesis that the skewness of the spatial mean took values on the empirical data no different from what one would expect if the copula structure of the data were multivariate normal (i.e., the same copula as a multivariate normal distribution), but the data were otherwise statistically unchanged. Significant differences indicate that non-normal copula structure in the data contributed to the skewness of the spatial mean time series, i.e., to its instability and “spikiness” through time.

For green spruce aphid abundance data, *C. furca* abundance data, and methane-flux data, because these datasets exhibited stronger lower- than upper-tail associations (Table 6), we compared empirical and surrogate skewness values via a one-tailed test in the left tail: the *p*-value was the fraction of surrogate skewnesses less than the skewness for the empirical data. The test examines whether stronger lower-tail associations between local time series caused the spatial average to have significantly lower skewness than would have been expected with symmetric tail associations. For leaf-curling plum aphid data, because that dataset exhibited stronger upper- than lower-tail associations (Table 6), we did the analogous one-tailed test in the right tail. The test examines whether upper-tail associations caused the spatial average to have significantly higher skewness than would have been expected with symmetric tail associations.

To further address Q3, we also examined a hypothesis that asymmetric tail associations across space of an environmental variable can influence the extinction risk of a metapopulation. We hypothesized that environmental noises exhibiting greater left-tail associations across different habitat patches would cause higher metapopulation extinction risks because then very bad years for the component populations would occur simultaneously in many patches, reducing rescue effects. Here we assume, for simplicity, that low values of the environmental variable are “bad” for the populations and high values are “good”. We tested the reasonableness of this hypothesis using a metapopulation extension of the Lewontin-Cohen model, 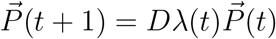, where the *i^th^* component of the length-*N* vector 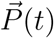 represents population density in the *i^th^* habitat patch at time *t*. The *N* × *N* matrix *λ*(*t*) was diagonal with *i^th^* diagonal entry exp(*r* + *ϵ_i_*(*t*)). Here *r* is a growth rate; we used *r* = 0. The *ϵ_i_*(*t*) represent environmental noises. They were standard-normally distributed, were independent through time, and showed the same spatial correlations for every simulation, but were made to exhibit stronger right- or left-tail associations between patches in different simulations (Appendix S10). The *N* × *N* matrix *D* was a dispersal matrix modelling local or global dispersal at rate *d*, in different simulations (Appendix S10). After each step, if the density in a patch was < 1, it was set to 0. We simulated the model 10000 times for each combination of parameters, starting from *p*_0_ = 50 in each patch, and calculated extinction risk after 25 time steps.

Finally, and pursuing ideas that sprang from earlier theoretical work (Reuman *et al.* 2017), we tested whether the copula structure of the dependences between population variables measured in different locations has consequences for the spatial version of Taylor’s law. Taylor’s law is a commonly observed and widely studied (Taylor 1961; Taylor *et al.* 1988; Cohen *et al.* 2013; Xu *et al.* 2016; Reuman *et al.* 2017) empirical pattern that relates the variances of groups, *g*, of population measurements to the means of the groups via a power law, 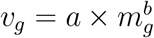, or equivalently, log(*v_g_*) = log(*a*) + *b* × log(*m_g_*). Here *b* is called the Taylor’s law *exponent* or *slope*, and log(*a*) is the Taylor’s law intercept. There are several versions of Taylor’s law. For spatial Taylor’s law, given population density or abundance data *x_i_*(*t*) measured in locations *i* = 1, …, *N* and times *t* = 1, …, *T*, a group *g* consists of all the measurements *x_i_*(*t*), *i* = 1, …, *N* made in different locations at the same time. So means *m_g_* and variances *v_g_* are computed across space. We refer to the matrix with *x_i_*(*t*) in the *t*^th^ row and the *i*^th^ column as a *population matrix*. We consider that Taylor’s law holds true for a dataset if the log(*v_g_*) versus log(*m_g_*) scatter plot for the dataset shows a linear, homoskedastic pattern. Taylor’s law has been verified empirically for hundreds of taxa and has been applied in numerous fields including fisheries management, estimation of species persistence times, and agriculture (Cohen & Xu 2015; Reuman *et al.* 2017), so the question of whether copula structure of spatial dependence influences Taylor’s law may have applied significance.

To explore the influence of copula structure and tail associations on spatial Taylor’s law, we carried out a series of simulations that generated sets of population matrices that had distinct copula structure between locations but that were otherwise statistically similar. For *k* = 1, …, 1000 and *X* representing a Clayton, normal, Frank, or a survival Clayton copula family, we generated *N* × *n* population matrices *m*^(*X,k*)^ with the following properties: 1) for any given values of *j* and *k*, the values 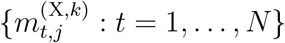 were exactly the same, as unordered sets, for all *X*, i.e., the same actual population values were used for location *j* in simulation *k*, regardless of *X*; 2) for any given *k* and *j*_1_ and *j*_2_ such that *j*_1_ ≠ *j*_2_, the Spearman correlations (computed through time) 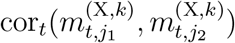 were the same, to within sampling variation, i.e., correlations through time of populations in two locations were the same, up to sampling variation, regardless of *X*; 3) the copula for the dependence between 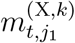 and 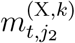 was from the family *X*. We used *N* = 50 and *n* = 25. Details of how these matrices were generated are in Appendices S11 and S12. Thus for each *k*, the poulation matrices *m*^(*X,k*)^ for *X* taking the values Clayton, normal, Frank, and survival Clayton were statistically similar except for the copula structure between locations, which was *X*. Thus comparing how the *log*(*v_g_*) versus log(*m_g_*) relationship may manifest differently for *m*^(*X,k*)^ for different values of *X* constitutes a test of the influence of copula structure on Taylor’s law. We selected *X* equal to the Clayton, normal, Frank, and survival Clayton copulas to explore a range of tail association patterns.

A variety of Taylor’s law statistics were computed for each populaton matrix *m*^(*X,k*)^. First, the *N* = 50 spatial means, *m_g_*, and variances, *v_g_*, were computed, and log(*v_g_*) vs. log(*m_g_*) plots were considered. Linearity of the log(*v_g_*) vs. log(*m_g_*) relationship was tested for each simulation by comparing the linear regression through these 50 points to a quadratic alternative via an *F*-test, producing a *p*-value result. If these *p*-values were uniformly distributed across the unit interval for the 1000 replicate simulations which were generated for *X*, it supported the linearity assumption of Taylor’s law for *X*; whereas if they were clustered toward smaller values the test tended to reject that assumption. We also tested, for each simulation, the assumption of Taylor’s law that the log(*v_g_*) vs. log(*m_g_*) plot was homoskedastic: we regressed the absolute residuals of the linear regression of log(*v_g_*) versus log(*m_g_*) against the predictions of that regression. A significant *p*-value result of this test indicates heteroskedasticity. For each simulation we also computed the root mean squared error of log(*v_g_*) vs. log(*m_g_*) data from the linear regression of log(*v_g_*) versus log(*m_g_*), as well as the intercept and slope of the linear regression. Finally we recorded the quadratic coefficient of the regression of log(*v_g_*) against log(*m_g_*) and (log(*m_g_*))^2^, and the mean curvature of the quadratic regression equation across the values log(*m_g_*). The distrubutions of all these statistics across the 1000 replicate simulations were compared for *X* the Clayton, normal, Frank, and survival Clayton copulas, to determine if copula structure influences Taylor’s law.

## 9 Results for Q3: Tail dependence influences skewness of the spatial average, extinction risk, and Taylor’s law

Empirical results were consistent with the hypothesis that the skewness of a spatial-average time series is influenced by tail associations between the local quantities being averaged. For datasets that exhibited highly asymmetric tail associations in earlier analyses (green spruce aphid abundance data had stronger lower-tail association, and, respectively, leaf-curling plum aphid first flight data had stronger upper-tail association, Table 6), skewness of the spatial average was less than (respectively, greater than) a significant fraction of surrogate skewness values (Fig. 13A,B). For datasets with moderately stronger lower than upper-tail dependence (*Ceratium furca* abundance and methane data), skewness of the spatial average showed a non-significant or marginally significant tendency toward being less than surrogate skewnesses (Fig. 13C,D). Thus copula structure and asymmetric tail associations are important for spatially averaged quantities. This result is represented in Fig. 2 as the solid box around “Instability/skewness of mean or total time series” and the solid arrows labeled “X”.

**Figure 13:**
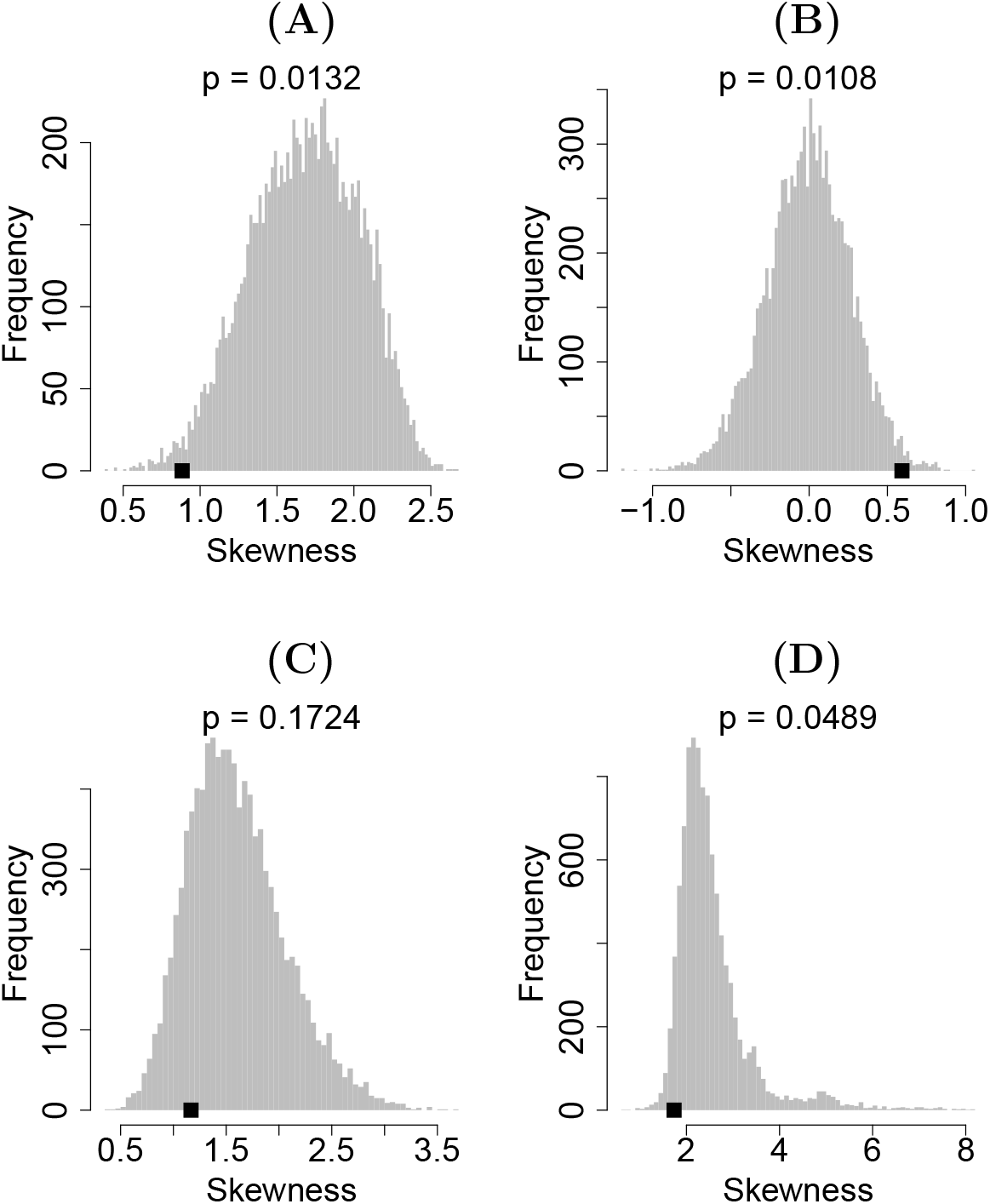
Skewness of spatially averaged green spruce aphid abundance (A), leaf-curling plum aphid first flight dates (B), *Ceratium furca* abundance (C), and methane-flux (D) time series compared to a multivariate normal-copula null hypothesis. Black dots are empirical skewnesses; see text and Appendix S9 for details of the null hypothesis. Results show a tendency for skewness of the spatial average to be affected as hypothesized by asymmetric tail associations.

Consistent with our extinction risk hypothesis, left-tail-associated environmental fluctuations increased metapopulation extinction risk for the spatial Lewontin-Cohen model, for *N* = 5 and *N* = 25 habitat patches, and for local and global dispersal (Fig. 14). This result is represented in Fig. 2 as the dashed box around “Extinction risk” and the dashed arrow labelled “Y”.

**Figure 14:**
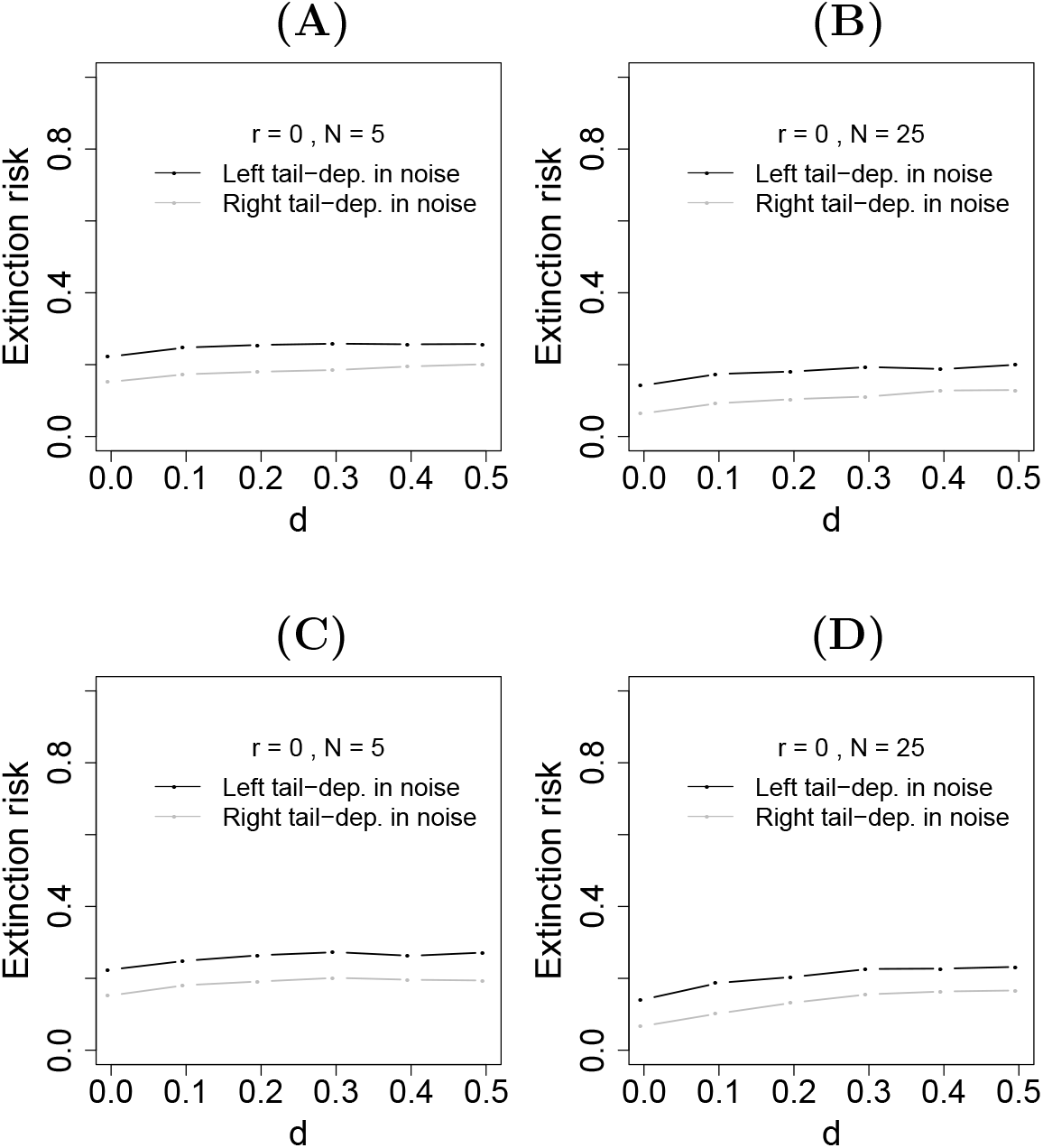
Extinction risk for the metapopulation extension of the Lewontin-Cohen model, after 25 time steps, was higher for environmental noise with stronger left-tail association across space. Dispersal was local (A, B) or global (C, D), with dispersal rate *d* (Appendix S10). Simulations used *N* patches for *N* = 5 (A, C) and *N* = 25 (B, D).

Also consistent with hypothesis, copula structure had a substantial effect on Taylor’s law for the models we considered. Taylor’s law was strongly influenced, and was often even invalidated in its assumptions of linearity and homoskedasticity, by non-normal copula structure. For normal copula structure (i.e., for simulations that gave normal-copula dependence between populations in different locations), *p*-values for the linearity and homoskedasticity tests were roughly uniformly distributed across replicate simulations and Taylor’s law appeared visually to be a reasonable approximation of the log(*v_g_*) versus log(*m_g_*) relationship (Fig. 15B,E,F). Furthermore, quadratic coefficients and curvature values were close to 0 (Fig. 15J,K), and root mean squared errors from the linear regression were relatively small (Fig. 15G). But linearity or homoskedasticity were violated more frequently for non-normal copula structure (Fig. 15A,C,D,E,F); quadratic coefficients and curvatures were frequently non-zero (Fig. 15J,K); and root mean squared errors from the linear regression were much higher (Fig. 15G). Slopes and intercepts of the linear regression were also strongly affected by copula structure (Fig. 15H,I), though some of the effect here was because linear regressions do not always adequately represent the log(*v_g_*) versus log(*m_g_*) relationship when copula structure was not normal. Thus our results substantiated the hypothesis, at least for the models we used, that Taylor’s law can be influenced by copula structure and asymmetric tail associations. This is represented in Fig. 2 by the arrow labelled “Z”.

**Figure 15:**
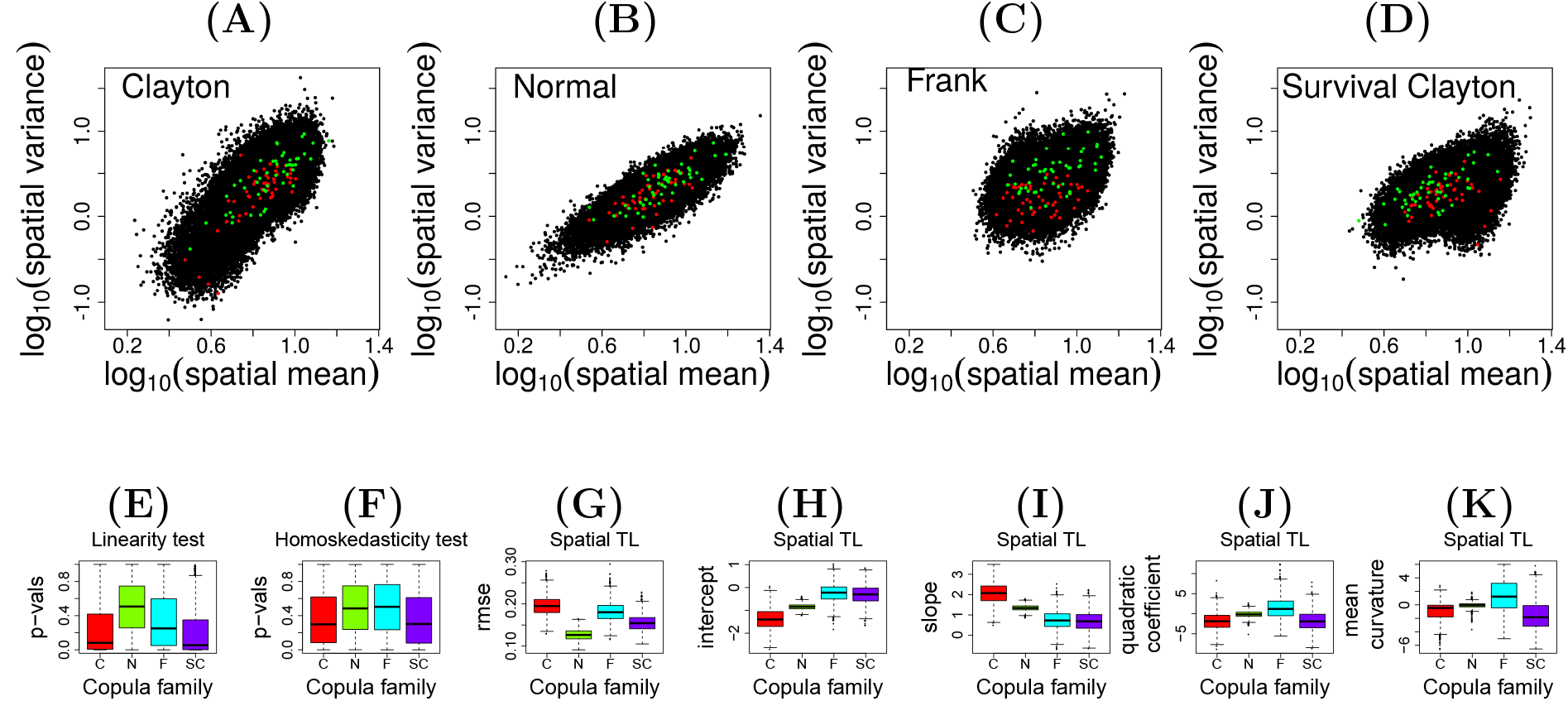
Spatial Taylor’s law results. Log(spatial variance) vs. log(spatial mean) relationships for 1000 simulations over 50 time steps for 25 locations using Clayton (A), normal (B), Frank (C), and survival Clayton (D) copula structures between locations. Results from all 1000 simulations were plotted on the same axes for each panel, but two example simulations are shown in red and green to help assess to what extent variation on the plots was between simulations or within simulations. For each simulation, we tested the linearity (E) and homoskedasticity (F) of the log(variance) vs. log(mean) plot for that simulation, and quantified the root mean squared error of points from the linear regression line (G). We also calculated the linear regression intercept (H) and slope (I), the quadratic term of a quadratic regression through the points on the log(variance) vs. log(mean) plot (J), and the curvature of that quadratic regression (K). Distributions of values across all 1000 simulations are displayed. Spearman-preserving surrogates were used, though results using Kendall-preserving surrogates were similar. See section 8 and Appendices S11 and S12 for details.

## 10 Discussion

We showed that non-normal copula structure and asymmetric tail associations are common across multiple sub-disciplines in ecology, although these facets of ecological data are only occasionally taken into account (Valpine *et al.* 2014; Anderson *et al.* 2018; Popovic *et al.* 2018). We hypothesized mechanisms that may cause non-normal copula structures and asymmetric tail associations; we discuss below how commonly some of our mechanisms may operate. We also hypothesized and demonstrated important consequences of non-normal copula structure and asymmetric tail associations for the field of ecology. For instance, the skewness of a spatial-average time series is influenced by asymmetric tail associations between its constituent time series: predominantly right-tail-associated local time series can lead to “spiky” spatially averaged time series, with large outbreaks; and predominantly left-tail-associated local time series can lead to spatially averaged time series showing accentuated “crashes”. Thus tail associations could have implications for pest and exploited species. Extinction risk and Taylor’s law can also be altered by tail association patterns across space. In our view, our results make it reasonable to suggest that a more comprehensive understanding of many ecological phenomena may be possible if a complete copula characterization of associations between variables is employed. Copula approaches are well developed, statistically (Nelsen 2006; Joe 2014; Mai & Scherer 2017), and have been introduced in accessible formats (Genest & Favre 2007; Anderson *et al.* 2018). Furthermore, open-source computer implementations exist (e.g., the `copula` and `VineCopula` packages in the R programming language). Ecologists can apply these tools immediately. We created several interrelated randomization procedures (Appendices S6, S9, S12) that have built upon existing copula methods.

The approaches we have demonstrated should apply equally well to data from tropical or temperate ecosystems. There is no reason to expect that non-normal copula structure and asymmetric tail associations should be special properties of datasets from temperate regions. The mechanisms we have proposed of non-normal copula structure and asymmetric tail associations seem equally likely to apply anywhere. For instance, the Moran effect, which underpins one of our proposed mechanisms (Fig. 2, B), is a standard mechanism that occurs whenever environmental variables influence populations. And Liebig’s law and nonlinear environmental influences on ecosystems, which underpin another one of our proposed mechanisms (Fig. 2, A), are widely demonstrated phenomena that apply in all biomes.

Our first proposed causal mechanism (Fig. 2, A) may well operate commonly, for two reasons. First, Liebig’s law and the idea of limiting nutrients are dominant paradigms in ecology, and many studies have documented nonlinear or threshold influences of environmental variables on ecological quantities. Second, fluctuations in environmental variables through time are very commonly correlated across space. Because these factors, which are the essential ingredients of the mechanism, are so common, it is reasonable to hypothesize that the mechanism may operate commonly. It may be a dominant cause of asymmetric tail associations and non-normal copula struture of ecological dependencies across space. We provide some further support for the mechanism in our discussion of green spruce aphids and winter temperature below.

There are also reasons to hypothesize that our Moran mechanism (Fig. 2, B) may operate commonly: Moran effects are common (Liebhold *et al.* 2004; Sheppard *et al.* 2016; Defriez & Reuman 2017a, b), and non-normal copula structures and asymmetric tail associations are often found in environmental variables. If intense meteorological events are also widespread, then environmental variables associated with these events should take extreme values simultaneously across large spatial areas, producing tail associations in measurements made through time at different locations. Non-extreme values may instead be associated with local phenomena, and therefore may be less correlated across large areas. Serinaldi (2008) examined the spatial dependence of rainfall in Central Italy. Gumbel or Student 2-copulas were candidates for modelling dependence, and neither of these is a normal copula. A long-term study (1950–2014) in the Loess Plateau of China (She & Xia 2018) showed that a Gumbel copula effectively modeled the spatial dependence of drought variables. The Gumbel copula demonstrates asymmetric tail associations. Bivariate copula analysis was also used in forecasting the co-occurrence of extreme events (flood or drought) over the North Sikkim Himalayas using spatial datasets (Goswami *et al.* 2018).

We suggested in the Introduction that asymmetric competitive relationships between species could yield asymmetric tail associations between abundance measurements for the species. This is another theoretical mechanism for non-normal copula structure and asymmetric tail associations, represented in Fig. 2 by the box “Asymmetric species interactions” and the arrow labelled “C”. It could be tested by analyzing copulas of abundances of competing species, sampled across space or time. We note that, whereas all the datasets we studied here have been positively associated when they were significantly associated, for negatively associated variables such as abundances of competing species, the definitions of left- and right-tail association no longer apply, strictly speaking: the left tail of one distribution corresponds to the right tail of the other. One must be careful with terminology, but it is still possible to study asymmetries of association.

Our simulations of character evolution suggest the hypothesis that changes through evolutionary time in bird and mammal BMR and body size may exhibit greater right- than left-tail association, contrary to standard normality assumptions of character evolution models. This is a hypothesis only, because the greater right-tail association shown in Fig. 8F,G could have come about in another, unknown way rather than via the mechanism we suggested which implicated asymmetric tail associations in evolutionary change. Our simulations show that asymmetric tail associations in evolutionary changes are sufficient, but may not be necessary, to produce the observed asymmetric tail associations in characters of extant species. For instance (see below, and Appendix S13 and Fig. S30), systematically missing data can also produce tail dependence and may have influenced results for the BMR-body mass datasets. Even if the hypothesized evolutionary mechanism (Fig. 2, D) is correct, our results only replace one question, i.e., why do we see greater right- than left-tail association between BMR and body mass, with another, i.e., why might we see greater right- than left-tail association in evolutionary changes in these traits?

Performing statistical tests for associations between continuous traits across different species was a primary motivating example of phylogenetic comparative methods. For example, Felsenstein’s method of phylogenetically independent contrasts (Felsestein 1985) is currently the second most-cited paper in the history of The American Naturalist journal (Huey *et al.* 2019). Frustratingly, the field still lacks a dependable procedure for dealing with branch lengths, which are a crucial input to the method. Ideally, the branch lengths used to correct for phylogenetic effects would represent the expected amounts of change for the characters that are being analyzed. Because researchers almost never have a reliable method for providing such branch lengths, most researchers rely on ultrametric trees – those for which the duration of the branch in time can be treated as a proxy for the branch length. Frequently these branch lengths are transformed to assess sensitivity to different assumptions about the degree of phylogenetic inertia displayed by the traits under study (Harmon 2018; Ives 2018). Even if one were able to simply use a time-based set of branch lengths, assigning dates to nodes in phylogenies is difficult. DNA sequence data can provide estimates of branch lengths, but these estimates are dependent on the adequacy of models which correct sequences for multiple substitutions occurring at the same location. Biases in estimating the evolutionary distance can affect downstream analyses (Phillips 2009). Additionally, changes in the rate of molecular evolution make the estimation of dates difficult (Heath & Moore 2014) even when branch lengths in time are accurately estimated.

Without reliable branch length estimates, it is difficult to interpret the significance of the magnitude of changes in traits across a tree. Developing tests of association based on copula structure may possibly lead the way to more robust methods for studying associations when we lack defensible estimates of branch lengths. We note that our phylogenetic analyses here consisted merely of simulations to assess whether interesting copula structure in a simple evolutionary process could leave a detectable signal on the trait data for extant species. Substantial work remains to be done before we have a copula-based method for analyzing data on a phylogenetic tree. But the nonparametric nature of approaches based solely on data ranks, as many of our copula methods are, may be a promising avenue for avoiding inaccuracies that arise via the highly structured model assumptions implicit in the method of phylogenetically independent contrasts and related methods.

Additionally, character evolution was simulated using one random draw from the relevant matrix per phylogeny branch. This was principally because branch lengths are often hugely uncertain. If branch lengths were well known, alternative simulation strategies may include selecting one draw from the matrix of evolutionary changes per unit time, or selecting one draw for the whole branch but rescaling the variances of the selected character changes according to the length of the branch. These choices amount to the same thing for normal copula structure, but not otherwise. Modelling choices such as these may have influenced our results. Additional research seems warranted testing the realism of our hypothesized mechanism and simulations.

Relationships between BMR and body mass relate to a trade-off between mass-specific BMR (BMR per unit body mass) and body mass itself. Copulas probably interrelate with life-history trade-off theory in additional ways beyond what we demonstrated. For instance, it is well known that energy allocation to a life function, F (e.g., reproduction) will reduce the energy that can be allocated to other functions, *G*_1_, *G*_2_, *G*_3_ (e.g., growth, predation avoidance). This is the principle of allocation. But F can trade off against any or all of the *G_i_*. Therefore, for large F, approaching absolute limitations, there may be a strong association between F and *G*_1_, for instance. For small F, there may be little association because resources not allocated to F can instead be allocated to any combination of the *G_i_*. This constitutes asymmetric tail association between F and *G*_1_. Winemiller & Rose (1992) described a three-way trade-off in fishes between age of reproductive maturity, juvenile survivorship, and fecundity. The trade-off should, in theory, produce a tight association between age of maturity and fecundity for fishes with low age of maturity, but little such association for later-maturing fishes because those fishes may invest the resources not invested in maturing quickly into either fecundity or juvenile survival. These ideas suggest that copulas may be useful for studying multi-dimensional trade-offs. But applications will require careful attention to the possible consequences of biased sampling: if the degree of completeness of a dataset is associated with one or more of the characters, then statistical artefacts can bias conclusions (Appendix S13, Fig. S30). Multi-dimensional copulas may also be the appropriate copula approach to studying multi-dimensional trade-offs. We used only bivariate analyses in this study solely for simplicity. But statistical theory on multi-dimensional copulas is also well developed (Nelsen 2006; Joe 2014; Mai & Scherer 2017), and open-source computer implementations exist for multivariate as well as bivariate copula methods (e.g., the `VineCopula` package for the R programming language). Such approaches may be a useful next step in life-history theory and other areas.

Additional mechanisms of non-normal copula structure and asymmetric tail associations probably also operate, and should be considered. For instance, measurement error may modify copula structure. Our models investigating potential causes of copula structure were intentionally simple in other respects, too, and did not include factors such as delayed density dependence, dispersal, population stage structure, trophic interactions, etc.; and we did not comprehensively explore parameter space for our models. We re-emphasize that fuller explorations, in future work, of some of our models and of variant model may be informative. We hope by enumerating a few potential mechanisms of copula structure we will inspire additional research on the potentially numerous mechanisms that may operate in diverse datasets, and their relative importance under different circumstances.

We also elaborated potential consequences of copula structure for ecological phenomena and understanding. Our results showed that the skewness of the spatial average of local time series is influenced by their tail associations. But the same logic should also apply to any collection of time series, whether associated with locations in space or not. Another potential application is time series of abundances of all species from a single community, e.g., all plants in a quadrat surveyed repeatedly over time. A large literature has focused on synchrony versus compensatory dynamics between such time series, and the influence of interspecific relationships on the variability of community properties such as total biomass (e.g., Doak *et al.* (1998), Tilman (1999), Tilman *et al.* (2006), Gonzalez & Loreau (2009)). Typically, variability of community biomass is measured with the coefficient of variation, but skewness may also be of interest because it can help characterize “spikiness” through time. Future work on copula structure of interspecific relationships in communities and its implications for community variability is likely to be valuable, in our opinion.

Although we demonstrated that tail associations between environmental variables can influence extinction risk, substantial work remains to determine the importance of this effect. First, we used a non-density-dependent model. Do similar results pertain when density dependence is involved? Second, we considered metapopulation extinction risk, but the large field of population viability analysis (PVA) via stochastic matrix modelling (Caswell 2000; Morris & Doak 2002) uses a framework in which a single population’s vital rates (e.g., life-stage-specific fecundity and survival rates) are considered to vary stochastically through time due to environmental variation. Do relationships between different vital rates exhibit asymmetric tail associations, and do tail associations influence extinction risk in this context? Finally, is the copula structure of environmental variables or vital rates changing through time, and, if so, how do such changes influence extinction risks? Climate change is known to amplify the factors that lead to extreme weather events (Hansen *et al.* 2012) and hence may alter spatial tail associations for weather variables.

Our hypotheses and results cover the presence, causes, and consequences of non-normal copula structure and asymmetric tail associations in ecological systems (Fig. 2), but this is done using a variety of datasets and models. We here take a closer look at the green spruce aphid, which simultaneously illustrates causes and consequences with one system. Green spruce aphid abundance, as measured in the data we use, is strongly positively associated with the temperature of the previous winter (Sheppard *et al.* (2016), their supplementary figure 6). Winter temperature for year *t* was here taken to be an average for December of year *t* − 1 through March of year *t*, was available for the locations of aphid sampling, and was preprocessed in the same way as Sheppard *et al.* (2016). For each of our 10 sampling locations, we therefore examined the copula of winter temperature and aphid abundance time series for the location, finding stronger left- than right-tail associations in 7 of the 8 locations for which independence of winter temperature and aphid abundance could be rejected, according to the data specific to the location (Table 8). Apparently winter temperature has an asymmetric influence on aphid abundance in that cold winters generally produce low abundances but warm winters often do not yield higher abundances than moderate winters. One of our hypothesized mechanisms (Fig. 2, A), which our modelling results supported (section 7), therefore suggests that spatial dependence between green spruce aphid counts in different locations should show stronger left- than right-tail associations. This is exactly what was observed (Table 6), providing empirical evidence supporting the mechanism (this is why the box around “Nonlinear environmental effects, Liebig’s law” and some of the arrows labelled “A” are solid instead of dashed in Fig. 2). The consequences of such tail dependence for the skewness of spatially averaged aphid counts was described previously (Fig. 13A). Thus the asymmetric influence of winter temperature ultimately causes spatially averaged aphid abundance time series to have lower skewness (i.e., less spikiness, and greater stability through time) than they would otherwise. It seems likely that asymmetric influence of winter termperature on populations may be a common phenomenon, so effects such as we have documented for green spruce aphid may be common.

**Table 8:** Summary results for analysis of copulas for winter temperature versus green spruce aphid abundance for each of the 10 sampling locations, using the same model selection and nonparametric methods detailed in section 4. The column labeled p, CvM is the p-value result for the goodness of fit test using the Cramer-von Mises statistic. The column labeled p, KS is the p-value result for the goodness of fit test using the Kolmogorov-Smirnov statistic. Entries with an NA are because model selection and other statistics were only employed for copulas for which independence was rejected (5 percent significance level).

| Site | Best-fit<br>cop-<br>ula | Best-fit<br>AIC | Normal<br>AIC | p,<br>CvM | p, KS | Model-avg.<br>LT | Model-avg.<br>UT | Model-avg.<br>LT<br>minus<br>UT | $cor_l - cor_u$ | $P_l - P_u$ | $D_u^2 - D_l^2$ |
| --- | --- | --- | --- | --- | --- | --- | --- | --- | --- | --- | --- |
| 2 | C | -10.35 | -8.19 | 0.860 | 0.830 | 0.373 | 0.019 | 0.354 | 0.152 | 0.046 | 0.024 |
| 3 | SJ | -18.79 | -14.85 | 0.520 | 0.750 | 0.529 | 0.003 | 0.526 | 0.165 | 0.079 | 0.027 |
| 4 | C | -25.48 | -17.39 | 0.520 | 0.230 | 0.703 | 0.001 | 0.702 | 0.206 | 0.080 | 0.030 |
| 5 | C | -9.37 | -7.26 | 0.640 | 0.340 | 0.393 | 0.022 | 0.371 | 0.187 | 0.073 | 0.034 |
| 6 | F | -22.13 | -18.95 | 0.810 | 0.410 | 0.262 | 0.212 | 0.050 | -0.038 | -0.024 | -0.006 |
| 7 | C | -21.56 | -19.96 | 0.320 | 0.480 | 0.564 | 0.028 | 0.536 | 0.055 | 0.023 | 0.007 |
| 8 | SJ | -7.33 | -3.72 | 0.131 | 0.079 | 0.427 | 0.012 | 0.415 | 0.147 | 0.047 | 0.016 |
| 9 | NA | NA | NA | NA | NA | NA | NA | NA | NA | NA | NA |
| 10 | SJ | -8.21 | -4.91 | 0.222 | 0.149 | 0.446 | 0.010 | 0.436 | 0.123 | 0.047 | 0.015 |
| 11 | NA | NA | NA | NA | NA | NA | NA | NA | NA | NA | NA |

We carried out most of our analyses on data ranks, *u_i_* and *v_i_*, for good reasons mentioned in sections 1, 2 and 4 and reviewed here; but under some circumstances it may also be appropriate to use techniques which parallel our nonparametric statistics but that use unranked data, *x_i_*, *y_i_*. As mentioned previously, Genest & Favre (2007) and others recommend carrying out inferences about dependence structures (which was our goal) using normalized ranks, stating explicitly that “statistical inference concerning dependence structures should always be based on ranks.” The reason for this is that measures of dependence which use unranked data conflate information about the marginal distributions of variables with information about the dependence between variables. For instance, suppose the true population densities *p*_1,*t*_ and *p*_2,*t*_ of two species of fish in a lake in year *t* are unknown, but are assessed using catch per unit effort (CPUE), a standard approach in fisheries science (Zale *et al.* 2013). The CPUE measurements, which we denote *c*_1,*t*_ and *c*_2,*t*_ may be differently correlated through time than the true densities would be, if they were known, if the function *f* relating *p_i,t_* and *c_i,t_* is nonlinear and if Pearson correlation is used. The function *f* modifies the marginal distributions, which modifies measures such as Pearson correlation that conflate dependence and marginal information. Rank-based measures such as Spearman or Kendall correlation, or any of our copula approaches, will be unaffected by nonlinear monotonic functions such as *f*. The same difficulty will pertain in any case for which a measurement of an ecological quantity is a nonlinear index of the true quantity of interest. Nevertheless, if one is certain that measurements are linearly related to the true quantities being measured, statistics based on unranked data may be more appropriate under some circumstances, in part because ecological mechanisms which produce relationships between variables are influenced not only by the ordering information in ranks but also by the relative-size information in the unranked data. Judgements of what statistics are appropriate depend, of course, on the purpose of the analysis. The goal of this study was to study dependence in isolation, so we used ranks. If one is interested in tail associations using unranked data, a partial Pearson correlation can be developed straightforwardly, by replacing *u* and *v* in equation 1 by *x* and *y*, but still computing the sum in that equation over data points with ranks *u_i_* and *v_i_* constrained by the bounds *u_i_* + *v_i_* > 2*l_b_* and *u_i_* + *v_i_* < 2*u_b_*. This would result in a different approach to tail associations that would conflate dependence and marginal information but that may also have its own utility for some applications.

Copula methods are numerous, and go well beyond the cases we have considered in this introductory paper. We hope our work helps inspire applications of copulas in ecology both of and beyond the specific tools we have employed. For instance, multivariate copulas are useful for studying relationships between multiple interacting variables, and have been used in finance to model tail risk for portfolio optimization problems (Joe & Kurowicka 2011; Czado 2019). Analogous ecological applications exist that may be amenable to the same approach, e.g., ecosystem functioning variables such as total primary productivity or carbon flux are the sum of the contributions of multiple species in the same way that the value of an investment portfolio is the sum of the values of its constituent assets. Ecologists may likewise be interested in managing the risk that an ecosystem functioning variable will take an extreme value. As another example, we assumed for this paper that univariate marginal distributions have continuous, strictly monotonic distribution functions. This was for simplicity and because the simpler approach was sufficient for our research questions. But count data in ecology have been analyzed with approaches not making this assumption (Anderson *et al.* 2018).

## Supporting information

Supporting information for main text

## 11 Acknowledgments

We thank Joel E. Cohen for making us aware of copulas and their potential ecological utility in the first place. We thank the many contributors to the large datasets we used; D. Stevens and P. Verrier for data extraction; and David Tilman, Lauren Hallet, Jonathan Walter, Thomas Anderson, Lei Zhao, Andrew Rypel, and editors and annonymous reviewers for helpful suggestions. We thank James Bell of the Rothamsted Insect Survey (RIS). The RIS, a UK Capability, is funded by the Biotechnology and Biological Sciences Research Council under the Core Capability Grant BBS/E/C/000J0200. SG, LWS and DCR were partly funded by US National Science Foundation grants 1714195 and 1442595 and the James S McDonnell Foundation.

