## Supporting information for main text for "Copulas and their potential for ecology"

#### Appendix

**S1 Transformation alters Pearson but not Spearman correlation** **1**

**S2 Proofs of statements related to Sklar's theorem in the case of continuous, strictly monotonic margins** **1**

**S3 Details of the data** **1**

**S4 Nonparametric statistics** **2**

**S5 Testing the nonparametric statistics** **2**

**S6 Surrogate testing with bivariate datasets for nonparametric statistics** **2**

**S7 Spatial resampling with multivariate datasets for nonparametric statistics** **3**

**S8 Methods for the hypothetical evolutionary mechanism for tail dependence between traits** **3**

**S9 Surrogate testing with multivariate datasets** **3**

**S10 Extinction risk noise and dispersal matrix** **4**

**S11 Effects of non-normal copula structure on Taylor's law** **4**

**S12 Another tool for producing surrogates for multivariate datasets** **5**

**S13 A means by which missing data can influence perceived copula structure** **5**

**S14 References** **5**

### S1 Transformation alters Pearson but not Spearman correlation

The data of Fig. 1A-B were generated to have standard normal marginals. Fig. S1A can be obtained from Fig. 1A via transformation of both variables by the cummulative distribution function (cdf) of the standard normal distribution, and then by the inverse of the cdf of the gamma distribution with shape and scale parameters 2. The functions `pnorm` and `qgamma` in R were used. Fig. S1B can be obtained from Fig. 1B in the same way. The Pearson correlations differ substantially after transformation, but the Spearman correlations are unchanged (Figs 1A-B, S1). A scatterplot very simlar to Fig. 1C can be obtained from Fig. 1A via transformation of both variables by the cdf of the standard normal distribution. Likewise a scatterplot similar to Fig. 1D can be obtained from Fig. 1B in the same way.

### S2 Proofs of statements related to Sklar's theorem in the case of continuous, strictly monotonic margins

Let $F$ be the bivariate distribution function of a random vector $(X, Y)$, with margins $F_X$ and $F_Y$ assumed to be continuous and strictly monotonic. We show the copula associated with $(X, Y)$, which exists and is unique by Sklar's theorem, is, in fact, the distribution function of the random variable $(U, V) = (F_X(X), F_Y(Y))$. Let $C$ denote the distribution function of $(U, V)$, which has uniform margins. By the uniqueness statement of Sklar's theorem, it suffices to show that $F(x, y) = C(F_X(x), F_Y(y))$ for all $(x, y)$ in the Euclidean plane. But $C(F_X(x), F_Y(y)) = P[U \leq F_X(x), V \leq F_Y(y)] = P[F_X(X) \leq F_X(x), F_Y(Y) \leq F_Y(y)] = P[X \leq x, Y \leq y] = F(x, y)$, where the third equality follows from the strict monotonicity assumption. In fact, that equality should hold even with non-strict monotonicity, so a stronger statement is possible (see below for more on this topic). The continuity assumption for $F_X$ and $F_Y$ is necessary, however: without it the random variables $U$ and $V$ are not uniformly distributed, so their joint distribution function is not even a copula.

Now let $D$ be any bivariate copula and let $G_A$ and $G_B$ be univariate distribution functions, so that $D(G_A(x), G_B(y))$ is guaranteed by Sklar's theorem to be a distribution function. We here show that if $G_A$ and $G_B$ are continuous and strictly monotonic, then $D(G_A(x), G_B(y))$ is the distribution function of the random variable $(G_A^{-1}(U), G_B^{-1}(V))$, where $(U, V)$ now denotes the random vector with uniform marginals associated with $D$ and $G_A^{-1}$ and $G_B^{-1}$ are the inverses of $G_A$ and $G_B$. The inverses exist because of the strict monotonicity assumption. To get to the desired result, note that the distribution function of $(G_A^{-1}(U), G_B^{-1}(V))$ is $P[G_A^{-1}(U) \leq x, G_B^{-1}(V) \leq y]$, which equals $P[U \leq G_A(x), V \leq G_B(y)]$, by strict monotonicity. But this is $D(G_A(x), G_B(y))$ because $D$ is the distribution function of $(U, V)$. In fact, a stronger statement assuming neither continuity nor strict monotonicity is possible, if one uses generalized inverses for $G_A^{-1}$ and $G_B^{-1}$ (Mai & Scherer 2017). We adopt the weaker statements for simplicity and because our applications are solely under the case of continuous, strictly monotonic margins. But see Anderson *et al.* (2018) and Popovic *et al.* (2018) for applications of copulas outside this case.

### S3 Details of the data

**Environmental data, soil C and N.** Measurments were from the central pedon of a soil core, to 1m depth.

**Phenological and population data on aphids.** Data pre-processing for the aphid data was the same as that of Sheppard *et al.* (2016), except that years for which data was unavailable were excluded rather than being replaced by median values. We also excluded one of the 11 sites they used in order to only use sites with at least 30 years data.

**Population data, plankton abundances.** These data were obtained from the Continuous Plankton Recorder (CPR) dataset, now operated and maintained by the Marine Biological Association of the United Kingdom. Data pre-processing was similar but not identical to that of Sheppard *et al.* (2017). The CPR plankton data presented in Sheppard *et al.* (2017) was aggregated in the following way. Fifty $2°$ by $2°$ location boxes were examined between $-10°$ and $10°$ longitude and between $50°$ and $60°$ latitude (Fig. S6). For each species, for each box, for each year, for each month, the average of any available CPR samples was taken to give monthly time series of population densities with some missing values. Missing monthly values were then replaced with the median value for that species for that location for that month. The data for each species for each box for each year was then averaged over all twelve months, and 26 usable annual time series were identified for wavelet analysis. The CPR plankton data used here is aggregated in the same way, except that for any species, box and year for which more than 4 months data were missing, missing values were not replaced with medians and those species, box, year combinations were not used. We used only data from the 14 locations which had at least 45 years of annual data subject to this restriction, shown by blue markers in Fig. S6. The CPR data are described in great detail elsewhere, e.g., by Batten *et al.* (2003) and Raitsos *et al.* (2014).

**Community-level data, Cedar Creek.** Above-ground biomass (Tilman 2018a) and percent cover data (Tilman 2018b) were obtained from biodiversity experiment E120 conducted at the Cedar Creek Ecosystem Science Reserve by Tilman *et al.* In 1994, 168 plots (Tilman 2018c) (each 9 m × 9 m) of 39, 35, 29, 30 and 35 replicates, were seeded with 1, 2, 4, 8, or

16 perennials, respectively, chosen randomly from a pool of 18 species (4 species, each, of C4 grasses, C3 grasses, legumes, non-legume forbs; and 2 species of woody plants). All plots received, in total, 10 $gm^{-2}$ of pure live seed in May 1994 and 5 $gm^{-2}$ in May, 1995 with seed mass divided equally among species. Treatments were maintained by weeding manually 3-4 times a year and sometimes selective herbicides were used. We downloaded the data on July, 2018 from the Cedar Creek website http://www.cedarcreek.umn.edu/research/experiments/e120 and analyzed those data for 6 years (1996-2000, 2007) for which both biomass and percent cover data were available. Plots were annually sampled by clipping different locations each year. For each available year, we analyzed above-ground biomass ($gm^{-2}$) and Shannon's biodiversity index (computed using percent cover data) averaged over the sampled strips/quadrats taken from each of 168 plots.

**Ecosystem functioning data on methane-flux.** The Great Miami Wetland Mitigation Bank was at $36°46^{'}51^{''}$ N, $84°20^{'}26^{''}$ W, in Trotwood, Montgomery County, Ohio. Additional information on the history of the site and soil types is detailed in Jarecke *et al.* (2016). Briefly, the closed static flux chamber method was used (Holland *et al.* 1999) to measure $CH_4$ fluxes at 28 locations across a hydrologic gradient (upland to seasonal wetland) at daily to weekly interval from September 2015 to September 2016. See Smyth *et al.* (2019) for additional details on greenhouse gas flux methods for this site. We considered 13 locations for which data were gathered for at least 50 dates.

### S4 Nonparametric statistics

We define $\mathrm{P}_{l_b,u_b}$, using the bounds $u+v=2l_b$ and $u+v=2u_b$ defined in the main text, as follows. For a distance $h$, we define $S(h)$ to be the number of points $(u_i, v_i)$ within the bounds and a distance less than $h$ from the line $v=u$, divided by the total number of points within the bounds. This function is defined for $h$ going from 0 to a distance $h_{\max}$ which is half the longer of the two segments obtained by intersecting the bounds with the unit square. It is easy to see $S(0)=0$ and $S(h_{\max})=1$. We define $S_i(h)$, also for $h$ going from 0 to $h_{\max}$, to be the area within the bounds and the unit square and within distance $h$ of the line $v=u$, divided by the area within the bounds and the unit square. This is the expected value of $S(h)$ for independent data. We then define

$$\mathrm{P}_{l_b,u_b} = \int_0^{h_{\max}} (S(h) - S_i(h))dh. \tag{1}$$

As for $\mathrm{cor}_{l_b,u_b}$, larger values of $\mathrm{P}_{l_b,u_b}$ indicate stronger positive association between $u$ and $v$ in the region given by the bounds. We use both statistics because $\mathrm{cor}_{l_{b_k},u_{b_k}}(u,v)$ is more familiar, but in some instances $\mathrm{P}_{l_b,u_b}$ appears to have more power to reveal tail dependence.

### S5 Testing the nonparametric statistics

Normal and Frank copula families are known to have symmetric tail dependence, i.e., strengths of dependence in the lower and upper tails are the same, for all copulas in these families (Nelsen 2006). Clayton copulas are known to have stronger lower- than upper-tail dependence, and survival Clayton copulas, being rotations of Clayton copulas by 180°, have stronger upper- than lower tail dependence (Nelsen 2006). Using the `iRho` function from the `copula` package, for each of the four families normal, Frank, Clayton, and survival Clayton, we obtained parameters for which the expected value of Spearman's $\rho$ was, separately, $0, 0.1, 0.2, ..., 0.9$. For each copula family and for each of the selected parameter values we generated $n$ points, 100 times, and thereby computed 100 values of the statistics $\mathrm{cor}_l - \mathrm{cor}_u$, $\mathrm{P}_l - \mathrm{P}_u$, and $\mathrm{D}_u^2 - \mathrm{D}_l^2$. This exercise was repeated, separately, for $n=1000$ and for $n=35$, these cases being chosen, respectively, to reflect asymptotic behavior of the statistics and their behavior on the small datasets which are common in some fields of ecology. The mean and standard error of the three difference statistics were computed for each copula family, parameter value, and value of $n$, and results were plotted against Spearman's rho (Figs S19 and S20), the expectation being that the difference statistics would be positive for Clayton copulas, negative for survival Clayton copulas (except for the case of $\rho=0$, a boundary case of these families corresponding to independence), and not significantly different from 0 for Frank and normal copulas. We used a $t$-test with a significance threshold of 0.05 to judge significance here. These expectations were fulfilled (Figs S19 and S20).

### S6 Surrogate testing with bivariate datasets for nonparametric statistics

For bivariate datasets, surrogates were produced as follows.

- Given data $(x_i, y_i)$ for $i = 1, \ldots, n$ and a one-parameter target copula family $\mathcal{B}(\theta)$ (here $\theta$ denotes the parameter), let $\tau$ be the Kendall correlation of the $x_i$ with the $y_i$ and find the parameter value $\theta_\tau$ for which $\mathcal{B}(\theta_\tau)$ has Kendall correlation $\tau$ (this is possible for the one-parameter copula families of this study using the `iTau` function of the `copula` package).
- Generate data $(a_i, b_i)$, $i = 1, \ldots, n$ from $\mathcal{B}(\theta_\tau)$.
- Permute the ordered set $(x_1, \ldots, x_n)$ such that the permutation $(x_{\sigma_x(1)}, \ldots, x_{\sigma_x(n)})$ has $\mathrm{rank}(x_{\sigma_x(i)})$ equal to $\mathrm{rank}(a_i)$ for all $i$, where $\mathrm{rank}(x_{\sigma_x(i)})$ is the rank of $x_{\sigma_x(i)}$ in the set $(x_{\sigma_x(1)}, \ldots, x_{\sigma_x(n)})$, and $\mathrm{rank}(a_i)$ is the rank of $a_i$ in the set $(a_1, \ldots, a_n)$ (here $\sigma_x$ is a permutation of the indices $1, \ldots, n$).
- Likewise permute $(y_1, \ldots, y_n)$ such that the permutation $(y_{\sigma_y(1)}, \ldots, y_{\sigma_y(n)})$ has $\mathrm{rank}(y_{\sigma_y(i)})$ equal to $\mathrm{rank}(b_i)$ for all $i$ ($\sigma_y$ is another permutation of $1, \ldots, n$).
- The surrogate dataset is $(x_{\sigma_x(i)}, y_{\sigma_y(i)})$ for $i = 1, \ldots, n$.

Our code for this algorithm is `copsurrog2d` in the `BIVAN` repository (https://github.com/sghosh89/BIVAN). Because $(x_{\sigma_x(1)}, \ldots, x_{\sigma_x(n)})$ is a permutation of $(x_1, \ldots, x_n)$ and $(y_{\sigma_y(1)}, \ldots, y_{\sigma_y(n)})$ is a permutation of $(y_1, \ldots, y_n)$, the marginal distributions of the surrogate data are exactly the same as those of the original data. Because ranks of the surrogate data are matched to ranks of the $a_i$ and $b_i$, the Kendall correlation of the surrogate dataset will be the same as that of the $a_i$ and $b_i$, and therefore will be close to $\tau$ (differences arising only through sampling variation). If instead of $\theta_\tau$ a parameter value $\theta_\rho$ is used such that the Spearman correlation of $\mathcal{B}(\theta_\rho)$ is the same as that of the $x_i$ with the $y_i$, then Spearman correlations of surrogates will very nearly equal (except for sampling variation) the Spearman correlation of the empirical data.

### S7 Spatial resampling with multivariate datasets for nonparametric statistics

For each pair of locations $i \neq j$, the statistic of interest, here denoted $s_{ij}$ ($\mathrm{cor}_{l_b,u_b}$, $\mathrm{P}_{l_b,u_b}$, $\mathrm{D}^2_{l_b,u_b}$, $\mathrm{cor}_l - \mathrm{cor}_u$, $\mathrm{P}_l - \mathrm{P}_u$, or $\mathrm{D}^2_u - \mathrm{D}^2_l$) was computed for those locations. The values $s_{ij}$ were then arranged in a matrix, $S$, with diagonal entries NA. The mean of the non-NA entries of $S$ was computed. For $\mathrm{cor}_{l_b,u_b}$, $\mathrm{P}_{l_b,u_b}$, or $\mathrm{D}^2_{l_b,u_b}$ this mean characterizes the average strength of dependence, across all pairs of distinct locations, for data in the portion of the distributions given by the bounds $l_b$ and $u_b$. For $\mathrm{cor}_l - \mathrm{cor}_u$, $\mathrm{P}_l - \mathrm{P}_u$, or $\mathrm{D}^2_u - \mathrm{D}^2_l$, the mean characterizes average asymmetry of tail dependence across all pairs of locations. Confidence intervals for the value of $\mathrm{mean}(S)$ were then computed as quantiles of a distribution of bootstrapped values. If $\mathrm{mean}(S)$ was significantly different from 0, as revealed by the confidence intervals, for the statistics $\mathrm{cor}_l - \mathrm{cor}_u$, $\mathrm{P}_l - \mathrm{P}_u$, or $\mathrm{D}^2_u - \mathrm{D}^2_l$, then tail dependence was significantly asymmetric, in contrast to a normal-copula null hypothesis. Bootstrapping was performed by resampling the locations of measurement, with replacement. This corresponds to resampling, with replacement, the rows and columns of $S$, and is described in detail elsewhere (Bjørnstad & Falck 2001 ; Walter *et al.* 2017).

### S8 Methods for the hypothetical evolutionary mechanism for tail dependence between traits

The matrices from which root character states and changes across phylogeny branches were randomly chosen were generated as follows. For extreme right-tail dependent noise, we first produced a 1,000,000 by 2 matrix, $m$, filled with 2,000,000 independent standard-normal random draws. Then, each row of $m$ was identified for a first set of modifications randomly and independently with a 50% chance; for those rows identified, every element of the row was replaced with the absolute value of the first element of the row (including the first element itself). For other rows, every element in the row was replaced by the negative of its own absolute value. This produces right-tail dependent noise, normally distributed for each of the two locations. If left-tail dependent noise was desired, we took the negative of the entire matrix. The moderately left-tail dependent matrix was based on a Clayton copula with parameter chosen (using the `iRho` function in the `copula` package) so that the Spearman correlation would be the same as for the extreme tail dependence datasets. The moderately right-tail dependent matrix and the symmetric tail dependence matrix were constructed similarly, but using an survival Clayton copula and an normal copula, respectively.

### S9 Surrogate testing with multivariate datasets

For multivariate datasets, surrogates were produced as follows:

- Given data $x_i(t)$ for $i = 1, \ldots, N$ and $t = 1, \ldots, T$, let $\tau_{ij}$ be the Spearman (or Kendall) correlation of $x_i(t)$ and $x_j(t)$.
- Calculate the covariance matrix of a multivariate normal distribution with standard-normal marginals and with pairwise Spearman (or Kendall) correlations equal to the $\tau_{ij}$. This is possible. using the `iRho` function (or the `iTau` function) of the `copula` package

- Generate data $a_i(t)$ for $i = 1, \ldots, N$ and $t = 1, \ldots, T$ by taking $T$ independent draws from the multivariate normal distribution of the previous step.
- For any $i, t$ for which the datum $x_i(t)$ was missing, replace $a_i(t)$ by `NA`.
- For each $k$ and $i$ for which $a_i(t)$ is not `NA`, replace the $k$th-smallest (non-`NA`) element of the time series $a_i(t)$, $t = 1, \ldots, T$ by the $k$th smallest (non-`NA`) element of $x_i(t)$, $t = 1, \ldots, T$. Call the result $b_i(t)$. This is a surrogate dataset.

Our code for this algorithm is `ncsurrog` in the `BIVAN` repository (https://github.com/sghosh89/BIVAN). The code for the Kendall version of the algorithm throws an error if there are ties in any of the time series. This is because the Kendall correlation in the very commonly used `cor` function in R is the Kendall tau-b coefficient in the event of ties, and index which differs in its definition from the commonly used tau-a that works when there are no ties; and `tau` and `iTau` are intended for continuous copulas, which cannot produce data with ties. Our code also throws an error in the event that the matrix generated in the second step is not positive semidefinite. Because ties are common in our data, we only used the Spearman case in analyses using `ncsurrog`, though, for expansibility, the `ncsurrog` code itself is written to handle the Kendall case as well when there are no ties.

Because, for each $i$, the final time series $b_i(t)$ is a permuted version of $x_i(t)$ (and with `NA`s for the same $t$), the marginal distributions of the surrogate dataset (i.e., the distributions of values for each sampling location, $i$) are exactly the same as for the original data. Because ranks are preserved in passing from $a_i(t)$ to $b_i(t)$, pairwise Spearman (or Kendall) correlations for the $b_i(t)$ will be the same as those of the $a_i(t)$. Discrepancies between the $\tau_{ij}$ and the pairwise Spearman (or Kendall) correlations of the $b_i(t)$ will therefore come about only from the presence of NAs and from sampling variation. These discrepancies should be minor if the numbers of NAs are not too large and $T$ is not too small.

### S10 Extinction risk noise and dispersal matrix

Noise was generated by first producing a 25 by $N$ matrix, $m$, of independent standard-normal random draws. Then, each row of $m$ was identified for a first set of modifications randomly and independently with a 50% chance; for those rows identified, every element of the row was replaced with the absolute value of the first element of the row (including the first element itself). For other rows, every element in the row was replaced by the negative of its own absolute value. This produces right-tail dependent noise, normally distributed for each location. If left-tail dependent noise is desired, take the negative of the entire matrix.

For local dispersal, the $N \times N$ dispersal matrix $D$ was generated by assuming, for simplicity, that the $N$ habitat patches were arranged, evenly spaced, in a line. A fraction $d$ of each population dispersed during any given time step, equally distributed to the two or one nearest neighbors. For global dispersal, the fraction $d$ that dispersed from each patch was divided up equally among all other patches.

### S11 Effects of non-normal copula structure on Taylor's law

As described in the main text, we examined using simulations whether copula structure influences spatial Taylor's law; details are provided here. First, an $n \times n$ covariance matrix $\Sigma$ was created with unit diagonal entries and off-diagonal entries $\rho = 0.7$; $n = 25$ was the number of locations at which populations were sampled. Then $N \times 1000$ random vectors were generated from a multivariate normal distribution with mean vector $(0, \ldots, 0)$ and covariance matrix $\Sigma$. These were organized into an $N \times n \times 1000$ array representing 1000 replicate simulations of (temporally independent, in this example) spatio-temporal population dynamics measured in $n$ locations over $N$ time steps. We used $N = 50$. The cdf of the standard normal distribution was applied to these numbers, followed by the inverse of the cdf of a gamma distribution with shape and scale parameters 7.5 and 1. These transformations made gamma distributed the marginal distributions of populations in each sampling location for each simulation. A skewed population marginal distribution is an important factor in Taylor's law (Cohen & Xu 2015; Reuman *et al.* 2017). We refer henceforth to the resulting $N \times n \times 1000$ array as our *normal-copula population array* for this simulation exercise; copula structure of dependences between populations in different locations was normal for this array. The array contains 1000 population matrices.

We then generated a series of arrays that were the same dimensions ($N \times n \times 1000$) as the normal-copula population array, and that were statistically similar to the normal-copula population array in all ways except the new arrays were constructed to have Clayton (respectively, Frank, survival Clayton) copula structure for the dependences between populations in different sampling locations within a simulation. We refer to the new arrays as our *Clayton-copula*, *Frank-copula*, and *Survival Clayton-copula population arrays*, respectively. The Clayton-copula array was constructed as follows; the others were constructed analogously. First, we constructed an $n$-dimensional Clayton copula with parameter chosen so that the Spearman correlations between each pair of the $n$ components were the same as those of the normal distribution constructed above. This was done using the `rho` and `iRho` functions in the `copula` package in R. Because the Clayton copula is a one-paremeter family, this step relies on the fact that all pairs of locations were equally correlated with each other in the original multivariate normal distribution constructed above. We then randomized each simulation in the normal-copula population array in a special way (see Appendix S12) so that the resulting surrogate/randomized dataset had the copula structure of the Clayton copula just constructed. The randomization procedure worked by permuting time series of values in the normal-copula array, separately for each sampling location and simulation. So population marginal distrbutions for each location and simulation were *exactly* the same as those in the normal-copula population array. Permutations were done in such a way that resulting pairwise Spearman correlations were the same as the Clayton copula constructed above, and hence were the same as the normal-copula array, except for sampling variation. Thus, to summarize using mathematical notation, if $m_{t,j}^{(\mathrm{N},k)}$ is our normal-copula population array (the three dimensions of the array are indexed, respectively, by $t$, $j$ and $k$, and $m_{t,j}^{(\mathrm{C},k)}$ is our Clayton-copula population array, then, for any given $j$ and $k$, the sets $\{m_{t,j}^{(\mathrm{N},k)} : t = 1, \ldots, N\}$ and $\{m_{t,j}^{(\mathrm{C},k)} : t = 1, \ldots, N\}$ are exactly the same (when considered as unordered sets); and for any given $j_1$, $j_2$ and $k$ with $j_1 \neq j_2$, the Spearman correlations $\mathrm{cor}_t(m_{t,j_1}^{(\mathrm{C},k)}, m_{t,j_2}^{(\mathrm{C},k)})$ and $\mathrm{cor}_t(m_{t,j_1}^{(\mathrm{N},k)}, m_{t,j_2}^{(\mathrm{N},k)})$ are the same, to within sampling variation. The whole exercise was repeated preserving Kendall instead of Spearman correlations.

### S12 Another tool for producing surrogates for multivariate datasets

Suppose one has data $x_i(t)$ for $i = 1, \ldots, N$ and $t = 1, \ldots, T$, with no missing values and no ties, and an $N$-dimensional copula $C$. The following algorithm produces a surrogate dataset $y_i(t)$ ($i = 1, \ldots, N$, $t = 1, \ldots, T$) such that: 1) the Kendall and Spearman correlations $\mathrm{cor}_t(y_i(t), y_j(t))$ are the same, up to sampling variation, as the Kendall and Spearman correlations, respectively, of the corresponding components of $C$, for all $i \neq j$; and 2) for any $i$, the values of the unordered set $\{y_i(t) : t = 1, \ldots, T\}$ are the same as the values of the unordered set $\{x_i(t) : t = 1, \ldots, T\}$.

- Generate data $a_i(t)$ for $i = 1, \ldots, N$ and $t = 1, \ldots, T$ by taking $T$ independent draws from $C$.
- For each $k$ and $i$, replace the $k$th-smallest element of the time series $a_i(t)$, $t = 1, \ldots, T$ by the $k$th smallest element of $x_i(t)$, $t = 1, \ldots, T$. Call the result $y_i(t)$. This is the surrogate dataset.

### S13 A means by which missing data can influence perceived copula structure

To illustrate the potential for missing data to influence perceived copula structure, we first generated 5000 points from a bivariate normal copula with parameter 0.8 (Fig. S30A), and then randomly deleted 80% of the points that occurred in the upper-right triangle of the copula plot. Data were then re-ranked as described in the Introduction to produce a new copula plot based only on the censored dataset (Fig. S30B). Copula structure in the tails was affected. Although real datasets have typically not been censored in the precise way we censored our artificial data, it may be common for species characters to be measured on a subset of the extant species from the clade of interest, and the probability of a species having been included in the dataset may depend on one or both of the character values. For instance, if larger or smaller bird or mammal species were more likely to have been measured for body mass or BMR, such accidental censoring of the data could have influenced our measurements of tail dependence. We have no reason to believe this is the case for our BMR versus body mass datasets, but we admit the possibility. Any systematic use of copulas to study life-history tradeoffs should take the possibility into account of artefacts such as described here.

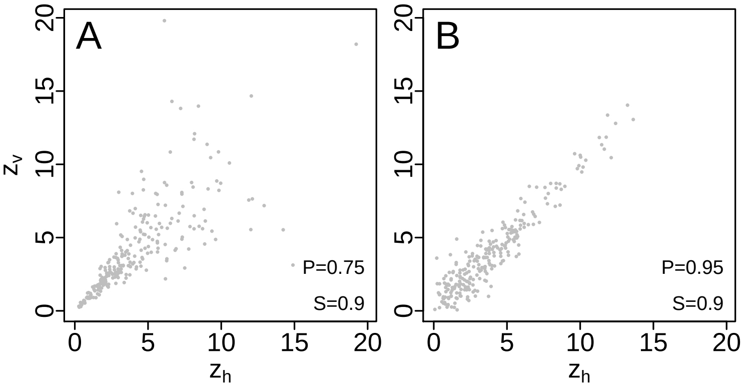

Figure S1: Transformation alters Pearson but not Spearman correlation. Displayed data are transformed from the data pictured on Fig. 1A-B. See Appendix S1 for details. P=Pearson, S=Spearman correlation.

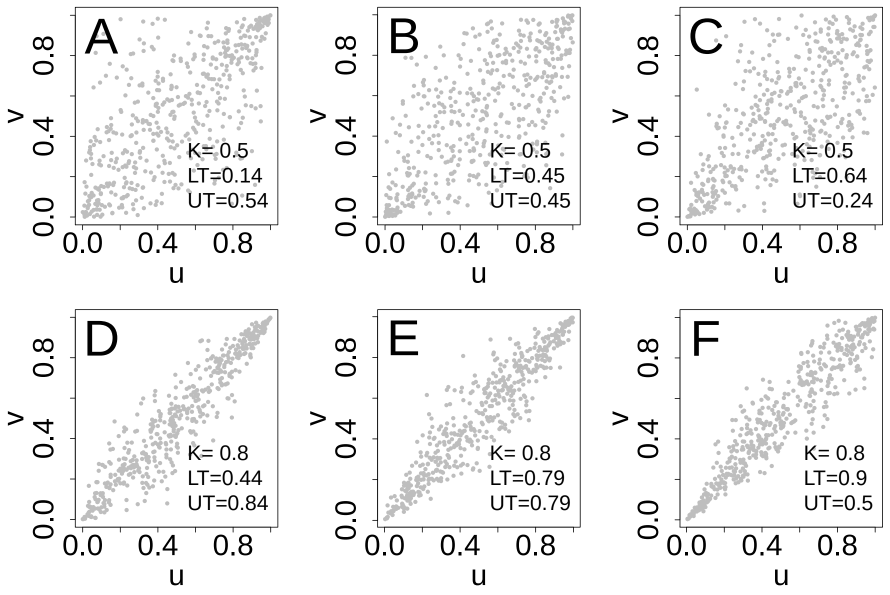

Figure S2: Lower- and upper-tail dependence can vary independently of correlation. (A-C) Data generated using the BB1 copula (see section 2 in the main text) with three different sets of parameters chosen to produce a variety of lower-tail (LT) and upper-tail (UT) dependence strengths, but with the same Kendall correlation (K) (to two digits) in all cases. (D-F) Similar to A-C, but with stronger correlation. Asymmetry of tail dependence (LT-UT) is the same within each column. See section **??** in the main text for a formal description of tail dependence and other measures of association in the tails.

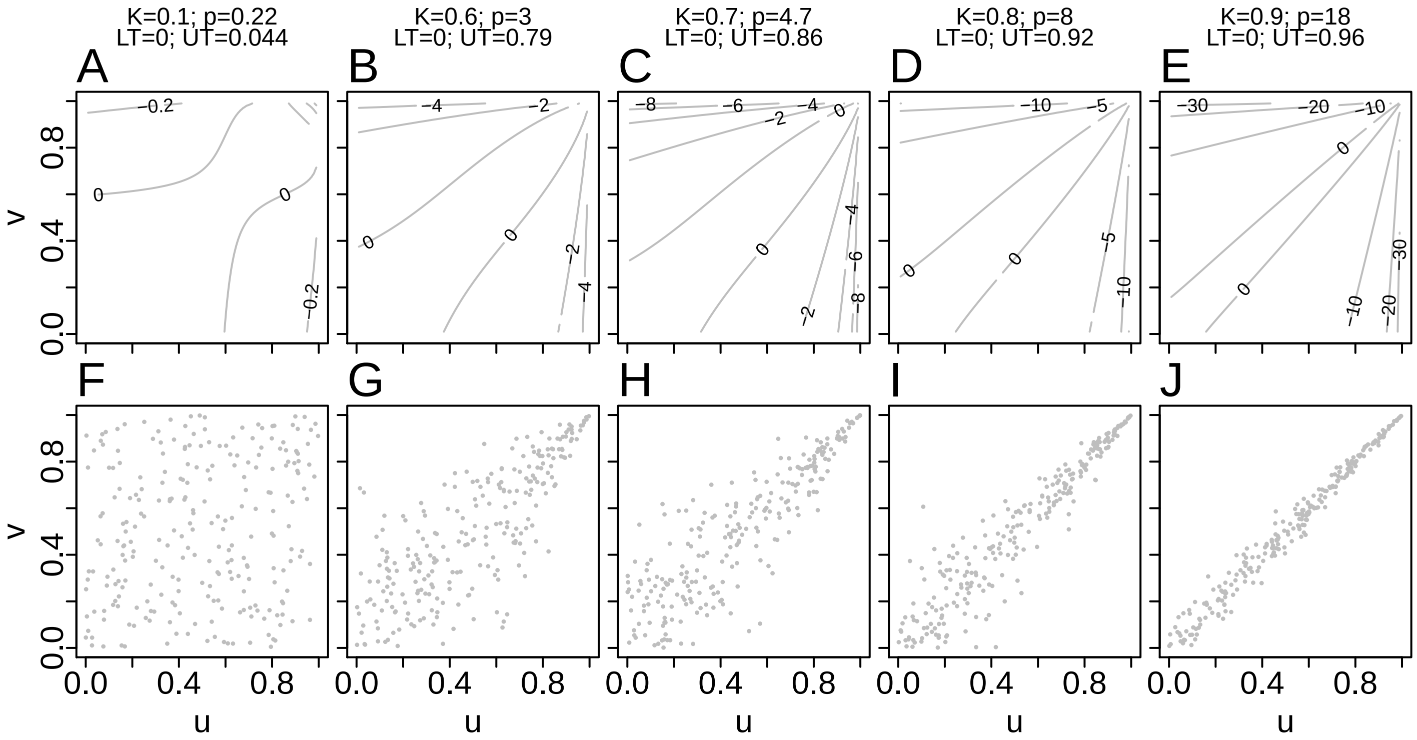

Figure S3: Log-transformed pdfs (A-E) and samples (F-J) from example survival Clayton copulas. K is Kendall correlation; p is the value of the parameter for the survival Clayton family (it is a one-parameter family); and LT and UT are the measures of lower- and upper-tail dependence, respectively. The parameter range for the family is $p \in (0, \infty)$, upper-tail dependence is $2^{-1/p}$ and lower-tail dependence is 0.

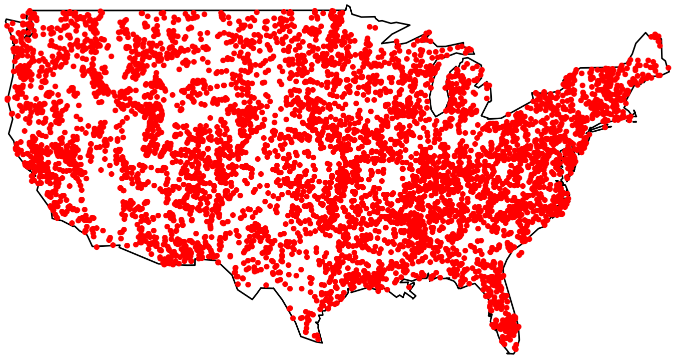

Figure S4: Locations of central pedons sampled for soil organic carbon and total soil nitrogen (Mg C or N per hectare of soil surface) at 5907 sites across the coterminous United States.

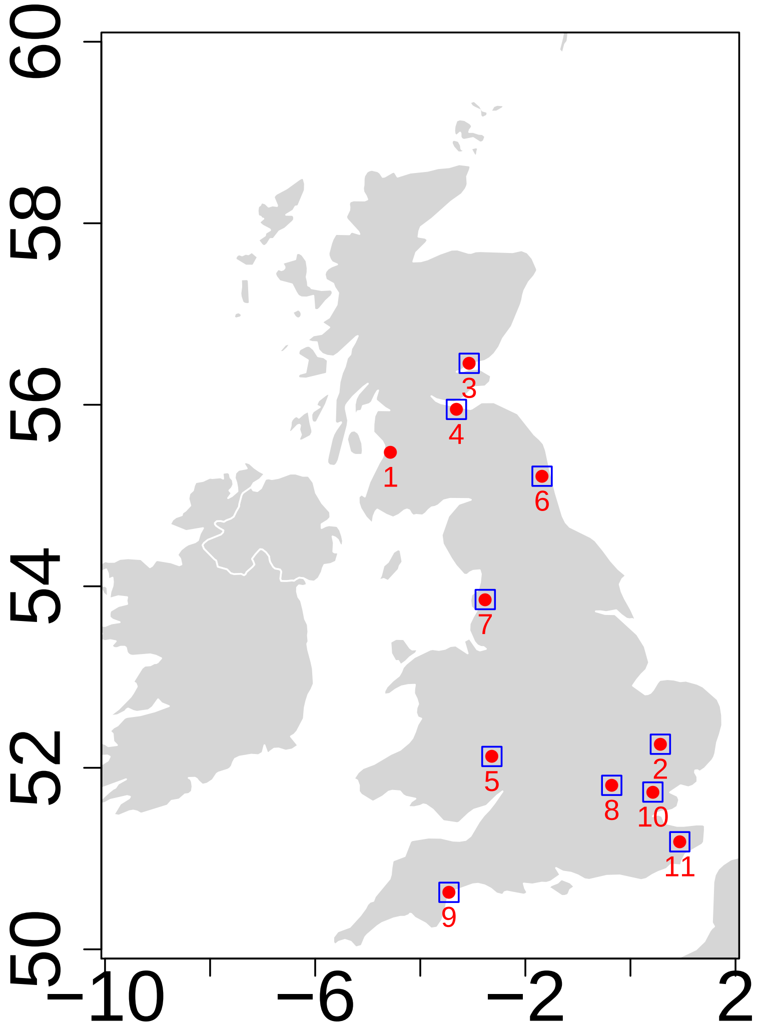

Figure S5: Locations of the Rothamsted Insect Survey suction traps from which data were available are indicated by the red circles, out of which only blue boxes had at least 30 years of data and were used for our study.

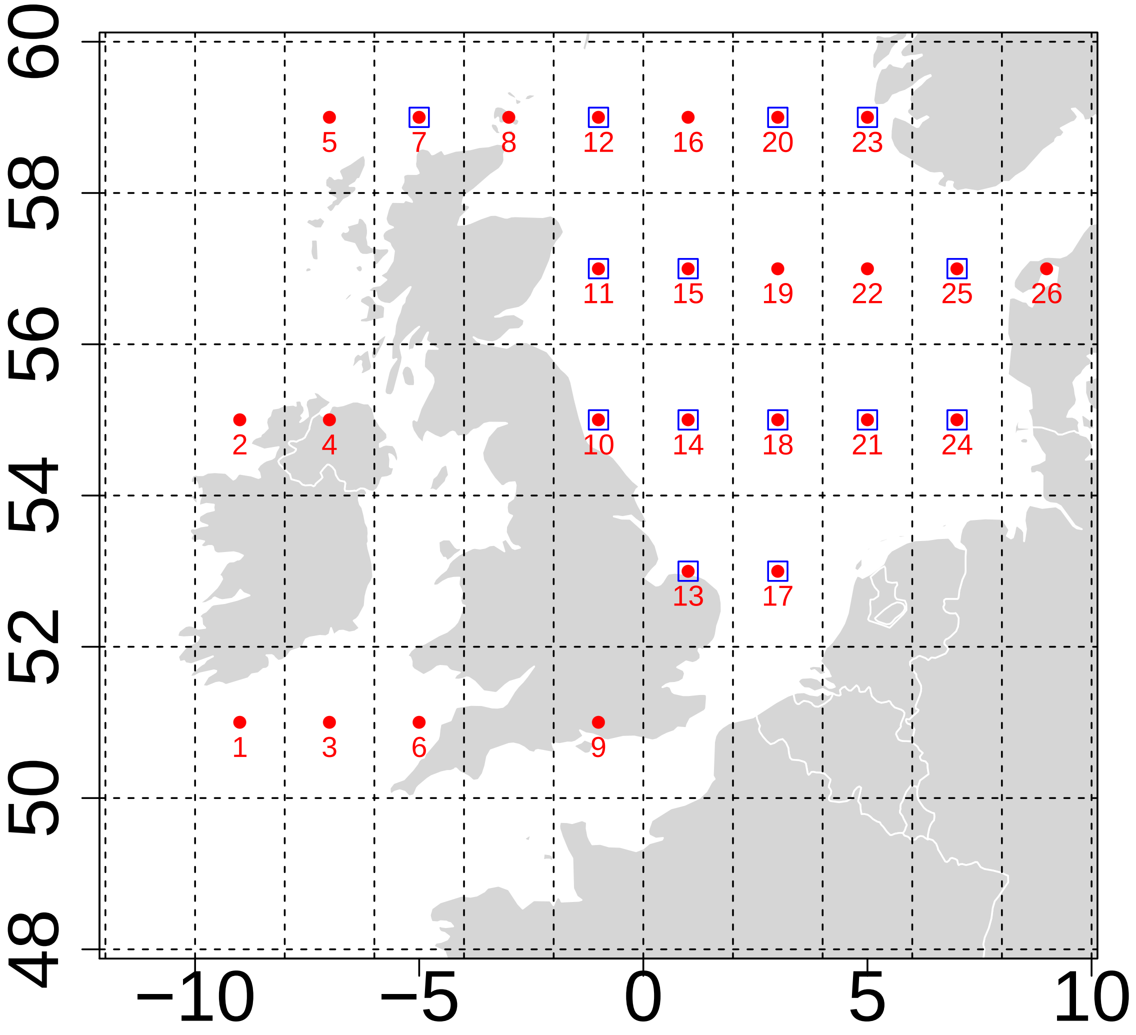

Figure S6: CPR data were available from 26 locations (2° × 2° regions on the map) around UK seas (red circles), out of which only blue boxes had at least 45 years of data and were used for our study.

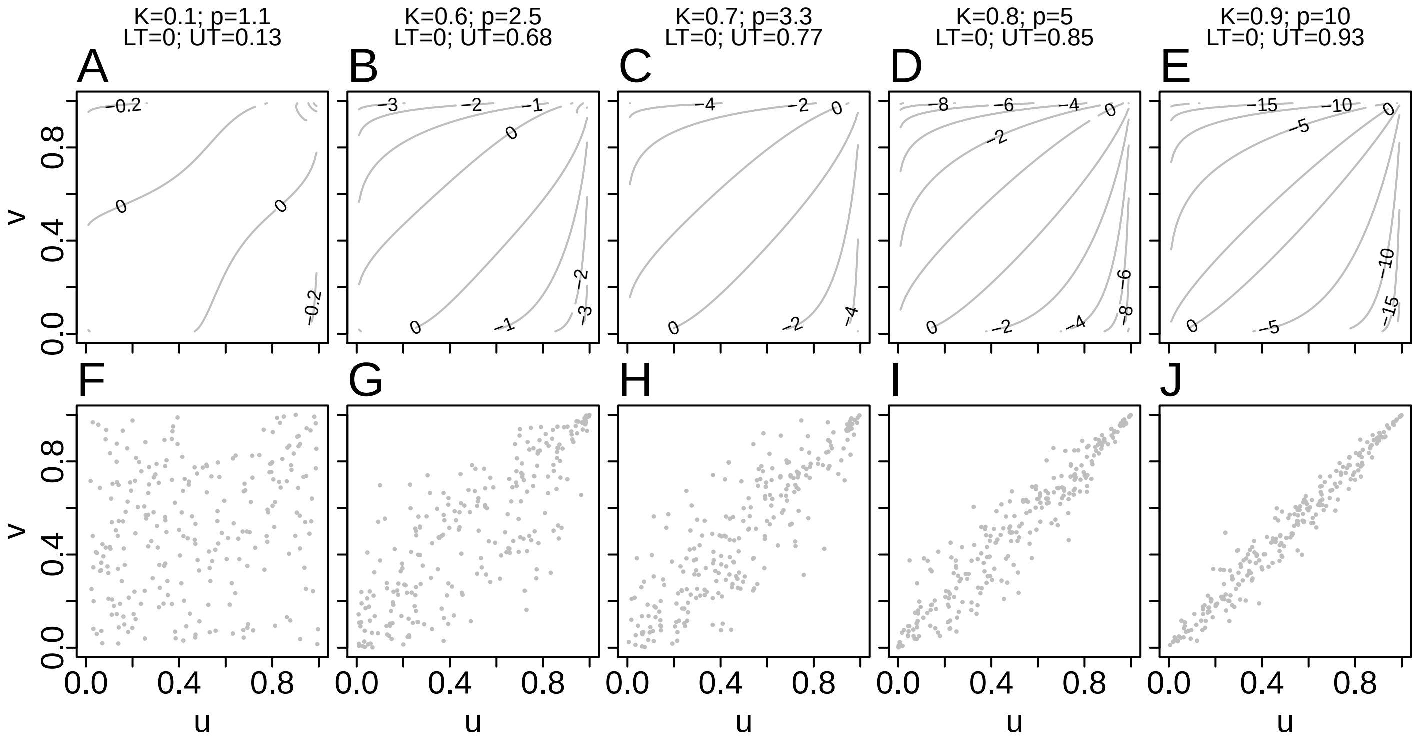

Figure S7: Log-transformed pdfs (A-E) and samples (F-J) from example Gumbel copulas. K is Kendall correlation; $p$ is the value of the parameter for the Gumbel family (it is a one-parameter family); and LT and UT are the measures of lower- and upper-tail dependence, respectively. The parameter range for the family is $p \in [1, \infty)$, lower-tail dependence is 0 and upper-tail dependence is $2 - 2^{1/p}$.

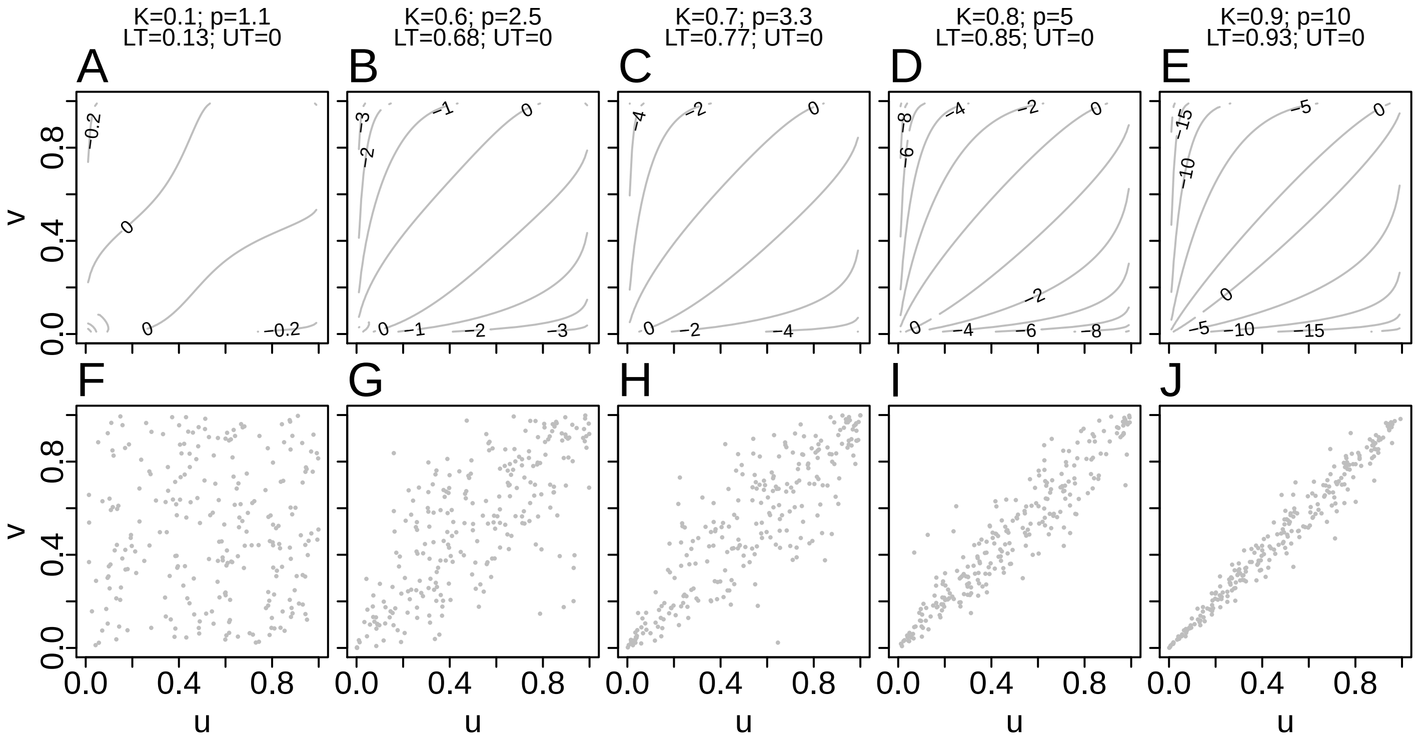

Figure S8: Log-transformed pdfs (A-E) and samples (F-J) from example survival Gumbel copulas. K is Kendall correlation; $p$ is the value of the parameter for the survival Gumbel family (it is a one-parameter family); and LT and UT are the measures of lower- and upper-tail dependence, respectively. The parameter range for the family is $p \in [1, \infty)$, upper-tail dependence is 0 and lower-tail dependence is $2 - 2^{1/p}$.

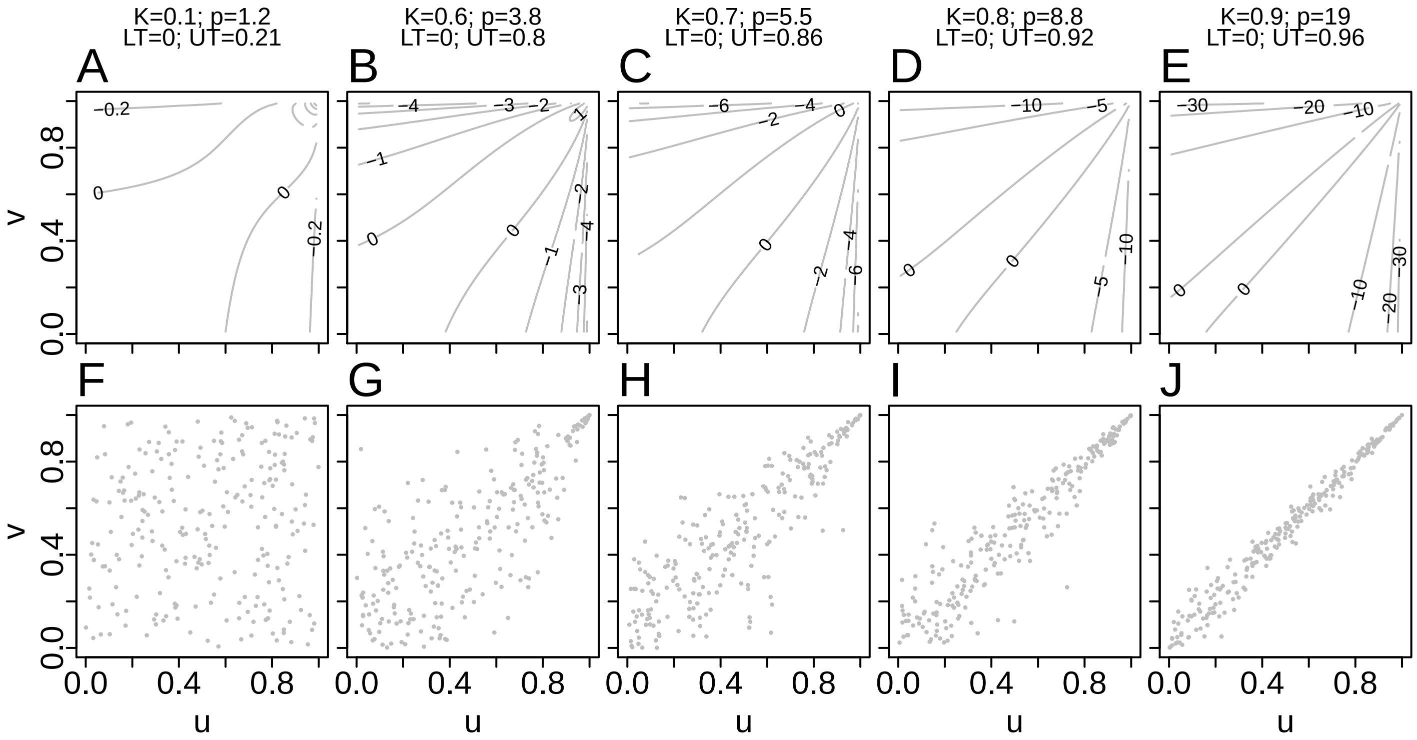

Figure S9: Log-transformed pdfs (A-E) and samples (F-J) from example Joe copulas. K is Kendall correlation; $p$ is the value of the parameter for the Joe family (it is a one-parameter family); and LT and UT are the measures of lower- and upper-tail dependence, respectively. The parameter range for the family is $p \in [1, \infty)$, lower-tail dependence is 0 and upper-tail dependence is $2 - 2^{1/p}$.

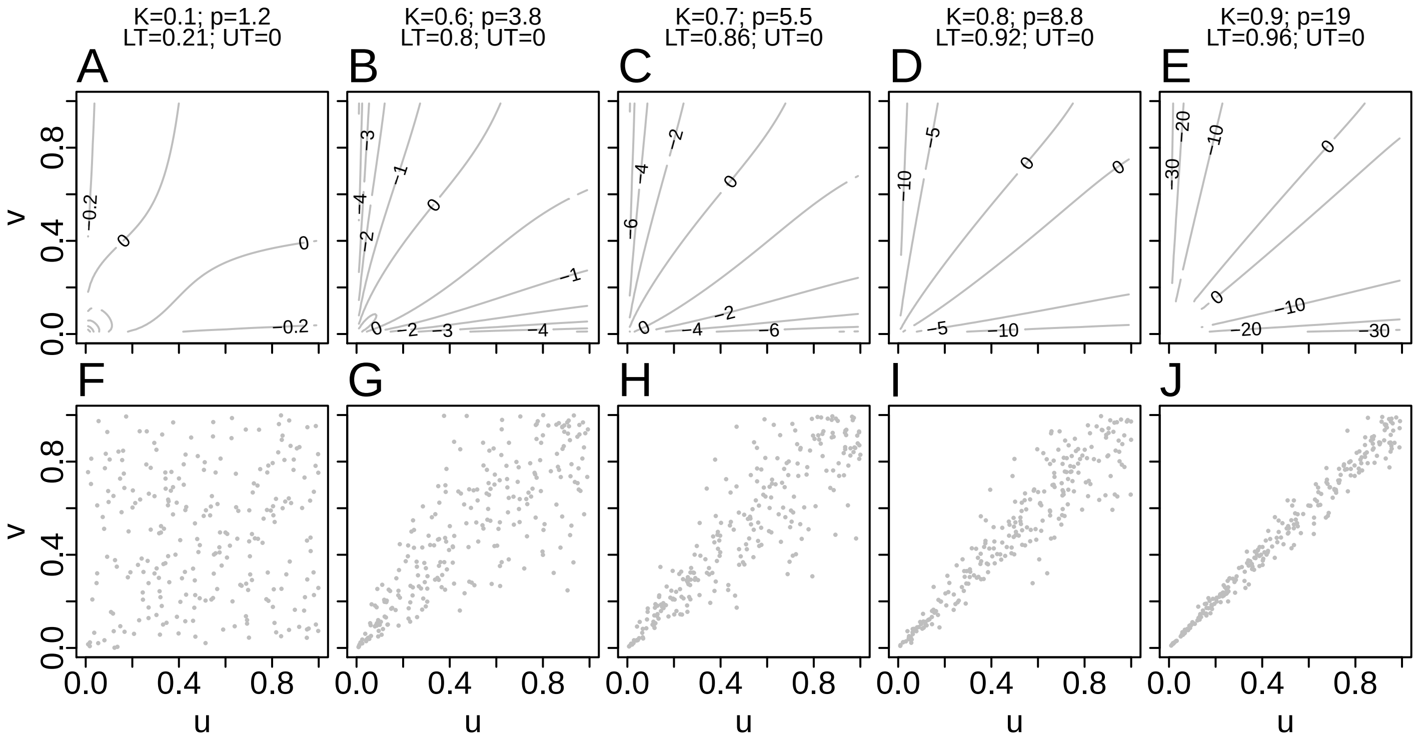

Figure S10: Log-transformed pdfs (A-E) and samples (F-J) from example survival Joe copulas. K is Kendall correlation; $p$ is the value of the parameter for the survival Joe family (it is a one-parameter family); and LT and UT are the measures of lower- and upper-tail dependence, respectively. The parameter range for the family is $p \in [1, \infty)$, upper-tail dependence is 0 and lower-tail dependence is $2 - 2^{1/p}$.

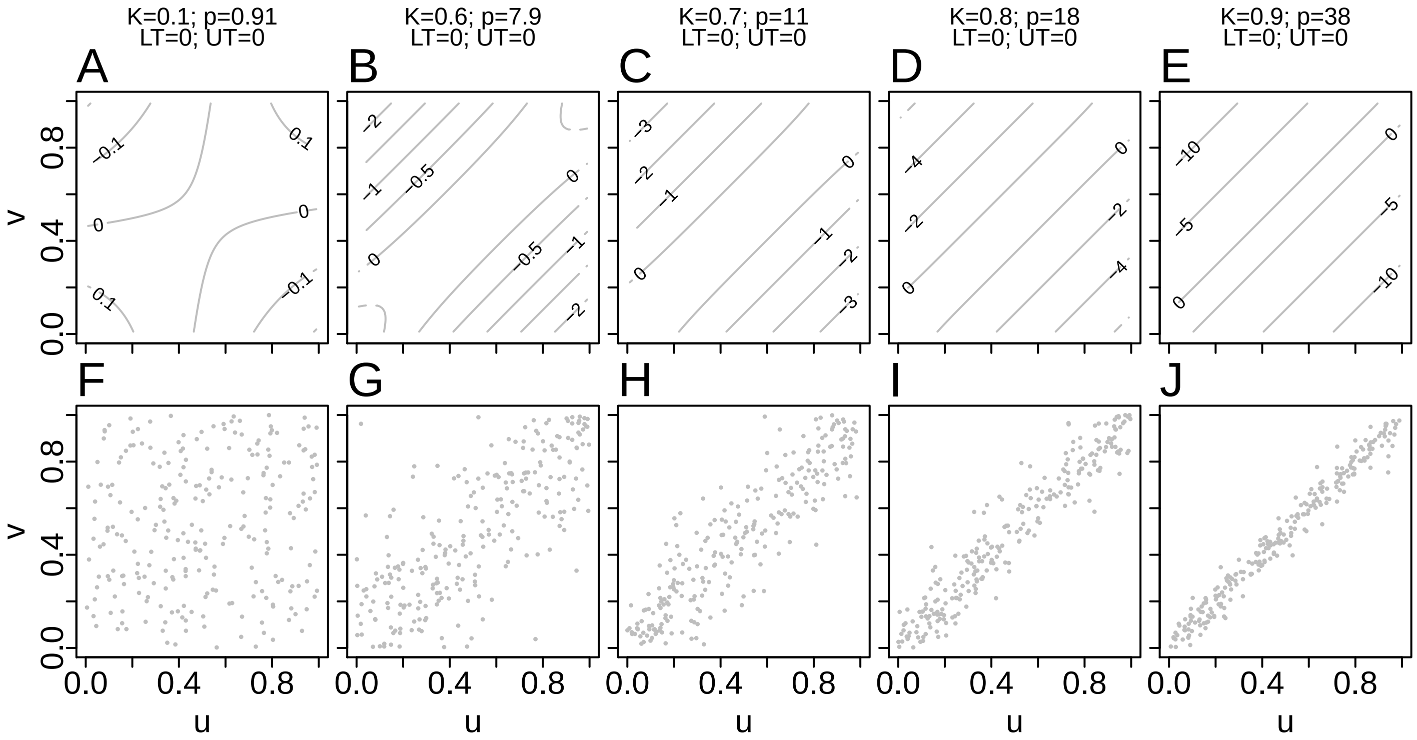

Figure S11: Log-transformed pdfs (A-E) and samples (F-J) from example Frank copulas. K is Kendall correlation; $p$ is the value of the parameter for the Frank family (it is a one-parameter family); and LT and UT are the measures of lower- and upper-tail dependence, respectively. The parameter range for the family is $p \in (0, \infty)$, lower-tail dependence is 0 and upper-tail dependence is 0.

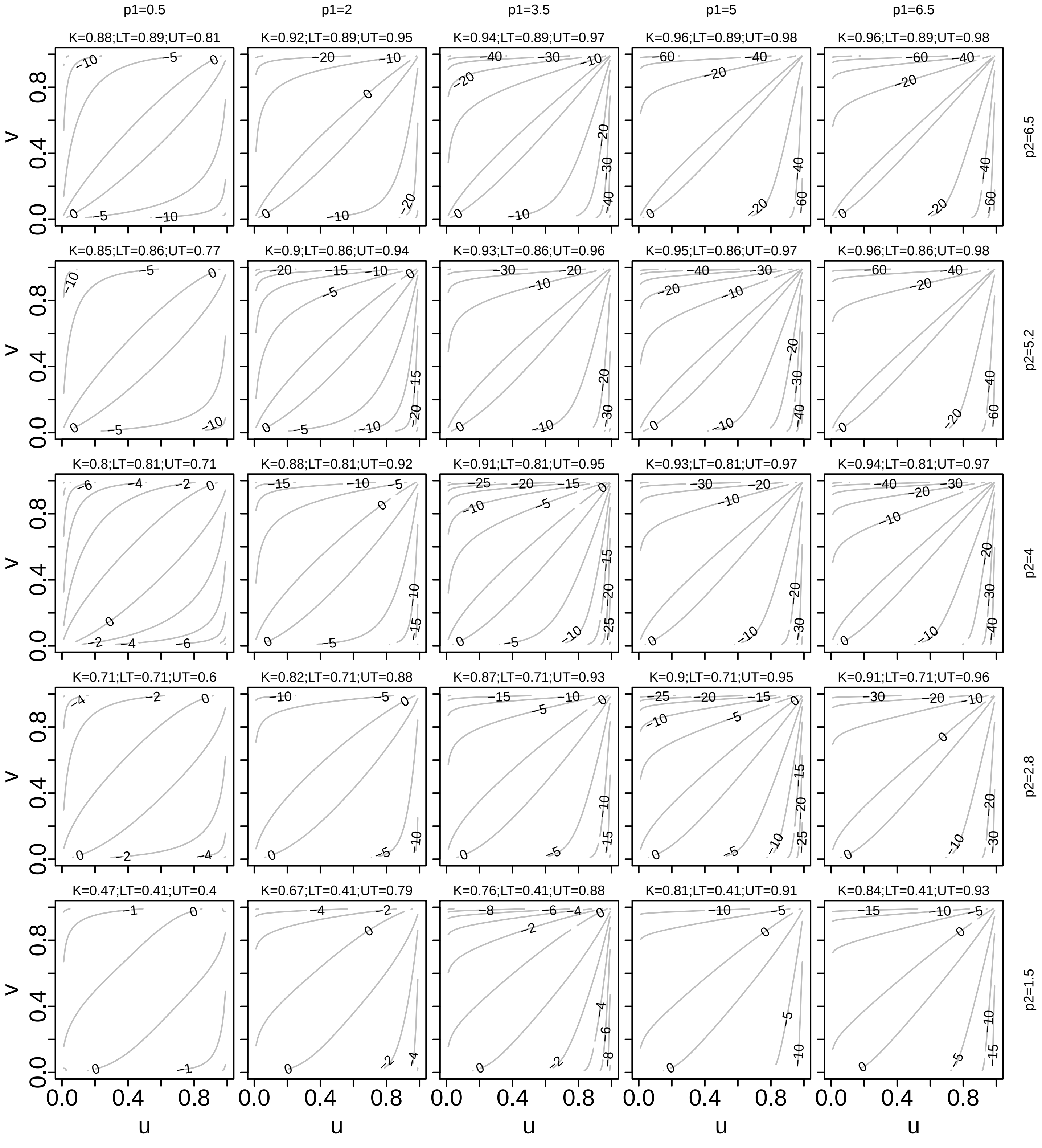

Figure S12: Log-transformed pdfs for example survival BB1 copulas. K is Kendall correlation; $p_1$ and $p_2$ denote the two parameters of the family (it is a two-parameter family); and LT and UT are the measures of lower- and upper-tail dependence, respectively. The parameter ranges for the family are $p_1 \in (0, \infty)$ and $p_2 \in [1, \infty)$, lower-tail dependence is $2 - 2^{1/p_2}$ and upper-tail dependence is $2^{-1/(p_1 p_2)}$.

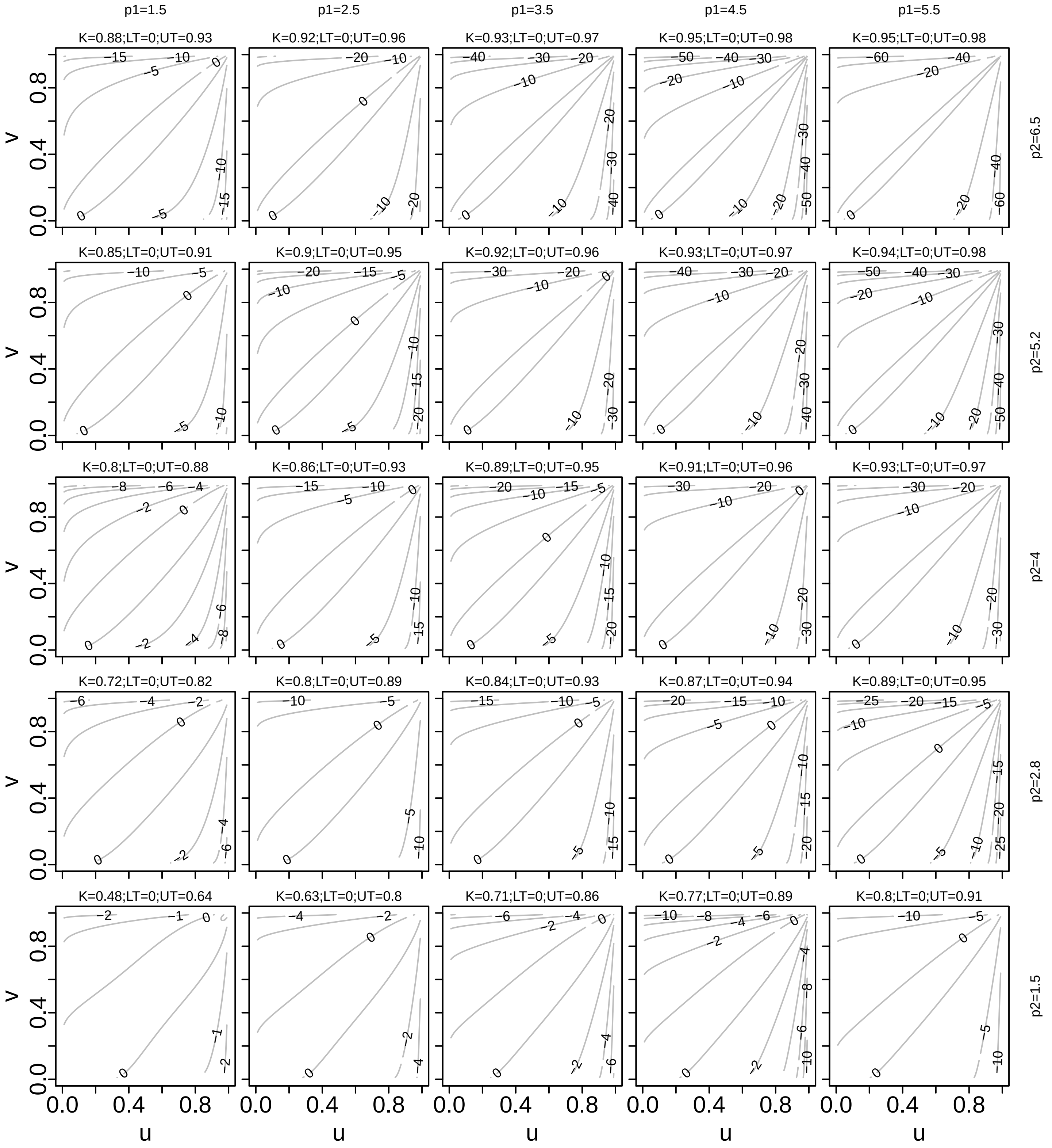

Figure S13: Log-transformed pdfs for example BB6 copulas. K is Kendall correlation; $p_1$ and $p_2$ denote the two parameters of the family (it is a two-parameter family); and LT and UT are the measures of lower- and upper-tail dependence, respectively. The parameter ranges for the family are $p_1 \in [1, \infty)$ and $p_2 \in [1, \infty)$, lower-tail dependence is 0 and upper-tail dependence is $2 - 2^{1/(p_1 p_2)}$.

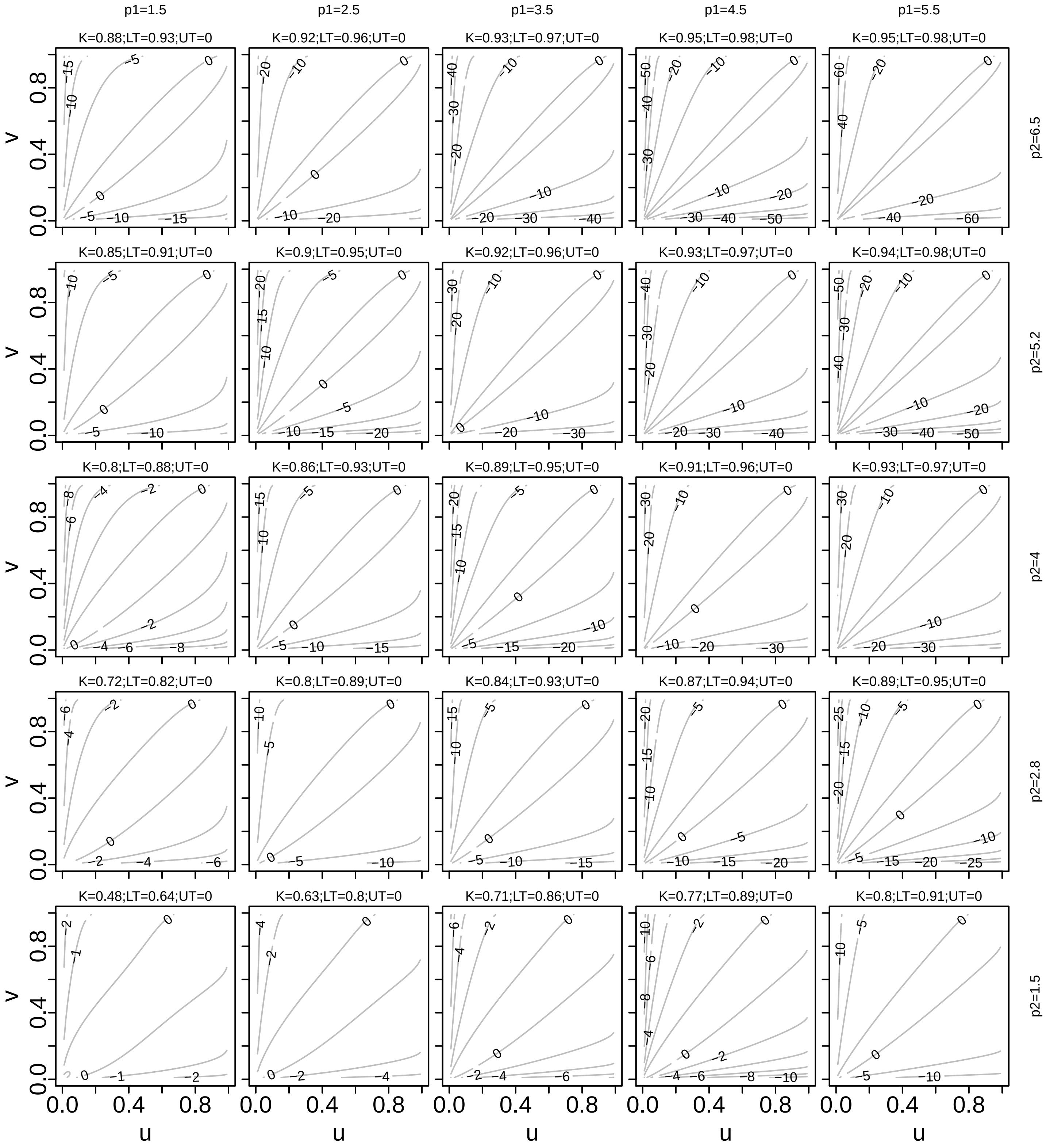

Figure S14: Log-transformed pdfs for example survival BB6 copulas. K is Kendall correlation; $p_1$ and $p_2$ denote the two parameters of the family (it is a two-parameter family); and LT and UT are the measures of lower- and upper-tail dependence, respectively. The parameter ranges for the family are $p_1 \in [1, \infty)$ and $p_2 \in [1, \infty)$, upper-tail dependence is 0 and lower-tail dependence is $2 - 2^{1/(p_1 p_2)}$.

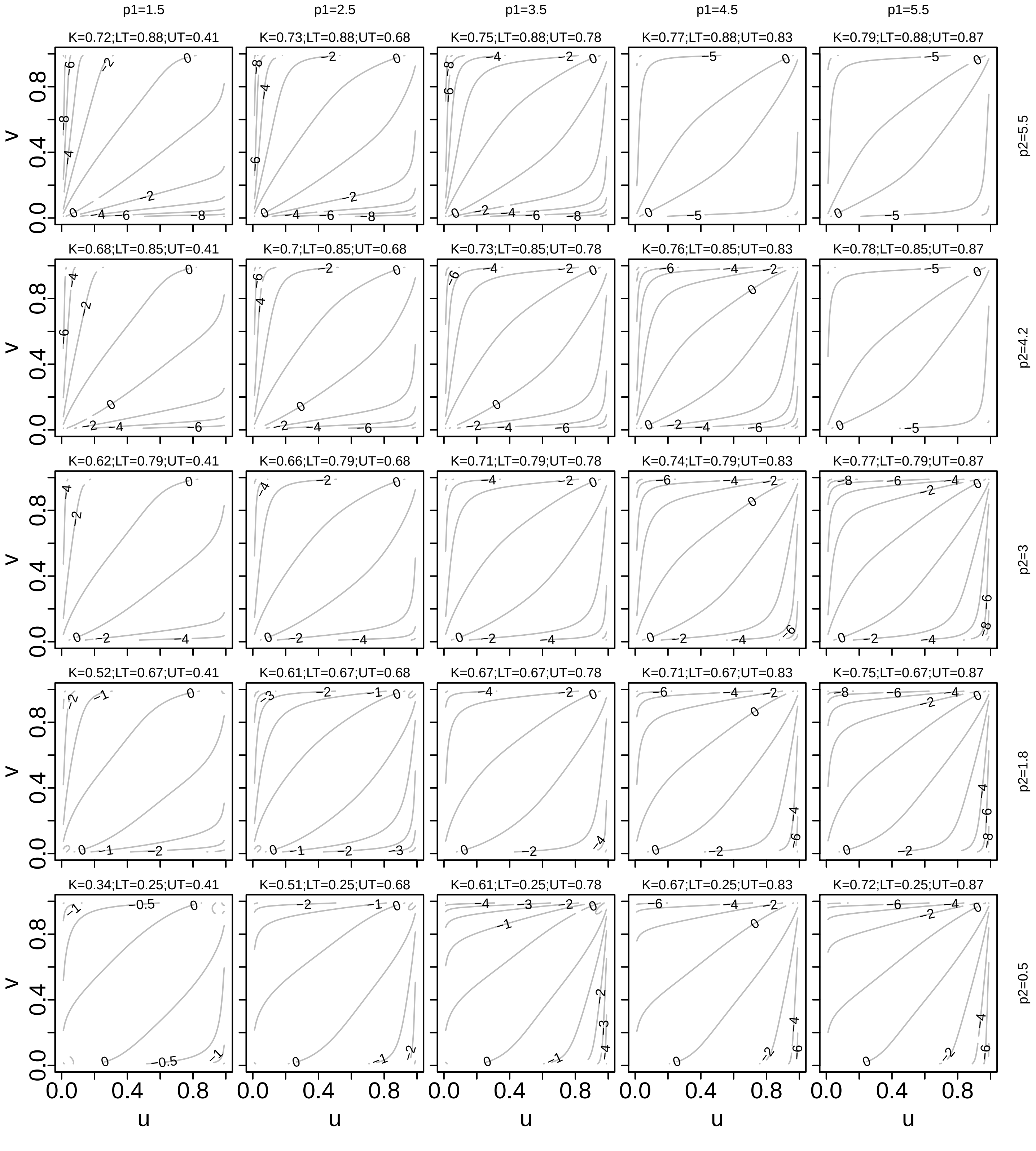

Figure S15: Log-transformed pdfs for example BB7 copulas. K is Kendall correlation; $p_1$ and $p_2$ denote the two parameters of the family (it is a two-parameter family); and LT and UT are the measures of lower- and upper-tail dependence, respectively. The parameter ranges for the family are $p_1 \in [1, \infty)$ and $p_2 \in (0, \infty)$, lower-tail dependence is $2^{-1/p_2}$ and upper-tail dependence is $2 - 2^{1/p_1}$.

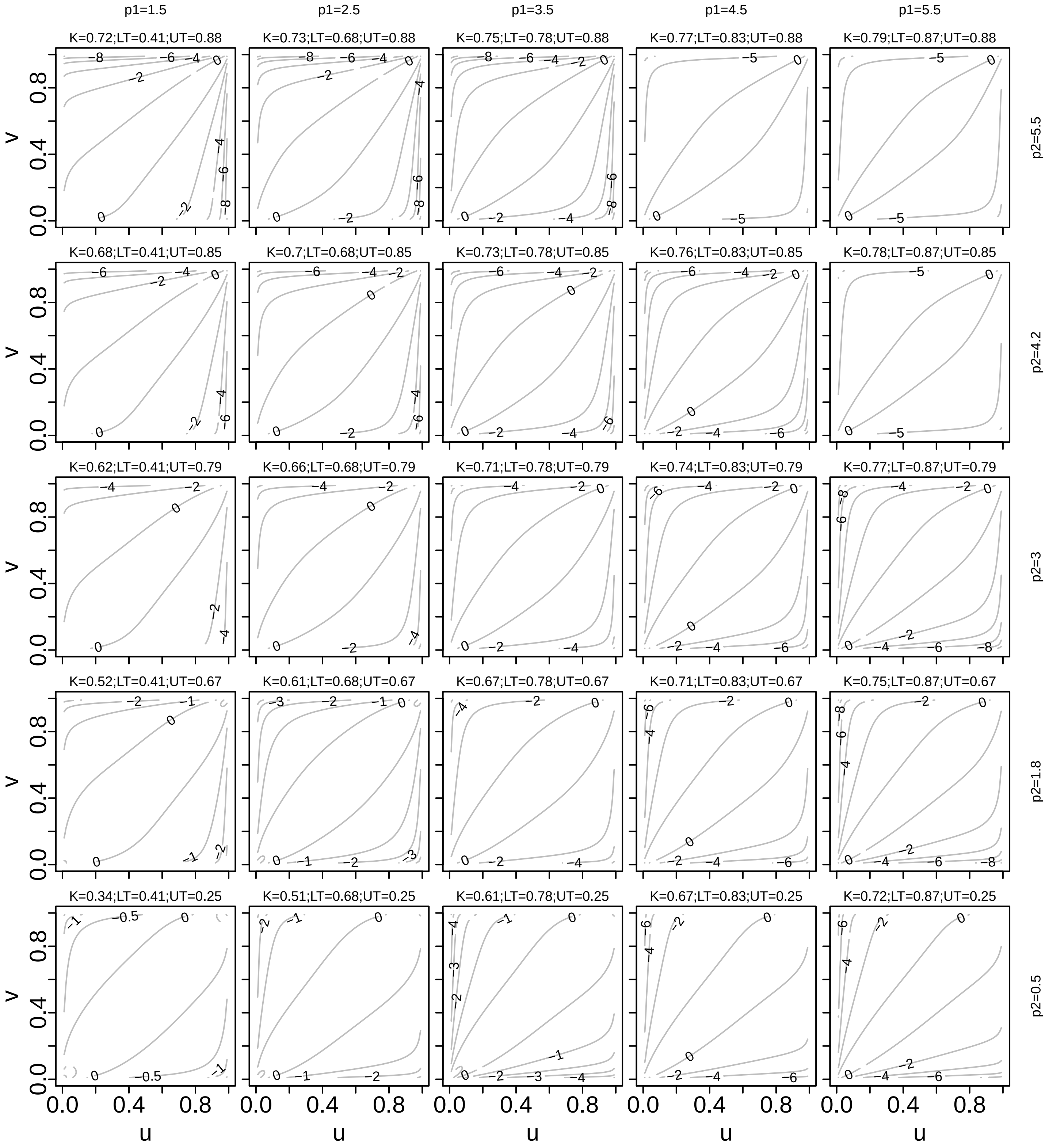

Figure S16: Log-transformed pdfs for example survival BB7 copulas. K is Kendall correlation; $p_1$ and $p_2$ denote the two parameters of the family (it is a two-parameter family); and LT and UT are the measures of lower- and upper-tail dependence, respectively. The parameter ranges for the family are $p_1 \in [1, \infty)$ and $p_2 \in (0, \infty)$, upper-tail dependence is $2^{-1/p_2}$ and lower-tail dependence is $2 - 2^{1/p_1}$.

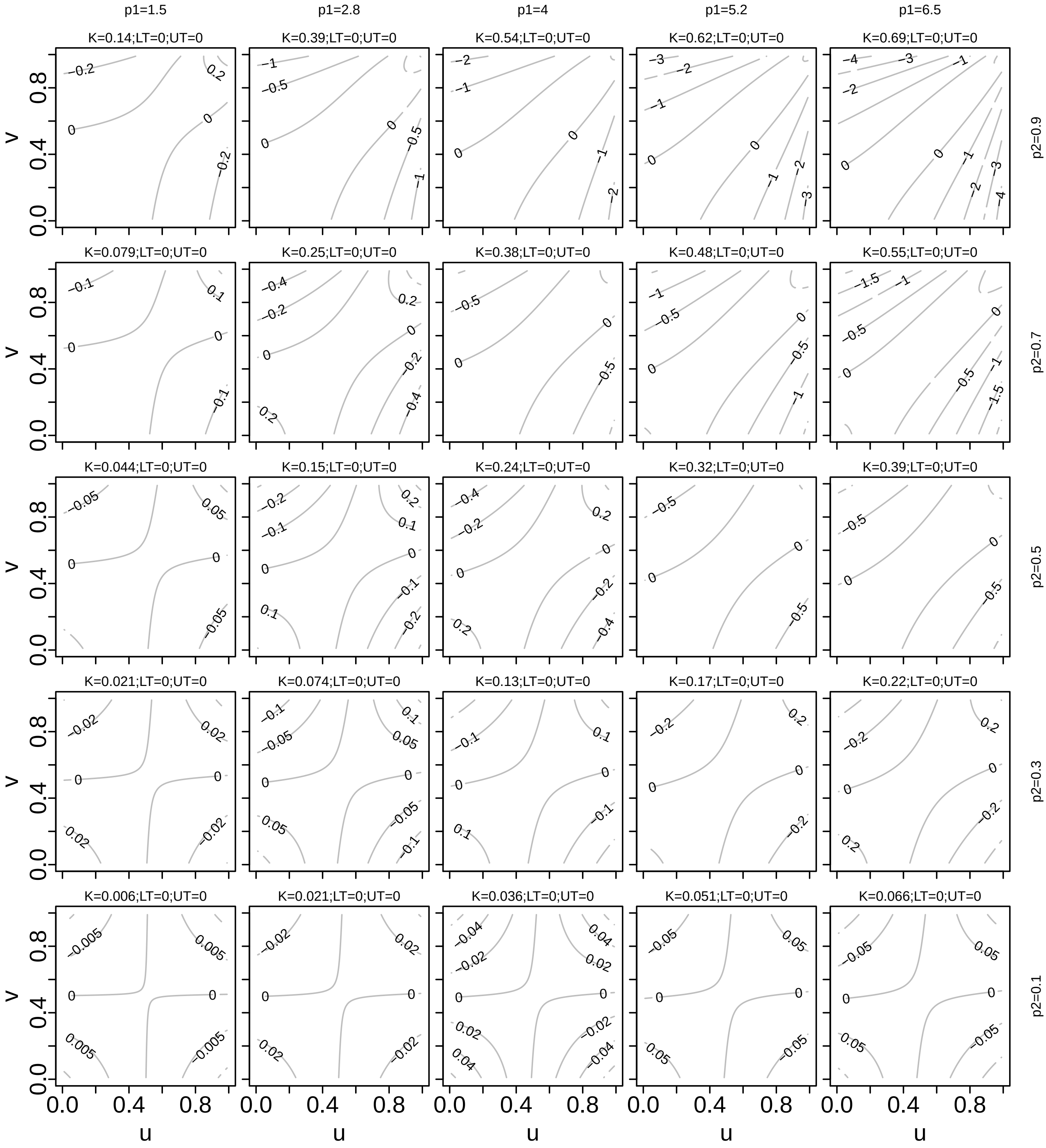

Figure S17: Log-transformed pdfs for example BB8 copulas. K is Kendall correlation; $p_1$ and $p_2$ denote the two parameters of the family (it is a two-parameter family); and LT and UT are the measures of lower- and upper-tail dependence, respectively. The parameter ranges for the family are $p_1 \in [1, \infty)$ and $p_2 \in (0, 1]$, lower-tail dependence is 0 and upper-tail dependence is $2 - 2^{1/p_1}$ of $p_2 = 1$ and 0 otherwise.

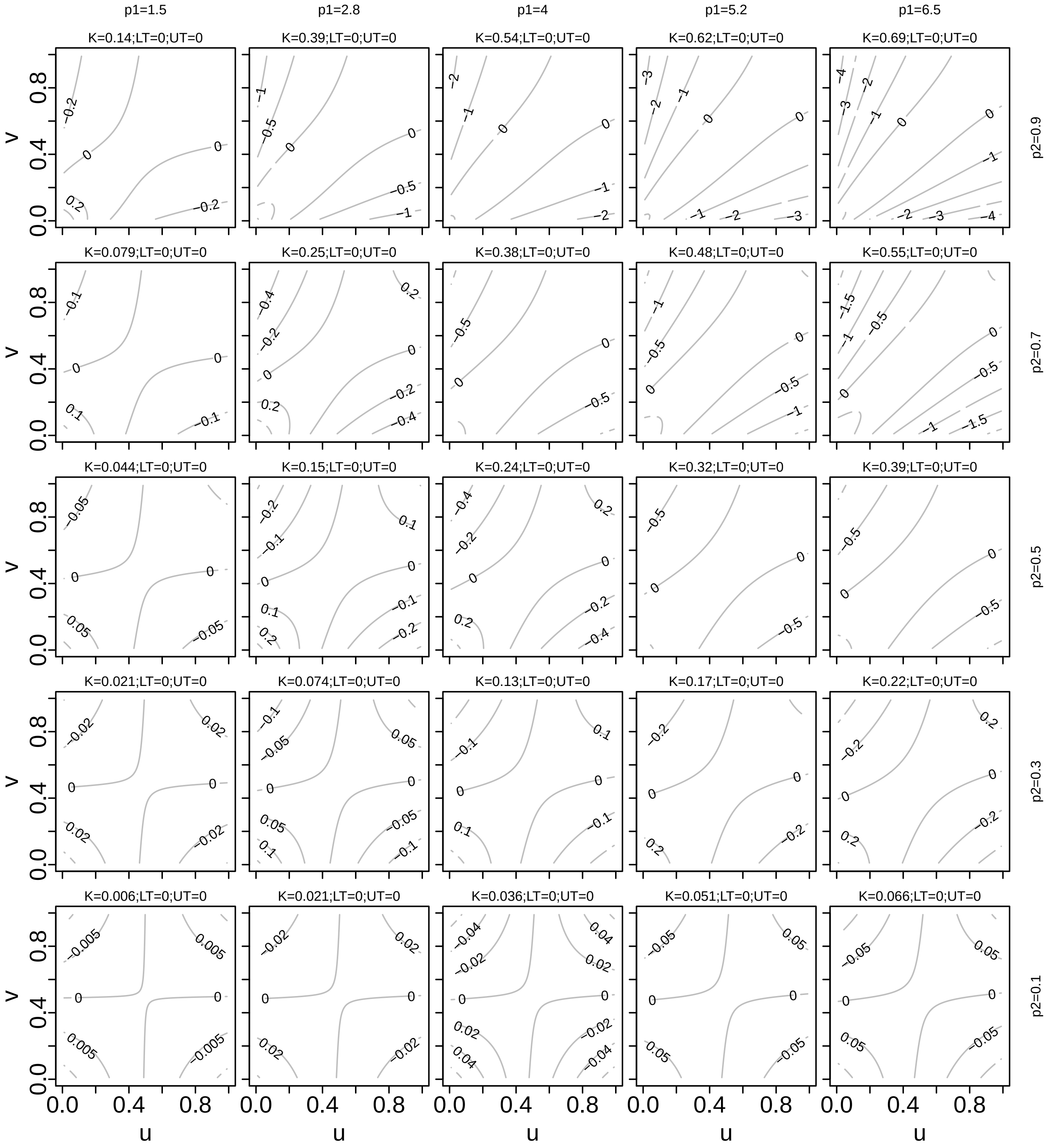

Figure S18: Log-transformed pdfs for example survival BB8 copulas. K is Kendall correlation; $p_1$ and $p_2$ denote the two parameters of the family (it is a two-parameter family); and LT and UT are the measures of lower- and upper-tail dependence, respectively. The parameter ranges for the family are $p_1 \in [1, \infty)$ and $p_2 \in (0, 1]$, upper-tail dependence is 0 and lower-tail dependence is $2 - 2^{1/p_1}$ of $p_2 = 1$ and 0 otherwise.

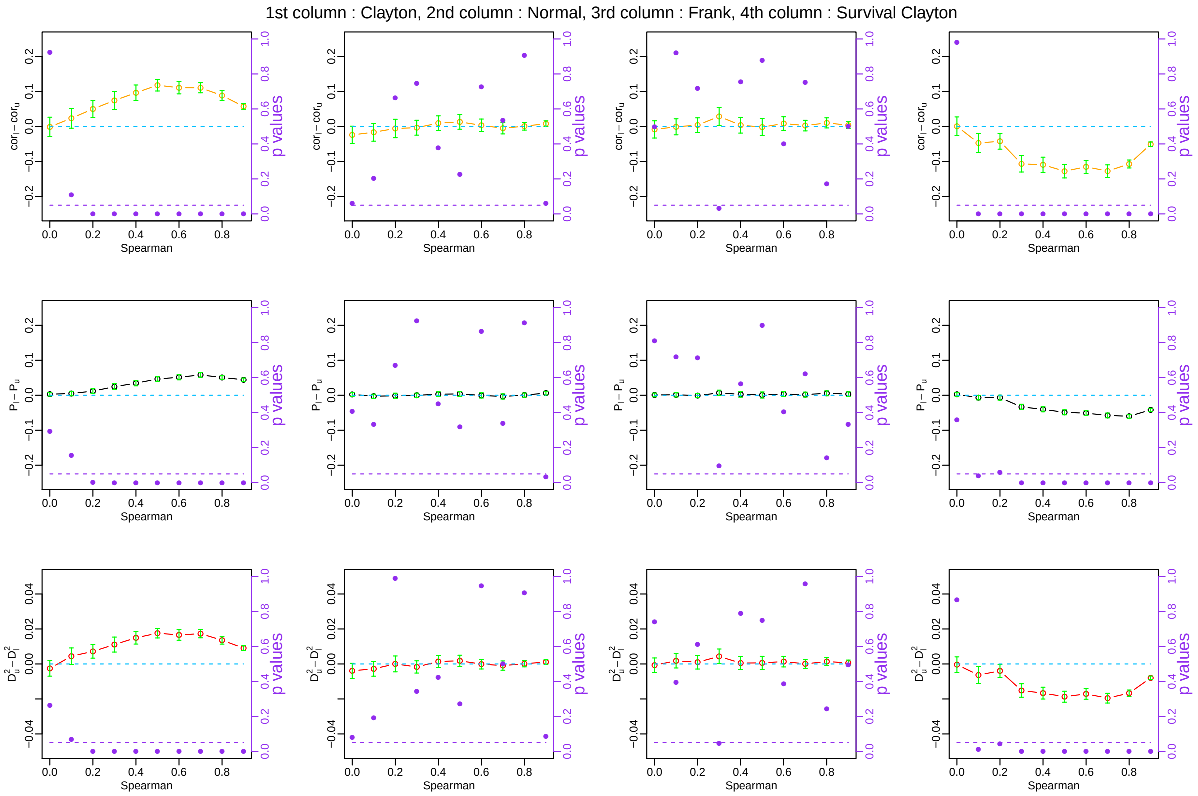

Figure S19: Results of tests of whether $\mathrm{cor}_l - \mathrm{cor}_u$, $\mathrm{P}_l - \mathrm{P}_u$, and $\mathrm{D}_u^2 - \mathrm{D}_l^2$ reveal the known asymmetry of tail dependence of the Clayton and survival Clayton copulas, and the known symmetry of tail dependence of the normal and Frank copulas. See Appendix S5 for details. $n = 35$.

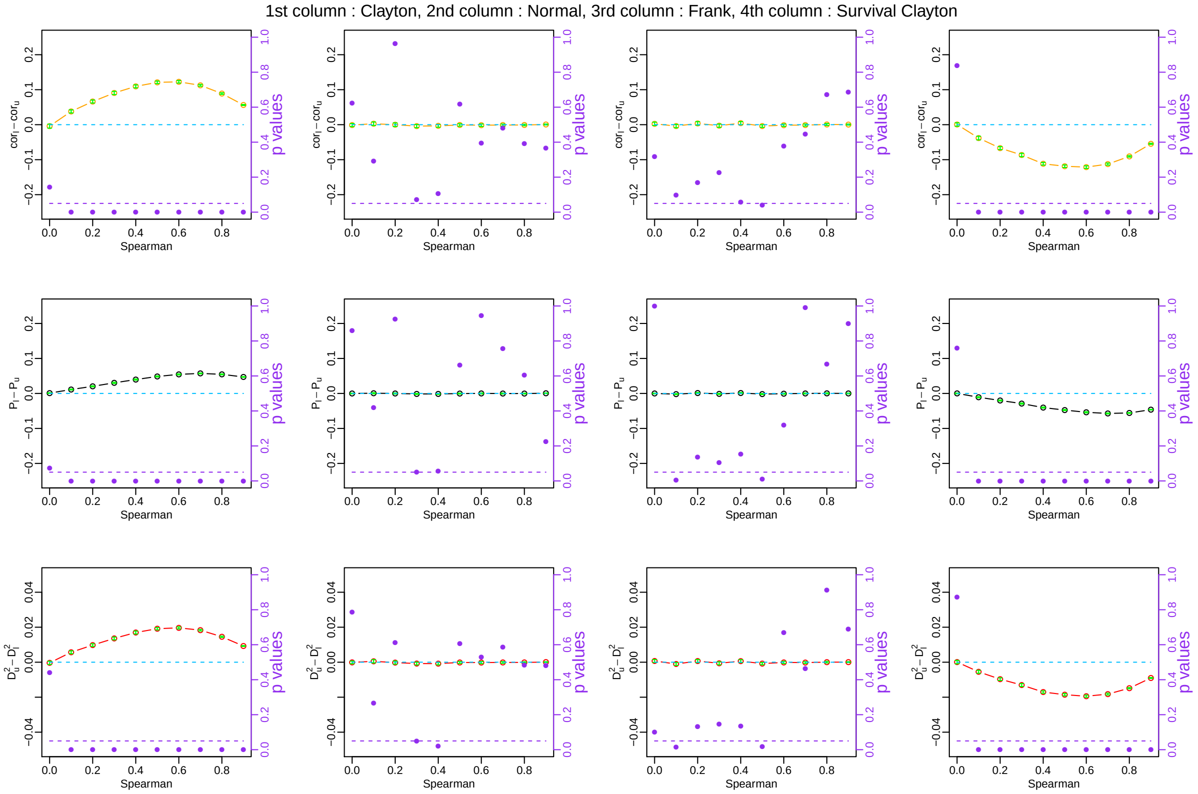

Figure S20: Results of tests of whether $\mathrm{cor}_l - \mathrm{cor}_u$, $\mathrm{P}_l - \mathrm{P}_u$, and $\mathrm{D}_u^2 - \mathrm{D}_l^2$ reveal the known asymmetry of tail dependence of the Clayton and survival Clayton copulas, and the known symmetry of tail dependence of the normal and Frank copulas. See Appendix S5 for details. $n = 1000$.

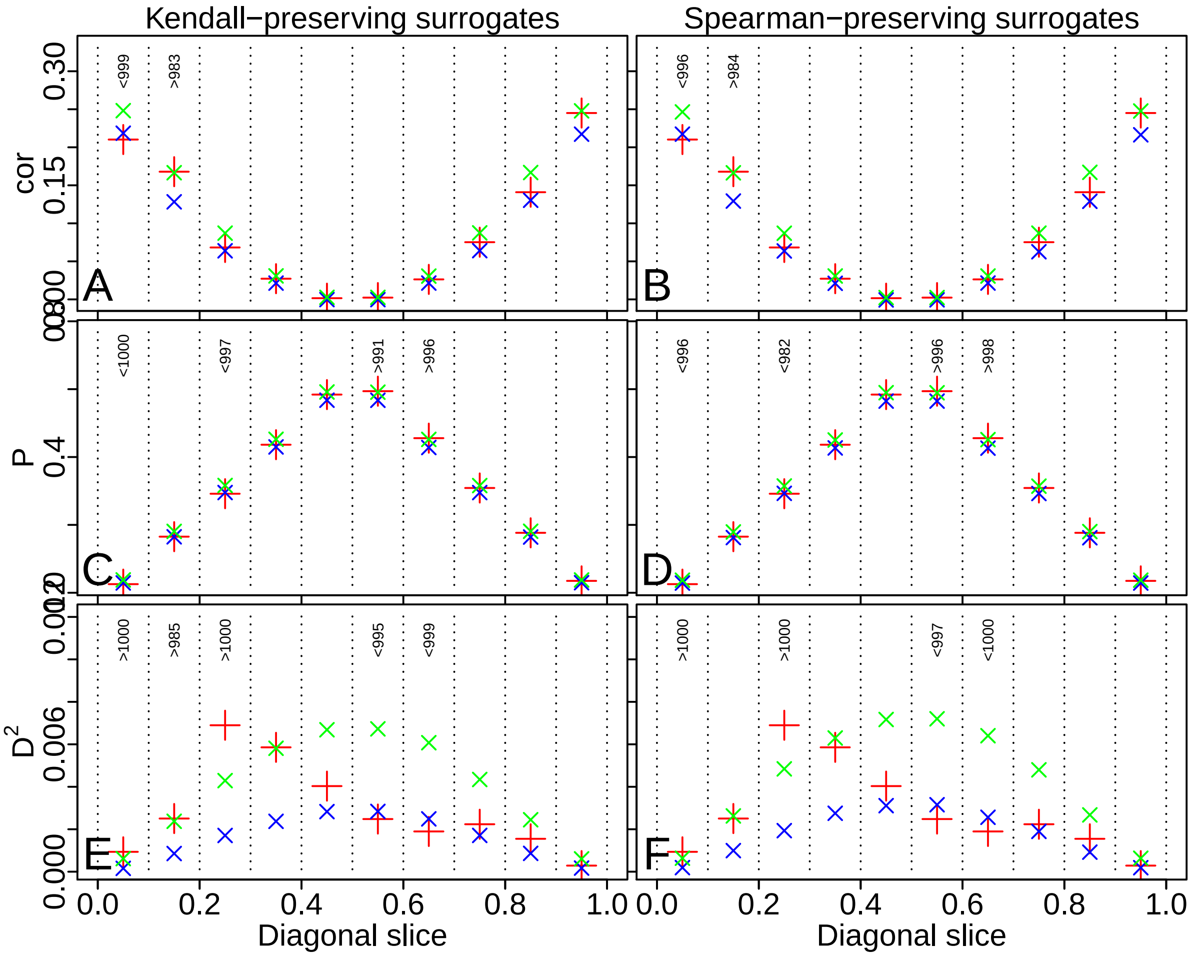

Figure S21: Nonparametric tests for tail dependence and other deviations from normal copula structure, for bird body mass and BMR data. Same format as Fig. 9, see caption of that figure.

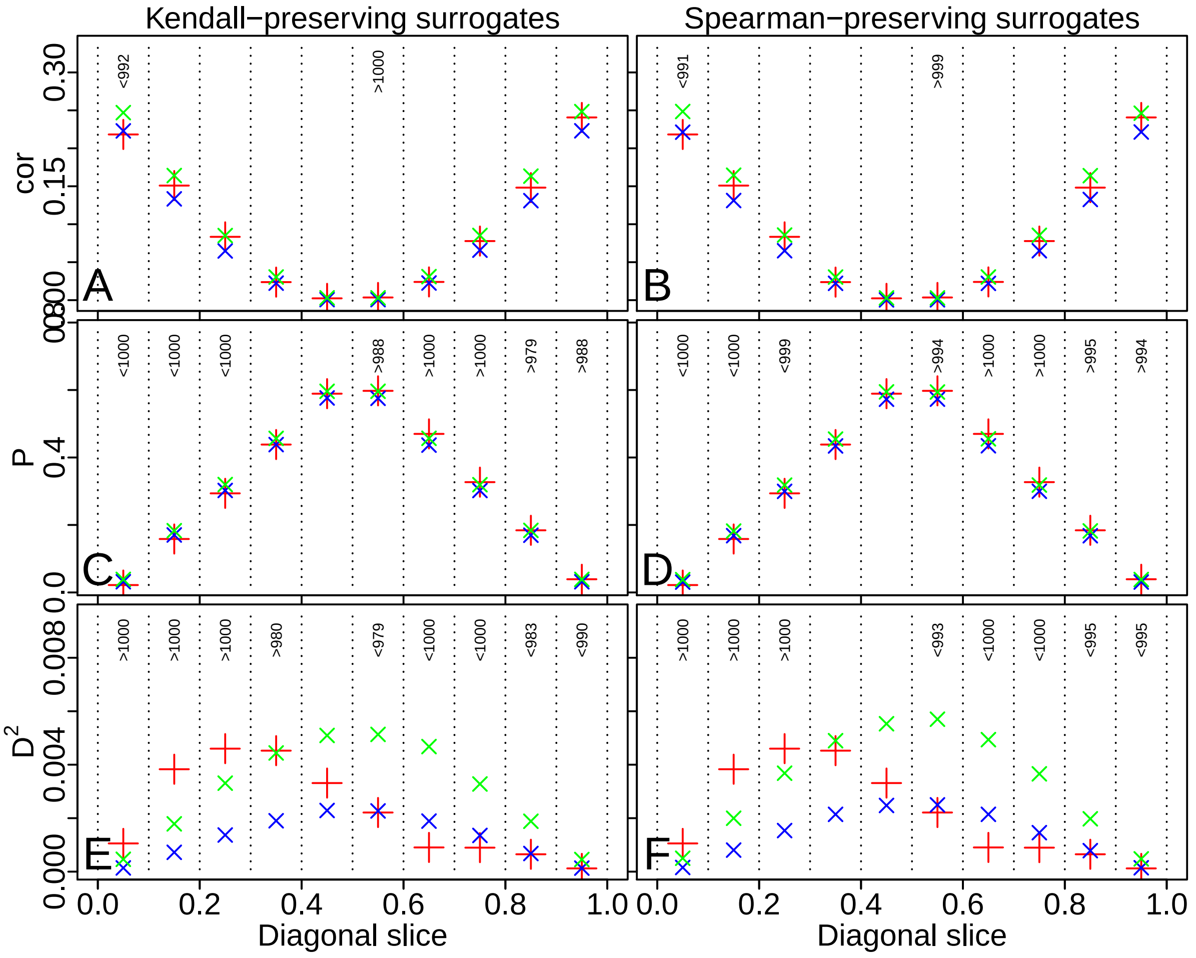

Figure S22: Nonparametric tests for tail dependence and other deviations from normal copula structure, for mammal body mass and BMR data. Same format as Fig. 9, see caption of that figure.

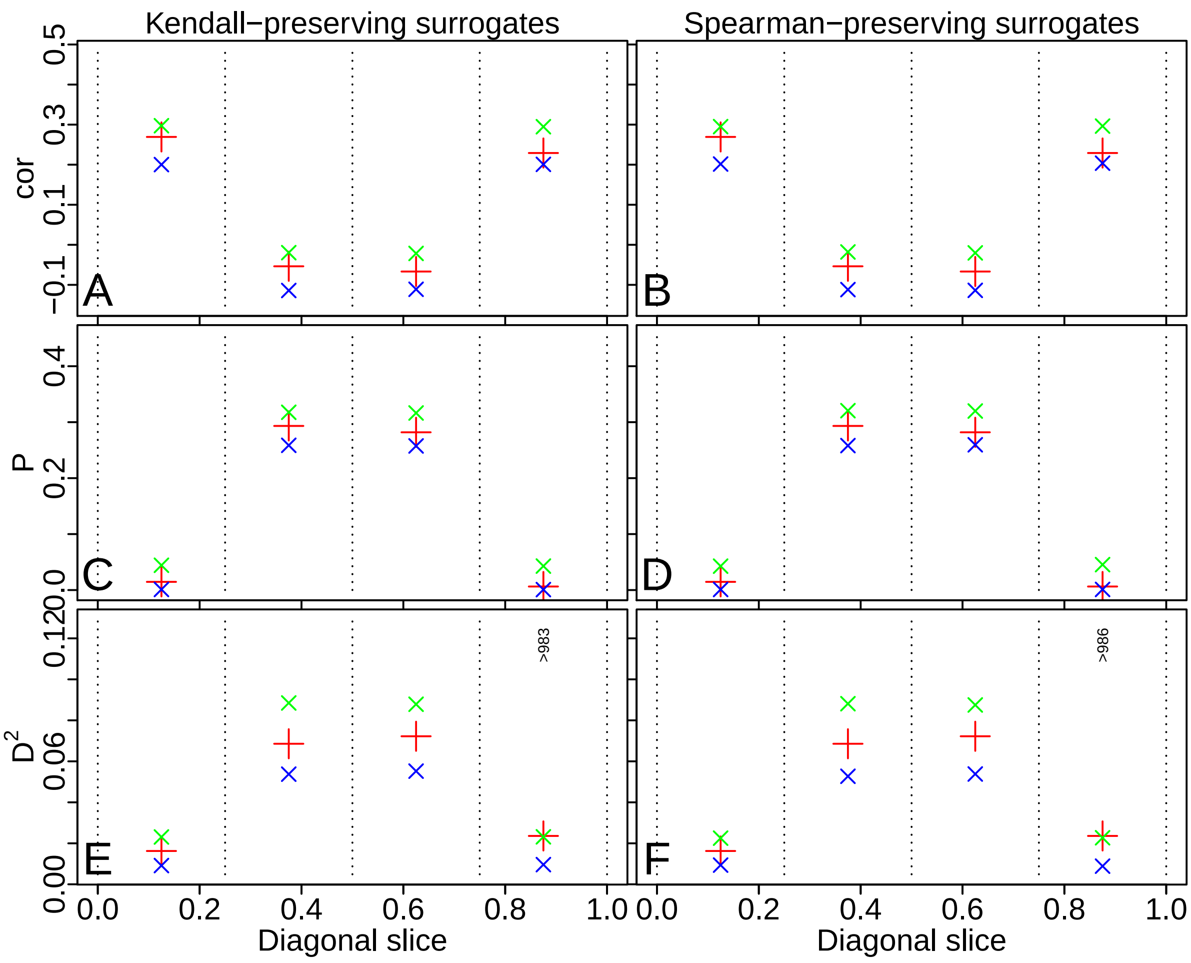

Figure S23: Nonparametric tests for tail dependence and other deviations from normal copula structure, for Cedar Creek data, year 2000. As described in the main text, the values of the statistics $\text{cor}_{l_b,u_b}$, $\text{P}_{l_b,u_b}$ and $\text{D}^2_{l_b,u_b}$ for real data (red crosses) were compared to distributions of their values on 1000 Kendall- or Spearman-preserving normal surrogates of the data (blue and green x's show 0.025 and 0.975 quantiles), separately in two comparisons for each of the ranges $(l_b, u_b) = (0, 0.25), \ldots, (0.75, 1)$. If the red cross was outside the range given by the x's, text at the top of panels indicates the number of surrogate values the real-data value was greater than or less than. For instance, a value $> N$ (respectively, $< N$) means the value of the statistic on real data was greater than (respectively, less than) its value on $N$ surrogates. When the statistic was greater than 975 or less than 975 surrogate values, it indicates significance (95% confidence level). When cor or P values (respectively, $\text{D}^2$ values) were greater than surrogates, it means dependence in that part of the distributions was stronger than (respectively, weaker than) expected from a normal-copula null hypothesis.

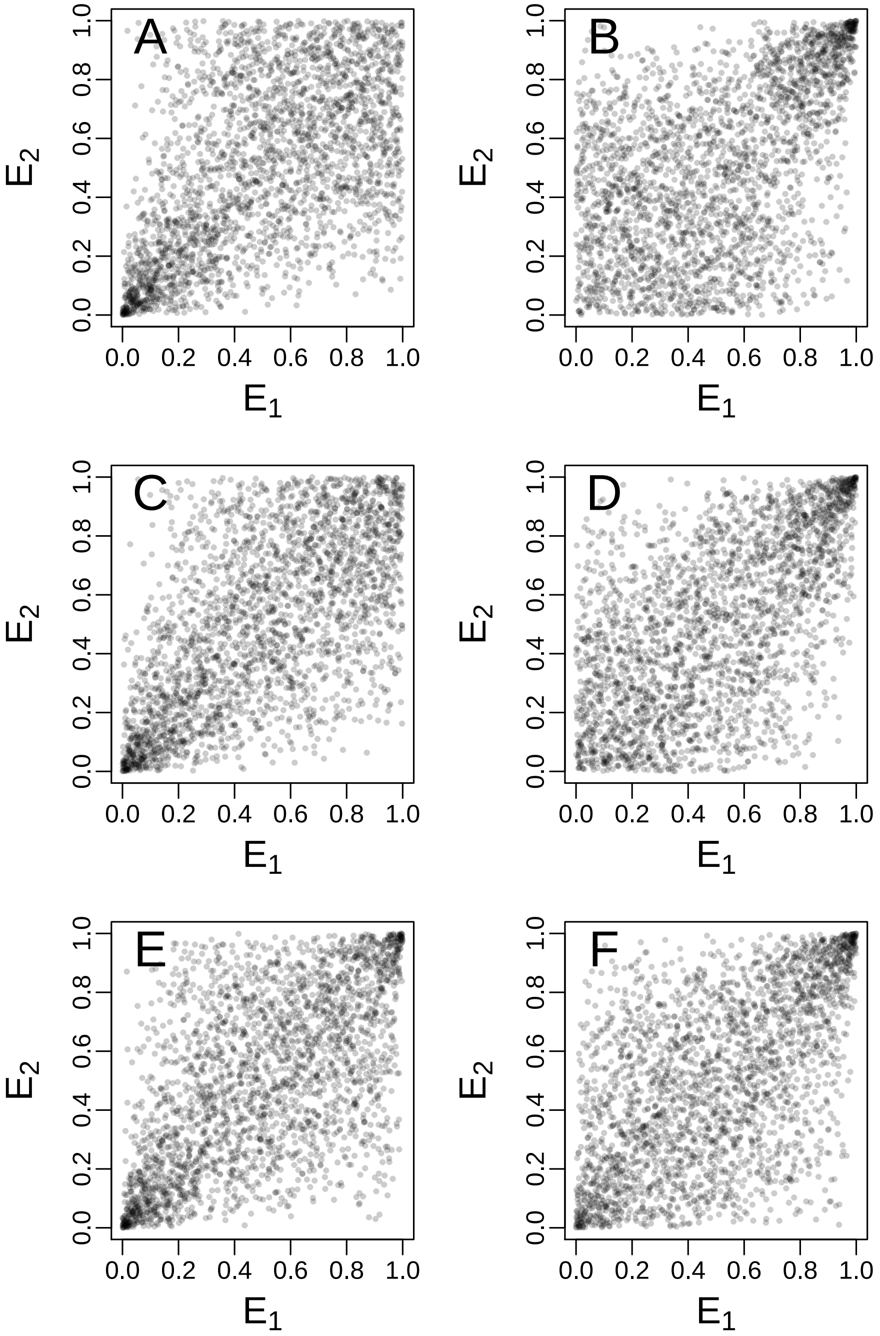

Figure S24: Analogous to Fig. 10, but using $b = -0.1$ (A-B), $b = 0.5$ (C-D) and $b = -0.5$ (E-F).

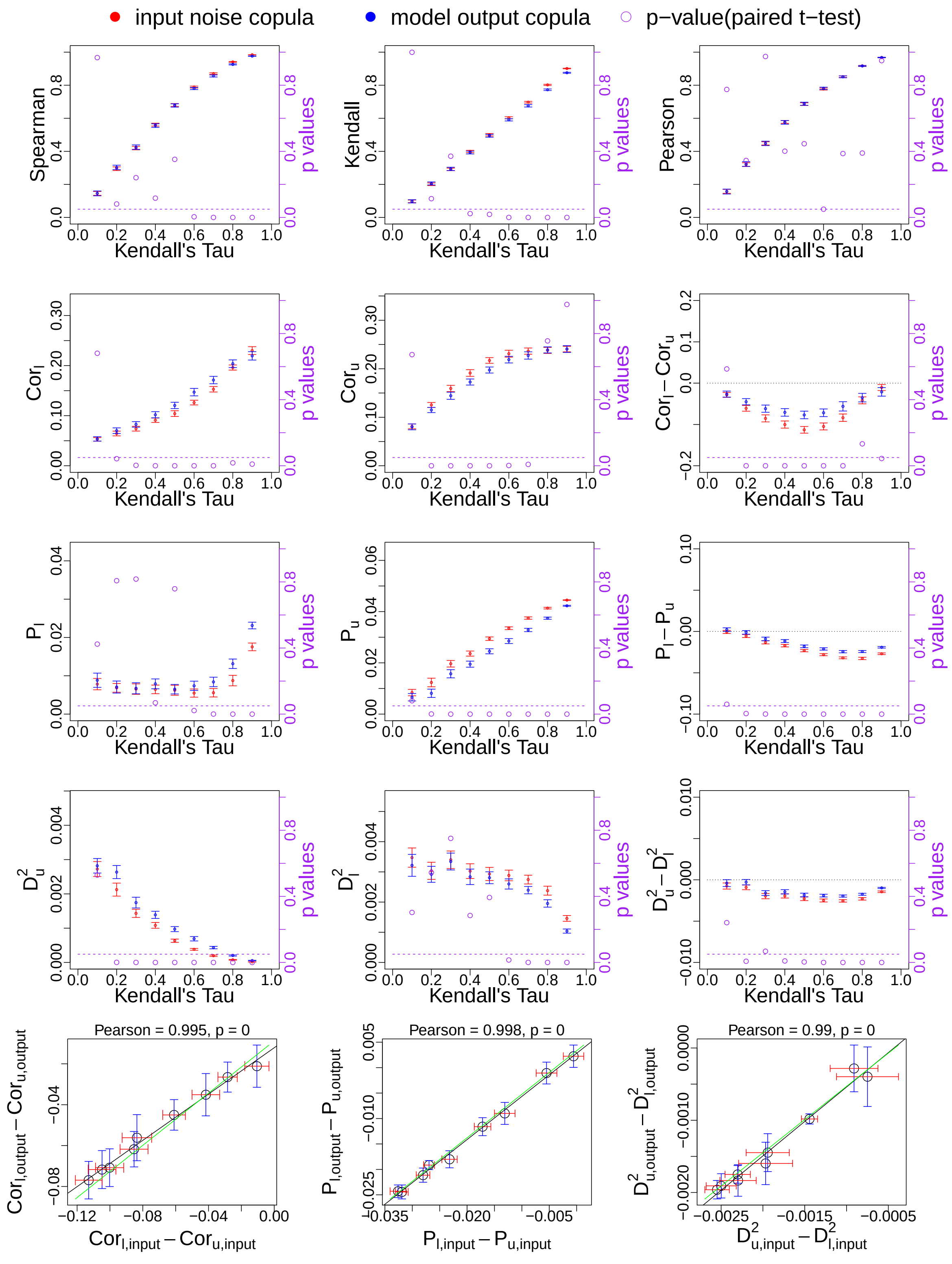

Figure S25: Comparison of copula structure of noise inputs and model outputs, AR(1) model, survival Clayton copula. Same format as Fig. 11, see caption of that figure for details.

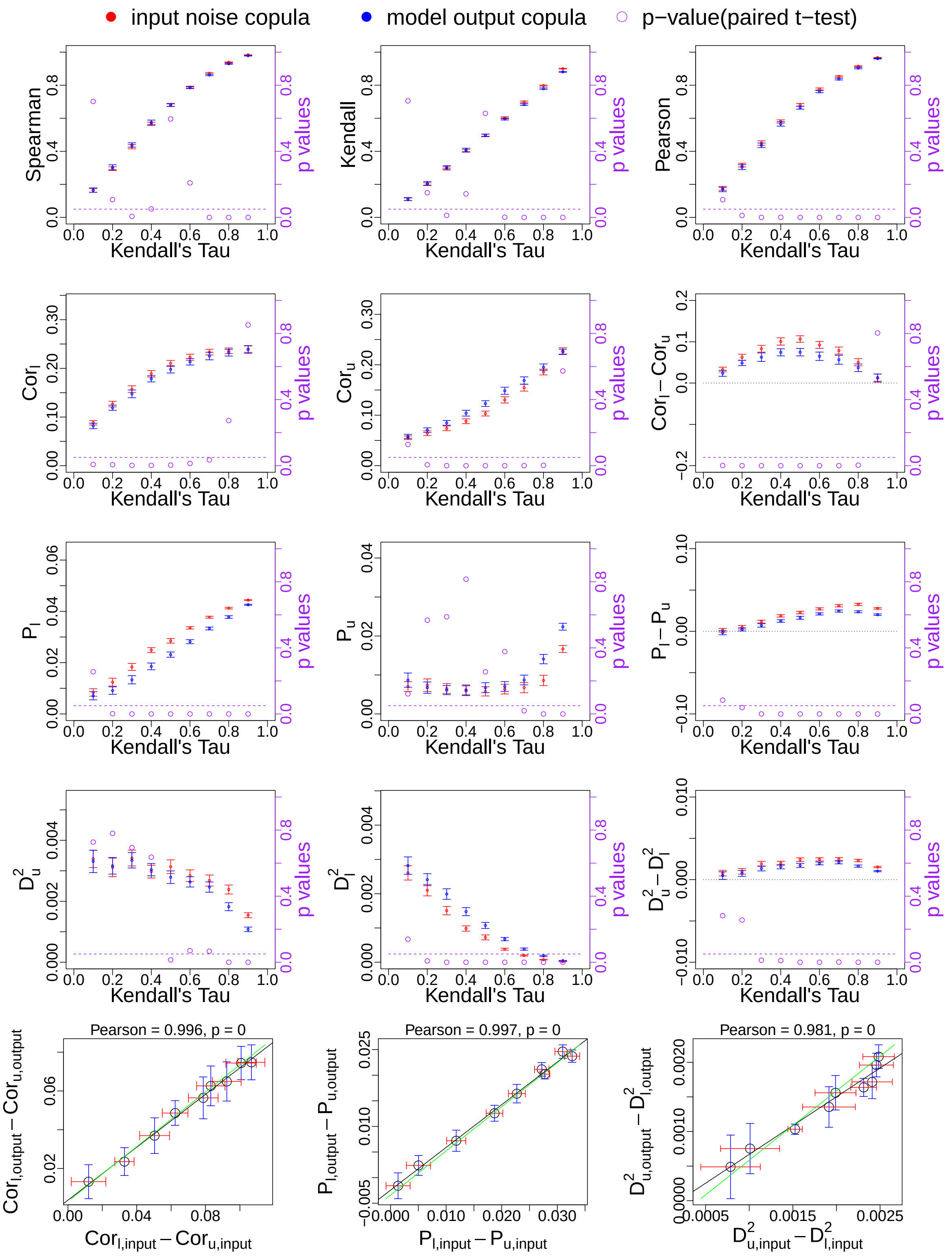

Figure S26: Comparison of copula structure of noise inputs and model outputs, stochastic Ricker model, weak noise, Clayton copula. Same format as Fig. 11, see caption of that figure for details.

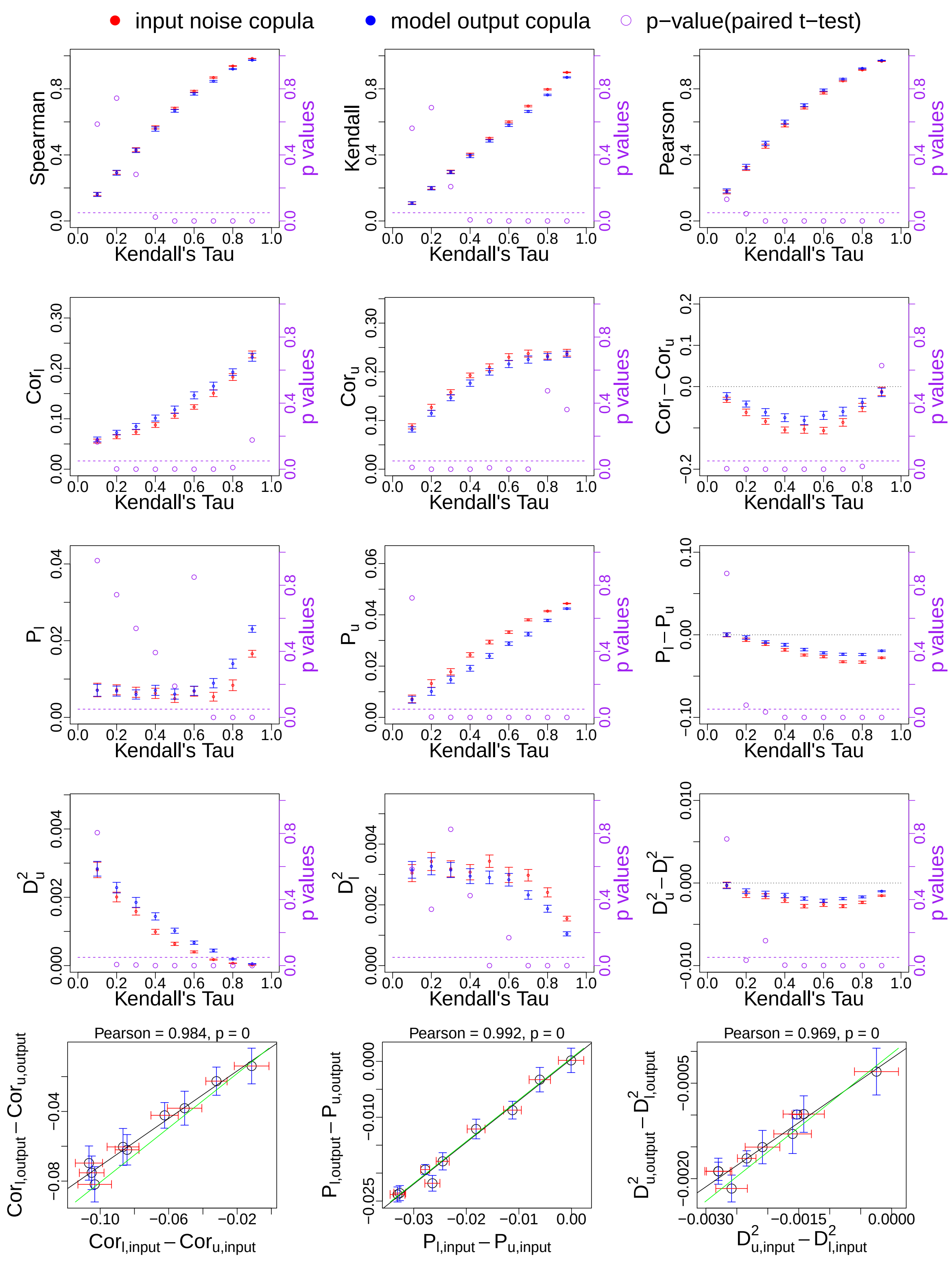

Figure S27: Comparison of copula structure of noise inputs and model outputs, stochastic Ricker model, weak noise, survival Clayton copula. Same format as Fig. 11, see caption of that figure for details.

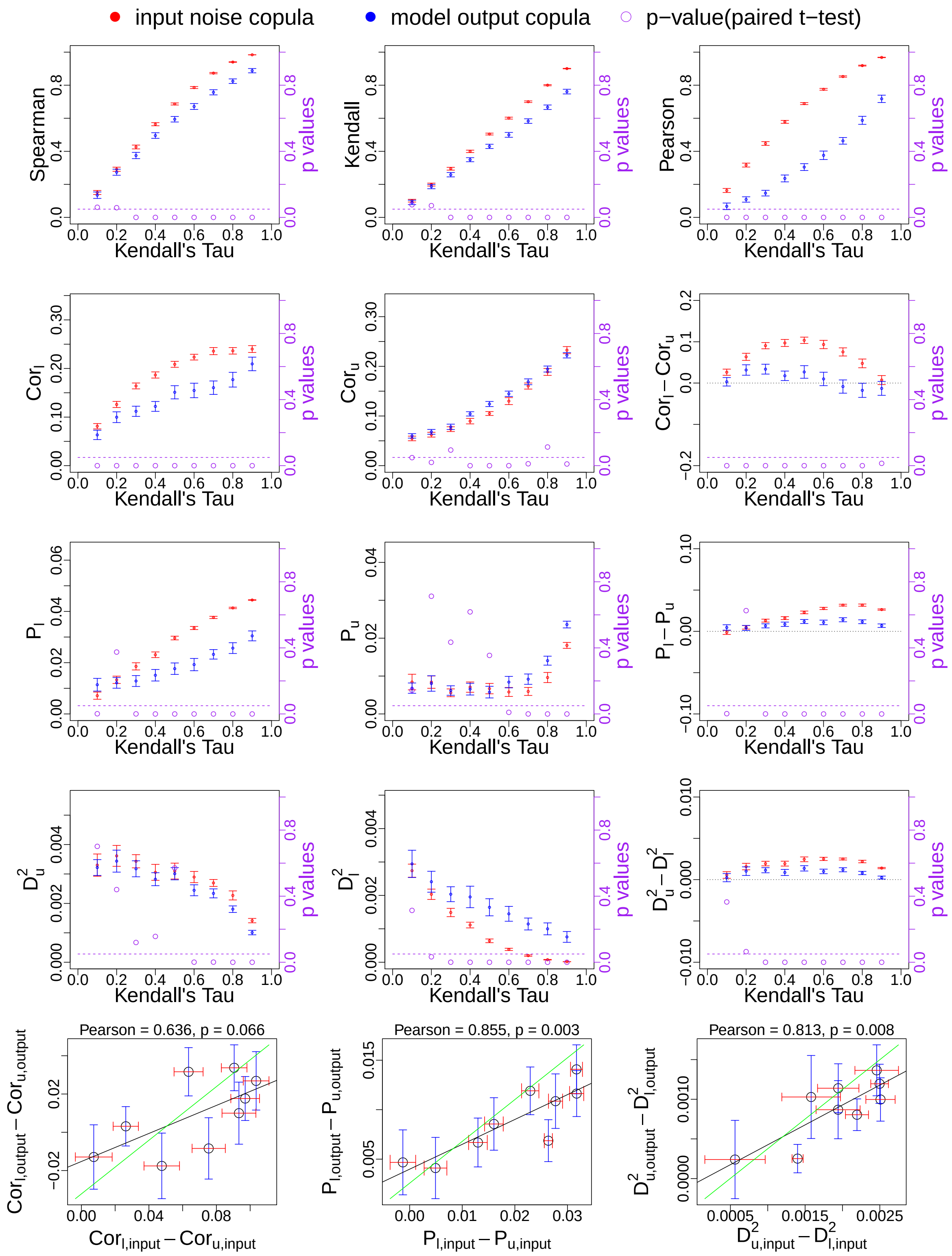

Figure S28: Comparison of copula structure of noise inputs and model outputs, stochastic Ricker model, strong noise, Clayton copula. Same format as Fig. 11, see caption of that figure for details.

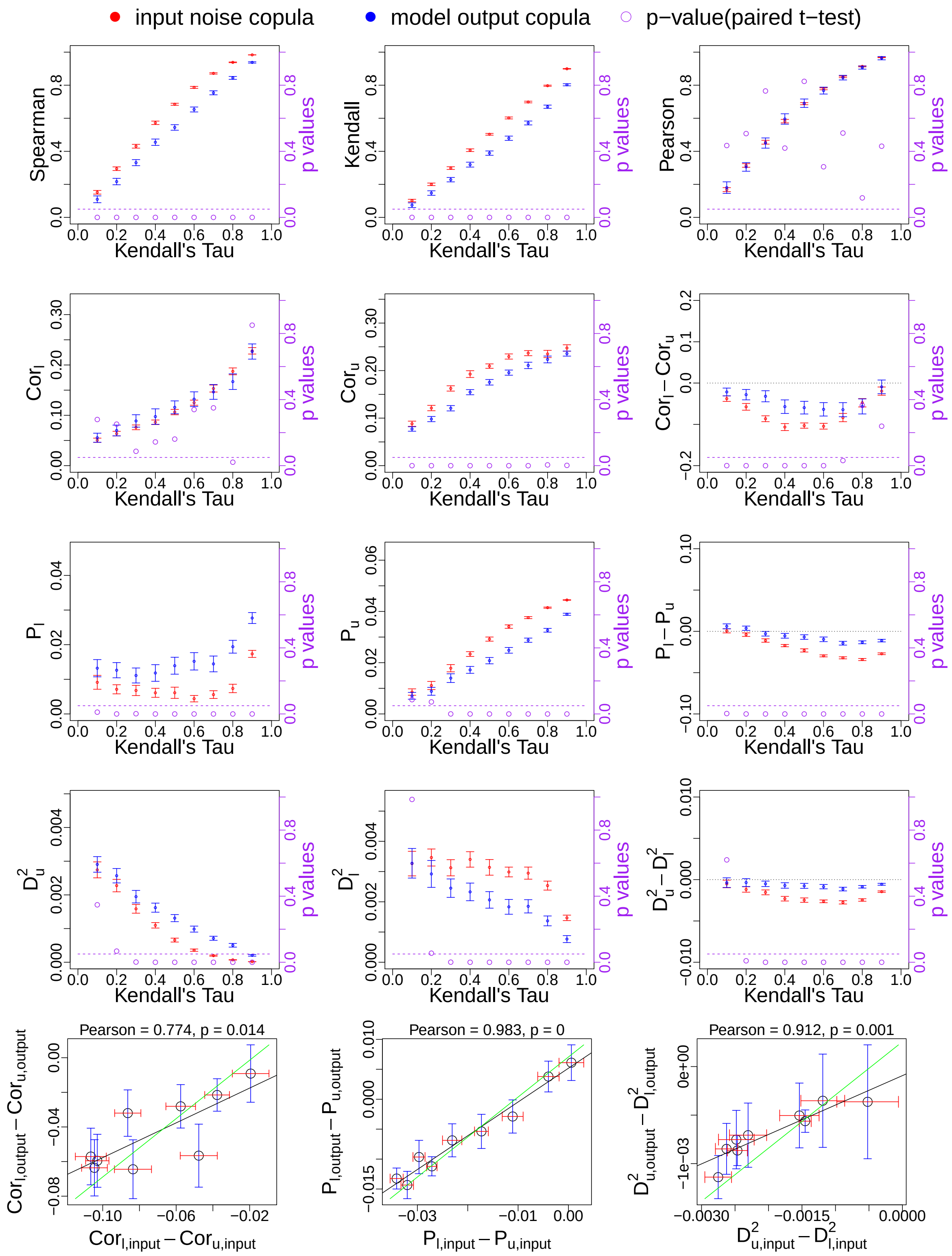

Figure S29: Comparison of copula structure of noise inputs and model outputs, stochastic Ricker model, strong noise, survival Clayton copula. Same format as Fig. 11, see caption of that figure for details.

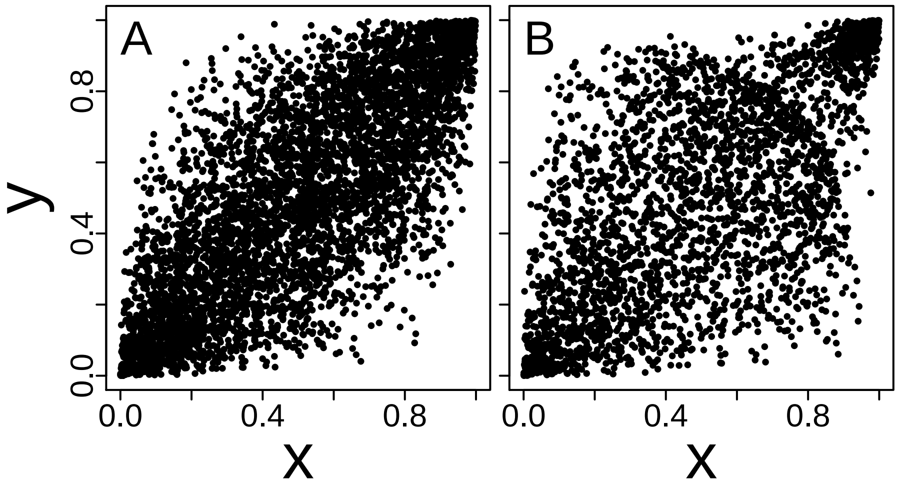

Figure S30: Missing data can influence perceived copula structure. Points were randomly removed from the upper right portion of the copula dataset pictured in A, and then remaining values were re-ranked to make B. Thus removing data in a non-random fashion can influence copula structure and tail dependence. See Appendix S13 for details.

Table S1: Species names of 20 aphids for which data were available

| Common name | Latin binomial |
|---|---|
| Apple grass aphid | *Rhopalosiphum insertum* |
| Bird cherry oat aphid | *Rhopalosiphum padi* |
| Black bean aphid | *Aphis fabae* |
| Blackberry cereal aphid | *Sitobion fragariae* |
| Blackcurrant sowthistle aphid | *Hyperomyzus lactucae* |
| Corn leaf aphid | *Rhopalosiphum maidis* |
| Currant lettuce aphid | *Nasonovia ribisnigri* |
| Damson hop aphid | *Phorodon humuli* |
| Grain aphid | *Sitobion avenae* |
| Green spruce aphid | *Elatobium abietinum* |
| Leaf-curling plum aphid | *Brachycaudus helichrysi* |
| Mealy cabbage aphid | *Brevicoryne brassicae* |
| Mealy plum aphid | *Hyalopterus pruni* |
| Pea aphid | *Acyrthosiphon pisum* |
| Peach potato aphid | *Myzus persicae* |
| Potato aphid | *Macrosiphum euphorbiae* |
| Rose grain aphid | *Metopolophium dirhodum* |
| Shallot aphid | *Myzus ascalonicus* |
| Sycamore aphid | *Drepanosiphum platanoidis* |
| Willow carrot aphid | *Cavariella aegopodii* |

Table S2: Species names of 22 plankton taxa for which data were available

| Plankton taxon name |
| --- |
| *Calanus stages I-IV* |
| *Para-pseudocalanus spp.* |
| *Temora longicornis* |
| *Acartia sp.* |
| *Centropages typicus* |
| *Oithona sp.* |
| Echinoderm larvae |
| *Calanus finmarchicus* |
| *Calanus helgolandicus* |
| *Metridia lucens* |
| Decapoda larvae |
| Euphausiids |
| *Thalassiosira sp.* |
| *Rhizosolenia styliformis* |
| *Ceratium fusus* |
| *Ceratium furca* |
| *Ceratium tripos* |
| *Ceratium macroceros* |
| *Proboscia alata* |
| *Pseudocalanus sp.* |
| *Pseudo-nitzschia delicatissima* |
| *Pseudo-nitzschia seriata* |

Table S3: Results of fitting our 16 copula families to the bird body mass and BMR dataset. Analogous to Fig. 2, see caption of that figure for details.

| Copula | AIC | AICw | BIC | BICw | LT | UT |
|---|---|---|---|---|---|---|
| G | -1447.36 | 0.58 | -1443.08 | 0.92 | 0.0000 | 0.8770 |
| BB6 | -1445.35 | 0.21 | -1436.79 | 0.04 | 0.0000 | 0.8771 |
| BB1 | -1445.34 | 0.21 | -1436.79 | 0.04 | 0.0000 | 0.8770 |
| SBB1 | -1417.77 | 0.00 | -1409.22 | 0.00 | 0.7009 | 0.8898 |
| F | -1370.90 | 0.00 | -1366.62 | 0.00 | 0.0000 | 0.0000 |
| N | -1336.77 | 0.00 | -1332.49 | 0.00 | 0.0000 | 0.0000 |
| SC | -1334.40 | 0.00 | -1330.12 | 0.00 | 0.0000 | 0.9125 |
| J | -1333.69 | 0.00 | -1329.41 | 0.00 | 0.0000 | 0.9133 |
| BB7 | -1306.66 | 0.00 | -1298.10 | 0.00 | 0.7305 | 0.8513 |
| SBB7 | -1301.78 | 0.00 | -1293.22 | 0.00 | 0.7048 | 0.8909 |
| BB8 | -1271.15 | 0.00 | -1262.59 | 0.00 | 0.0000 | 0.8775 |
| SG | -1204.58 | 0.00 | -1200.30 | 0.00 | 0.8486 | 0.0000 |
| SBB6 | -1202.42 | 0.00 | -1193.86 | 0.00 | 0.8486 | 0.0000 |
| SBB8 | -1051.19 | 0.00 | -1042.64 | 0.00 | 0.0000 | 0.0000 |
| SJ | -913.23 | 0.00 | -908.95 | 0.00 | 0.8647 | 0.0000 |
| C | -908.59 | 0.00 | -904.31 | 0.00 | 0.8611 | 0.0000 |

Table S4: Results of fitting our 16 copula families to the mammal body mass and BMR dataset. Analogous to Fig. 2, see caption of that figure for details.

| Copula | AIC | AICw | BIC | BICw | LT | UT |
|---|---|---|---|---|---|---|
| G | -1854.31 | 0.60 | -1849.85 | 0.93 | 0.0000 | 0.8897 |
| BB6 | -1853.50 | 0.40 | -1844.59 | 0.07 | 0.0000 | 0.8981 |
| BB1 | -1843.87 | 0.00 | -1834.95 | 0.00 | 0.0000 | 0.8775 |
| SBB1 | -1821.03 | 0.00 | -1812.11 | 0.00 | 0.6182 | 0.9168 |
| F | -1775.90 | 0.00 | -1771.44 | 0.00 | 0.0000 | 0.0000 |
| J | -1772.26 | 0.00 | -1767.80 | 0.00 | 0.0000 | 0.9278 |
| SC | -1769.98 | 0.00 | -1765.53 | 0.00 | 0.0000 | 0.9274 |
| N | -1678.76 | 0.00 | -1674.30 | 0.00 | 0.0000 | 0.0000 |
| SBB7 | -1668.07 | 0.00 | -1659.16 | 0.00 | 0.0014 | 0.8909 |
| BB8 | -1625.27 | 0.00 | -1616.35 | 0.00 | 0.0000 | 0.0000 |
| BB7 | -1619.64 | 0.00 | -1610.72 | 0.00 | 0.7152 | 0.8513 |
| SG | -1460.21 | 0.00 | -1455.75 | 0.00 | 0.8507 | 0.0000 |
| SBB6 | -1457.98 | 0.00 | -1449.06 | 0.00 | 0.8507 | 0.0000 |
| SBB8 | -1262.61 | 0.00 | -1253.70 | 0.00 | 0.0000 | 0.0000 |
| SJ | -1072.97 | 0.00 | -1068.52 | 0.00 | 0.8601 | 0.0000 |
| C | -1070.43 | 0.00 | -1065.97 | 0.00 | 0.8558 | 0.0000 |

Table S5: Results of fitting our 16 copula families to the Cedar Creek dataset, year 2000. Analogous to Fig. 2, see caption of that figure for details.

| Copula | AIC | AICw | BIC | BICw | LT | UT |
|---|---|---|---|---|---|---|
| F | -22.68 | 0.38 | -19.56 | 0.54 | 0.0000 | 0.0000 |
| SBB8 | -21.64 | 0.22 | -15.39 | 0.07 | 0.0000 | 0.0000 |
| N | -20.42 | 0.12 | -17.29 | 0.17 | 0.0000 | 0.0000 |
| BB8 | -19.38 | 0.07 | -13.13 | 0.02 | 0.0000 | 0.0000 |
| SG | -19.07 | 0.06 | -15.95 | 0.09 | 0.2908 | 0.0000 |
| C | -18.17 | 0.04 | -15.04 | 0.06 | 0.2573 | 0.0000 |
| SBB1 | -17.07 | 0.02 | -10.82 | 0.01 | 0.2905 | 0.0000 |
| SBB6 | -17.07 | 0.02 | -10.82 | 0.01 | 0.2912 | 0.0000 |
| BB1 | -16.60 | 0.02 | -10.36 | 0.01 | 0.2044 | 0.0817 |
| BB7 | -16.21 | 0.01 | -9.96 | 0.00 | 0.2478 | 0.0334 |
| SJ | -15.57 | 0.01 | -12.45 | 0.02 | 0.3661 | 0.0000 |
| SBB7 | -14.75 | 0.01 | -8.51 | 0.00 | 0.3148 | 0.0181 |
| G | -13.14 | 0.00 | -10.01 | 0.00 | 0.0000 | 0.2625 |
| BB6 | -11.12 | 0.00 | -4.87 | 0.00 | 0.0000 | 0.2629 |
| SC | -10.09 | 0.00 | -6.97 | 0.00 | 0.0000 | 0.1723 |
| J | -6.12 | 0.00 | -3.00 | 0.00 | 0.0000 | 0.2850 |

Table S6: Nonparametric asymmetry tests against a normal-copula null model with birds, mammals and Cedar Creek data. For bird and mammal data, results should be interpreted in the same way as soil C and N results (Table 3). For Cedar Creek data, first slice ($FS$) and last slice ($LS$) stand for the ranges $(0, 0.25)$ and $(0.75, 1)$ for the lower and upper bounds.

| Data | Statistic | Kendall | Spearman |
|---|---|---|---|
| Bird masses and BMR | $\text{cor}_{FS}$-$\text{cor}_{LS}$ | <1000 | <999 |
| | $\text{P}_{FS}$-$\text{P}_{LS}$ | <1000 | <996 |
| | $\text{D}^2_{LS}$-$\text{D}^2_{FS}$ | <1000 | <999 |
| Mammal masses and BMR | $\text{cor}_{FS}$-$\text{cor}_{LS}$ | <991 | <993 |
| | $\text{P}_{FS}$-$\text{P}_{LS}$ | <1000 | <1000 |
| | $\text{D}^2_{LS}$-$\text{D}^2_{FS}$ | <1000 | <1000 |
| Cedar Creek | $\text{cor}_{FS}$-$\text{cor}_{LS}$ | >880 | >897 |
| | $\text{P}_{FS}$-$\text{P}_{LS}$ | >702 | >693 |
| | $\text{D}^2_{LS}$-$\text{D}^2_{FS}$ | >925 | >939 |

Table S7: Average values of statistics across all location pairs and confidence intervals based on spatial resampling for leaf-curling plum aphid first flight data.

| | $2.5^{th}$ quantile | Mean | $97.5^{th}$ quantile |
|---|---|---|---|
| Spearman | 0.528 | 0.569 | 0.616 |
| Kendall | 0.386 | 0.427 | 0.476 |
| $\text{cor}_l$ | 0.175 | 0.212 | 0.245 |
| $\text{cor}_u$ | 0.327 | 0.362 | 0.400 |
| $\text{P}_l$ | 0.063 | 0.074 | 0.086 |
| $\text{P}_u$ | 0.118 | 0.136 | 0.157 |
| $\text{D}_l^2$ | 0.038 | 0.042 | 0.046 |
| $\text{D}_u^2$ | 0.015 | 0.022 | 0.029 |
| $\text{cor}_l$ - $\text{cor}_u$ | -0.207 | -0.150 | -0.096 |
| $\text{P}_l$ - $\text{P}_u$ | -0.085 | -0.062 | -0.041 |
| $\text{D}_u^2$ - $\text{D}_l^2$ | -0.029 | -0.020 | -0.012 |

Table S8: Average values of statistics across all locations pairs and confidence intervals based on spatial resampling for *Ceratium furca* abundance data.

| | $2.5^{th}$ quantile | Mean | $97.5^{th}$ quantile |
|---|---|---|---|
| Spearman | 0.383 | 0.477 | 0.573 |
| Kendall | 0.272 | 0.342 | 0.418 |
| $\text{cor}_l$ | 0.222 | 0.280 | 0.336 |
| $\text{cor}_u$ | 0.157 | 0.203 | 0.252 |
| $\text{P}_l$ | 0.075 | 0.094 | 0.115 |
| $\text{P}_u$ | 0.054 | 0.067 | 0.082 |
| $\text{D}_l^2$ | 0.026 | 0.035 | 0.045 |
| $\text{D}_u^2$ | 0.038 | 0.046 | 0.054 |
| $\text{cor}_l$ - $\text{cor}_u$ | 0.031 | 0.077 | 0.118 |
| $\text{P}_l$ - $\text{P}_u$ | 0.011 | 0.027 | 0.044 |
| $\text{D}_u^2$ - $\text{D}_l^2$ | 0.003 | 0.011 | 0.019 |

Table S9: Average values of statistics across all locations pairs and confidence intervals based on spatial resampling for methane-flux data.

| | $2.5^{th}$ quantile | Mean | $97.5^{th}$ quantile |
|---|---|---|---|
| Spearman | 0.180 | 0.300 | 0.421 |
| Kendall | 0.126 | 0.211 | 0.300 |
| $\mathrm{cor}_l$ | 0.094 | 0.157 | 0.219 |
| $\mathrm{cor}_u$ | 0.085 | 0.150 | 0.218 |
| $\mathrm{P}_l$ | 0.038 | 0.056 | 0.075 |
| $\mathrm{P}_u$ | 0.034 | 0.050 | 0.071 |
| $\mathrm{D}_l^2$ | 0.044 | 0.055 | 0.066 |
| $\mathrm{D}_u^2$ | 0.046 | 0.057 | 0.067 |
| $\mathrm{cor}_l$ - $\mathrm{cor}_u$ | -0.039 | 0.007 | 0.050 |
| $\mathrm{P}_l$ - $\mathrm{P}_u$ | -0.008 | 0.006 | 0.019 |
| $\mathrm{D}_u^2$ - $\mathrm{D}_l^2$ | -0.007 | 0.002 | 0.010 |
